# Secondary structure determines electron transport in peptides

**DOI:** 10.1101/2024.02.18.578245

**Authors:** Rajarshi Samajdar, Moeen Meigooni, Hao Yang, Jialing Li, Xiaolin Liu, Nicholas E. Jackson, Martín A. Mosquera, Emad Tajkhorshid, Charles M. Schroeder

## Abstract

Proteins play a key role in biological electron transport, but the structure-function relationships governing the electronic properties of peptides are not fully understood. Despite recent progress, understanding the link between peptide conformational flexibility, hierarchical structures, and electron transport pathways has been challenging. Here, we use single-molecule experiments, molecular dynamics (MD) simulations, non-equilibrium Green’s function-density functional theory (NEGF-DFT) calculations, and unsupervised machine learning to understand the role of primary amino acid sequence and secondary structure on charge transport in peptides. Our results reveal a two-state molecular conductance behavior for peptides across several different amino acid sequences. MD simulations and Gaussian mixture modeling are used to show that this two-state molecular conductance behavior arises due to the conformational flexibility of peptide backbones, with a high-conductance state arising due to a more defined secondary structure (beta turn) and a low-conductance state occurring for extended peptide structures. Conformer selection for the peptide structures is rationalized using principal component analysis (PCA) of intramolecular hydrogen bonding distances along peptide backbones. Molecular conformations from MD simulations are used to model charge transport in NEGF-DFT calculations, and the results are in reasonably good agreement with experiments. Projected density of states (PDOS) calculations and molecular orbital visualizations are further used to understand the role of amino acid side chains on transport. Overall, our results show that secondary structure plays a key role in electron transport in peptides, which provides new avenues for understanding the electronic properties of longer peptides or proteins.

**Significance Statement:** Electron transport in proteins serves as a biological power line that fuels cellular activities such as respiration and photosynthesis. Within cells, proteins act as conduits, shuttling electrons through a series of reactions and pathways to generate proton gradients and to fuel ATP synthesis. Despite recent progress, the mechanisms underlying the flow of energy in protein complexes are not fully understood. Here, we study electron transport in peptides at the single-molecule level by combining experiments and molecular modeling. Our results reveal two distinct molecular sub-populations underlying electron transport that arise due to the flexibility of peptide backbones and the ability to fold into compact structures. This work provides a basis for understanding energy flow in larger proteins or biomolecular assemblies.

## Introduction

Electron transport in proteins is essential for maintaining fundamental life processes such as respiration and photosynthesis^1^. In recent years, a wide range of experiments and theoretical studies has focused on understanding electron transfer in biological systems ^2–4^, ranging from redox events in metalloproteins ^5,6^ and redox-active cofactors ^7,8^ to metal-reducing bacteria ^9^. Recent work has shown that proteinaceous nanowire filaments of metal-reducing bacteria such as *Geobacter sulfurreducens* exhibit remarkable abilities for long-distance electron transport on the micron scale^10,11^. During such redox-mediated electron transport events, intervening residues between redox centers are thought to provide a conductive matrix for electron transport ^12^. However, proteins exhibit complex secondary structures due to intramolecular hydrogen (H)-bonding interactions within the underlying conductive protein matrix. Despite recent progress, understanding how secondary structure formation in peptides and proteins affects electron transport is not yet fully understood.

Electron transport in molecules can occur by different mechanisms such as single-step (coherent) tunneling, multi-step (incoherent) hopping, resonant tunneling, or flickering resonant tunneling^13–15^. The dominant mechanism for nanoscale charge transport in short peptide sequences has been reported as non-resonant coherent tunneling^3,4,16–22^, where conductance decays exponentially with molecular length. However, electron transport in long peptide or protein sequences also occurs by hopping^7,23^, where conductance decreases inversely with distance. The environment around a protein affects the driving force for the electron transfer reaction and the reorganization energy, in accordance with Marcus theory^24^. Molecular conformation and intramolecular H-bonding that arise due to the protein sequence and environment are pivotal for controlling biological electron transport over long distances^25^. Prior work has focused on understanding electron transport in helical peptides ^26–29^ using bulk conductivity, electrochemistry, thin-film conductivity, or electronic measurements on assembled peptide monolayers ^7,19,27^. However, key knowledge gaps remain in understanding how other types of secondary structures in peptides and proteins affect electron transport in biological systems. Elucidating electron transport at the single-molecule level holds the potential to provide valuable new insights into the electronic properties of more complex peptide or protein structures.

Single-molecule techniques offer the ability to characterize conformation-dependent electron transport in the absence of intermolecular interactions in monolayers or bulk-scale measurements. In recent years, single-molecule conductance measurements for peptides have primarily focused on short peptide sequences containing up to two or three amino acids^22,30^ or chemically functionalized peptides to facilitate metal electrode contact^31^. However, peptide backbones are generally more flexible compared to π-conjugated carbon backbones commonly used in synthetic organic electronic materials, and this enhanced backbone flexibility could give rise to conformation-dependent electron transport pathways in oligopeptides. By using the scanning tunneling microscope break junction (STM-BJ) technique, the phenomenon of electron tunneling while pulling^32,33^ single molecules has been studied. In addition, it has been reported that a special arrangement of hydrogen bonds^34,35^ could give rise to conducting pathways in non-conjugated peptide backbones. From this view, single-molecule methods offer intriguing routes to understand the role of amino acid sequence and the effect of secondary structure on molecular charge transport in peptides, thereby adding new insights into electronic phenomena and their correlation to structure in biomolecular systems.

The ability to combine molecular simulation with single-molecule electronics experiments provides a powerful approach to understand biophysical processes. The rich conformational space of biomolecules^36^ such as peptides^37^ can be explored using molecular dynamics (MD) simulations. Biomolecular simulation offers a predictive tool for structural biology due to the high spatial and temporal resolutions and the extensively tested and validated force fields^38,39^. MD simulations have been used to understand the influence of atomic structure on the electronic properties of synthetic organic materials by modeling the structural dynamics of molecular junctions ^40–42^. However, classical force fields are limited in their description of molecular junctions that involve transition metal atoms such as gold. Incorporating Au atoms into classical MD simulations requires either a physically rigorous but computationally demanding quantum mechanical (QM) description of gold and its interaction with the surrounding system, or an approximate but more computationally feasible model of interactions with Au atoms. Examples of the latter include representing gold atoms as dummy particles restricted to only interact with specified anchor atoms through harmonic potentials^41^ and utilization of reactive force fields to model bond formation and disruption^43^. Molecular conformations generated by MD can be used in computationally efficient QM calculations for improved comparison between theory and experimental results.

In this work, we investigate the role of amino acid sequence and secondary structure on the electronic properties of peptides using a combination of experiments and computational modeling. A key feature of our work lies in using MD simulations to understand the conformational dynamics of molecular junctions in single-molecule charge transport experiments. A scanning tunneling microscope break junction (STM-BJ) technique is used to experimentally characterize the molecular charge transport properties of oligopeptides. Our results reveal a two-state conductance behavior for peptide sequences containing 4 or 5 amino acids. Our results further indicate that longer amino acid sequences can show enhanced conductance values for the extended state due to the presence of aromatic or constrained amino acid side chains. Gaussian mixture modeling (GMM) and MD simulations are used to show that this two-state molecular conductance behavior arises due to the conformational flexibility of the peptide backbone. Classical MD simulations with custom potentials for implicitly representing gold are used to understand the molecular basis for conformation-dependent electron transport in peptides. Characteristic conformers for each peptide sequence are selected from MD simulations and quantitatively analyzed using principal component analysis (PCA) to understand the role of hydrogen bonding (H-bonding) interactions along the peptide backbone. Interestingly, results from PCA show that specific H-bonding distances between peptide backbone atoms significantly contribute to the structural variation observed in MD simulations. Molecular conformations from MD simulations are then used in non-equilibrium Green’s function-density functional theory (NEGF-DFT) calculations to understand the role of molecular conformation on charge transport. Projected density of states (PDOS) calculations and molecular orbital visualization are further carried out to understand the role of amino acid side chains and the underlying transport mechanisms. Our results reveal that an extended peptide sequence gives rise to a low conductance state, whereas a folded conformation (beta turn) gives rise to a high conductance state. Overall, our work highlights the importance of molecular conformation and secondary structure on the electron transport behavior of peptides.

## Results and Discussion

### Single-molecule conductance measurements and chemical characterization

Tetra- and pentapeptides were designed with different amino acid sequences to understand the role of non-polar aliphatic R groups, aromatic R groups, or sterically constrained R groups on electron transport (**Figures 1a,b,c** and **Supplementary Figures 1-10**). The N- and C-terminal residues of the tetra- and pentapeptides were selected as methionine, which contains a methyl sulfide (-S-CH_3_) group that readily binds to gold ^44^, thereby providing robust electrical contacts to metal electrodes in STM-BJ. All STM-BJ measurements on peptides were carried out in water (peptide concentration <1 mM).

**Figure 1:**
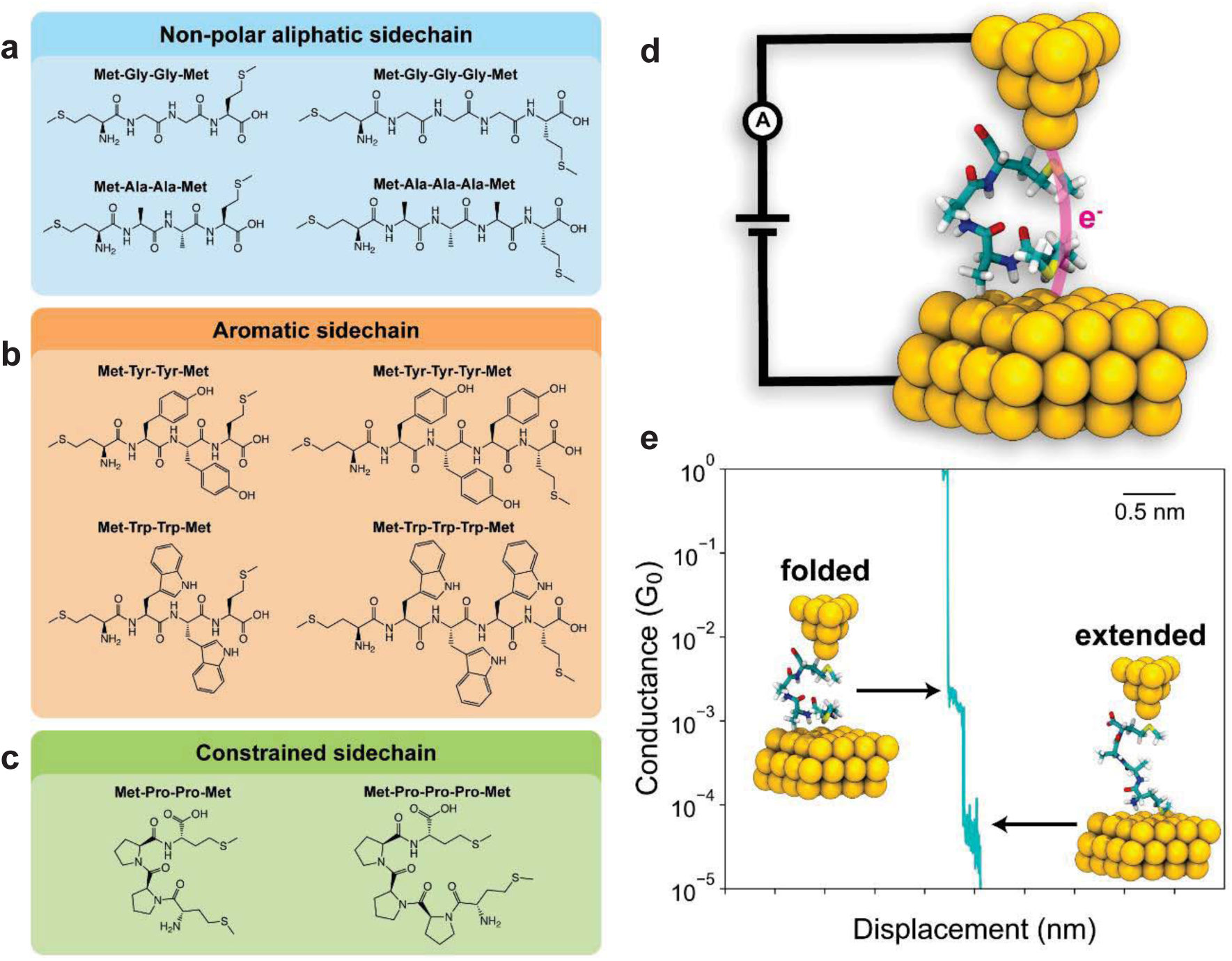
Schematic of experimental setup and chemical structures of peptides studied in this work. Structures of tetra- and pentapeptide with **(a)** nonpolar aliphatic, **(b)** aromatic, or **(c)** sterically constrained R-groups. **(d)** Schematic of a single-molecule junction containing peptide with sequence Met-Ala-Ala-Met (MAAM) using conformations from MD simulations. **(e)** Characteristic single-molecule trace for a peptide with sequence MAAM revealing two distinct conductance populations.

Circular dichroism (CD) spectra were first obtained for all tetra- and pentapeptides in water at room temperature under identical solvent conditions used in STM-BJ experiments (**Supplementary Figures 11-15**). CD spectra clearly indicate the presence of H-bonding interactions for all tetra- and pentapeptides and show spectral features expected for 3_10_ helices, such as a maximum or minimum around ∼200-210 nm and a shoulder or small peak around ∼220 nm ^45,46^. CD spectral features for 3_10_ helices are qualitatively different than the spectral features observed for alpha helices, beta sheets, or random coils ^47^. Based on results from CD experiments, the proline and alanine-based peptide sequences show minima in CD spectra around 200 nm, which is consistent with a tendency to adopt right-handed 3_10_ helices. On the other hand, peptide sequences containing glycine, tyrosine, and tryptophan show peaks in CD spectra around 200 nm, which is consistent with left-handed 3_10_ helices. Overall, these results clearly indicate the presence of H-bonding interactions amongst the tetra- and pentapeptides characterized in single-molecule electronics experiments.

We began by characterizing the electronic properties of peptides containing non-polar aliphatic R groups. The molecular conductance of oligopeptides was determined using a custom-built STM-BJ instrument (**Figure 1d**), as described in prior work^48,49^. Our experiments revealed the presence of two distinct conductance populations, as shown in characteristic single-molecule conductance traces (**Figure 1e**). We hypothesized that the high and low conductance states could arise due to a folded, compact conformation and an extended peptide conformation, respectively. Characteristic single-molecule conductance traces for all tetra- and pentapeptides (**Figures 2a,b**) indicate that the two conductance states occur in the same individual traces rather than in two separate molecular sub-populations. This behavior suggests that dynamic conformational changes during molecular pulling events give rise to multiple conductance states.

**Figure 2:**
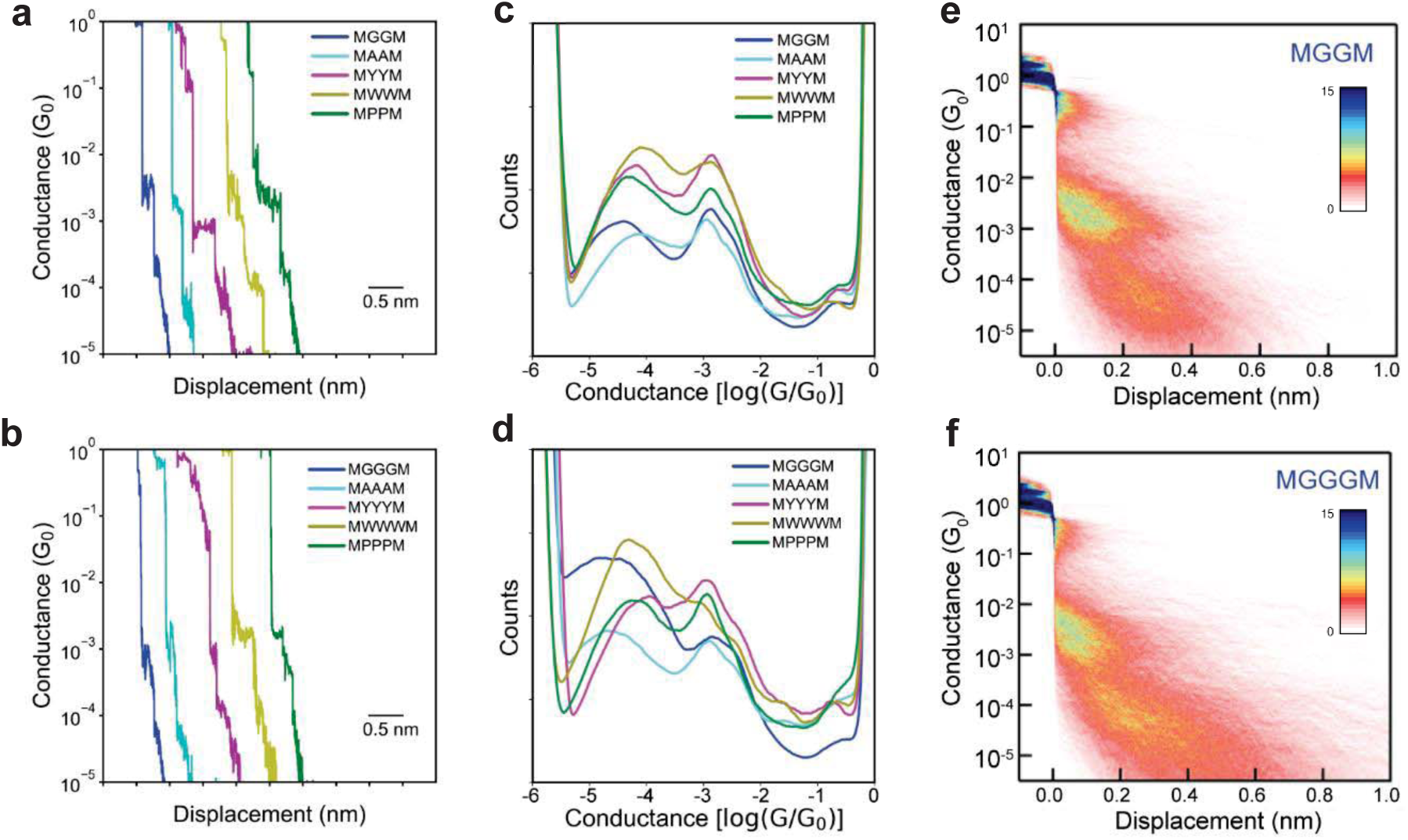
Scanning tunneling microscope-break junction (STM-BJ) measurements of oligopeptides at 250 mV applied bias. **(a)**, **(b)** Characteristic single-molecule traces for tetra- and pentapeptides. **(c), (d)** 1D conductance histograms for the tetra- and pentapeptides. **(e), (f)** 2D conductance histograms for MGGM and MGGGM.

One-dimensional and two-dimensional molecular conductance histograms were generated for the tetra- and pentapeptides across ensembles of >5000 single molecules (**Figures 2c,d,e,f** and **Supplementary Figures 16-17**). A bimodal conductance distribution is observed for all oligopeptide sequences across the entire range of applied biases (100 mV - 400 mV) studied in this work (**Supplementary Figure 18**). Bimodal conductance distributions can arise due to conformationally distinct molecular sub-populations (static heterogeneity) or due to conformation-dependent conductance during molecular pulling (dynamic heterogeneity). To investigate the origins of this behavior, we determined the most probable conductance of the low and high conductance states from a Lorentzian fit to the conductance data^50^ (**Supplementary Tables 1-2**). The high-conductance peak (∼10^-2.80^ - 10^-2.90^ *G_0_*) occurs at nearly the same value for all the oligopeptide sequences. The low conductance peak value shows a small dependence on the backbone sequence and side chain composition. In addition, the molecular displacement corresponding to the low conductance peak is significantly larger than the displacement for the high conductance peak. Based on these results, we hypothesized that the low conductance peak arises due to an extended peptide configuration, whereas the high conductance peak is related to a folded or more compact peptide conformation.

There are some subtle differences in the low conductance state for the tetra- and pentapeptides studied in this work, which suggests that amino acid side chain identity plays a role in transport. For the tetrapeptides, the low conductance state of MGGM is ∼0.2-0.3 log *G_0_* lower compared to all other sequences (MAAM, MYYM, MWWM, and MPPM). These results show that changing the amino acid side chain from hydrogen to a methyl, aromatic, or a constrained side chain leads to an enhancement in conductance. For the pentapeptides, the conductance values for MGGGM and MAAAM are approximately half an order-of-magnitude smaller compared to MYYYM, MWWWM, and MPPPM. The higher conductance values for MYYYM and MWWWM indicate that aromatic side chains can lead to enhanced conductance values. MPPPM has a higher conductance compared to the glycine or alanine-based sequences, as proline provides a constrained side chain that reduces the conformational flexibility and increases the rigidity of the molecule. The higher conductance values observed for the extended conformations for peptides containing tyrosine, tryptophan, and proline sequences are also corroborated by NEGF-DFT simulations (**Figure 5 e,f**), as discussed below. Based on these results, STM-BJ experiments reveal several intriguing findings regarding the role of amino acid side chains on oligopeptide charge transport.

Single-molecule data can be quantitatively analyzed using unsupervised learning algorithms to classify molecular charge transport behavior into characteristic groups and to identify underlying structure-property relationships^31,51–54^. Here, we use silhouette clustering^55^ (**Supplementary Figure 19**) to determine the optimal number of clusters for data sets corresponding to molecular ensembles for each peptide sequence. Silhouette clustering indicates that the optimal number of clusters for all tetra- and pentapeptides is two. Gaussian mixture modeling (GMM) is further used to analyze the two different clusters identified by Silhouette clustering (**Supplementary Figures 20-21**).

Results from GMM show that Cluster 1 accounts for 85-95% of the single-molecule traces and shows both characteristic conductance populations appearing together in the same molecular traces. Cluster 2 accounts for only 5-15% of the data and represents traces in which no molecule is detected or only background signal is observed. If the bimodal distribution arose due to stable, conformationally distinct molecular sub-populations (static heterogeneity), then the two characteristic conductance populations would segregate into different clusters. However, our results show that the bimodal conductance populations appear sequentially in single-molecule traces for all tetra- and pentapeptides, which strongly supports conformation-dependent charge transport behavior in peptide backbones (dynamic heterogeneity).

### MD simulations

To understand the role of molecular conformation on charge transport, we performed MD simulations for all tetra- and pentapeptides (**Figures 1a,b,c**) in explicit solvent with a series of custom potentials to implicitly represent interactions between peptides and gold electrodes. These custom potentials and their resulting collective variable distributions are shown in **Figures 3a,b,c** and **Supplementary Figure 22**.

**Figure 3:**
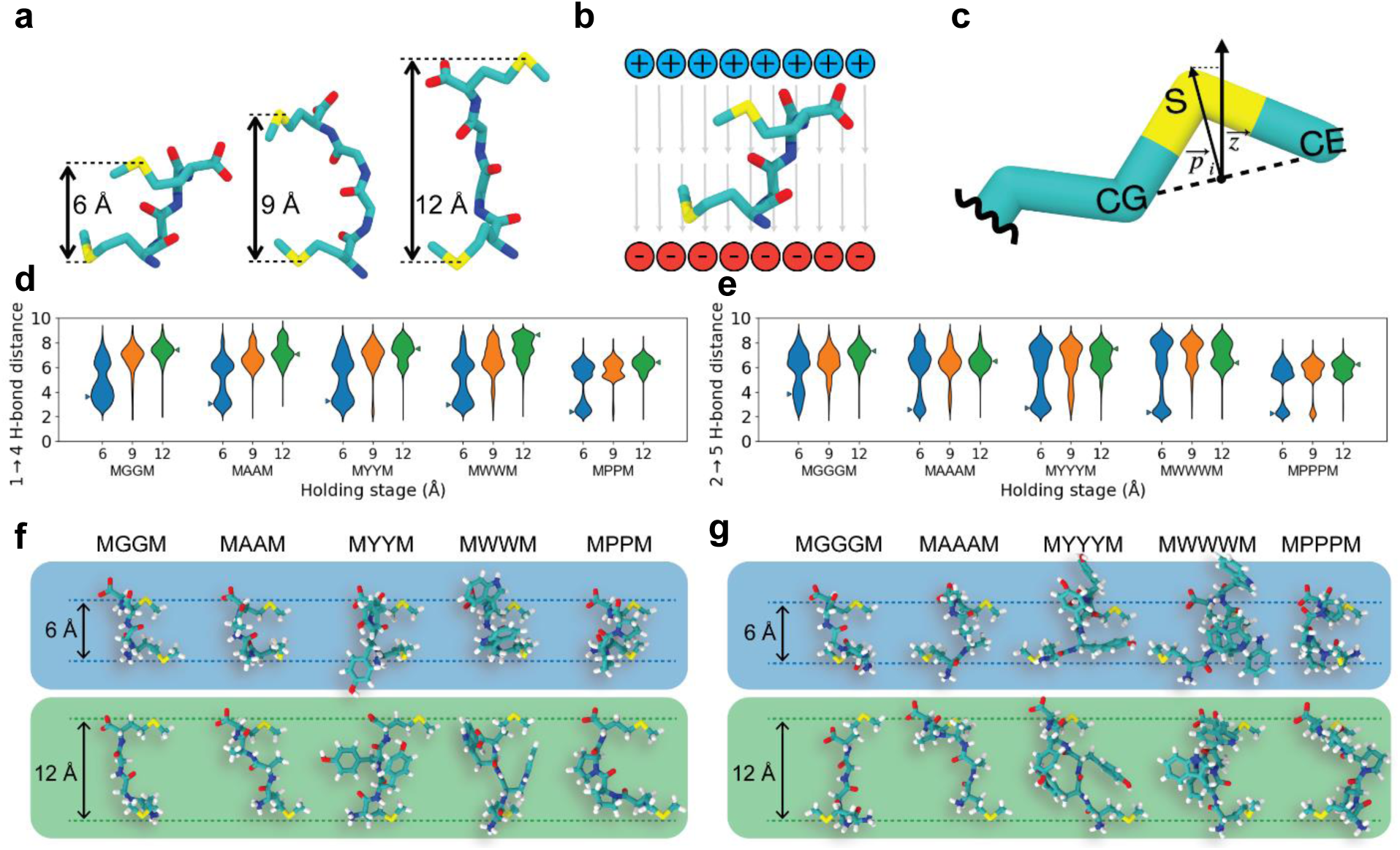
Molecular dynamics (MD) simulations methodology and results. **(a)** Inter-anchor displacement potentials at 6 Å, 9 Å, and 12 Å holding stages defined in Eq. 1. **(b)** Schematic of applied electric field defined in Eq. 2. **(c)** Sulfur-orienting potential defined in Eq. 3. **(d), (e)** Violin plots showing backbone H-bonding distance distribution for tetra- and pentapeptides indicating elimination of intramolecular H-bonds at larger displacements (12 Å). Triangles indicate peaks in the molecular extension distribution at which peptide conformers are selected for NEGF-DFT calculations. **(f), (g)** Snapshots for tetra- and pentapeptide conformations at small (blue, 6 Å) and large (green, 12 Å) displacements.

The projection of the end-to-end distance (sulfur anchor-to-anchor distance on terminal methionines) of the peptide along the experimental pulling axis was harmonically restrained to a range of values (6 Å, 9 Å, and 12 Å), allowing the peptide to adopt an ensemble of conformations. The conformations observed in MD simulations are not significantly affected by a change in applied voltage (**Supplementary Figure 23**), which is consistent with single-molecule conductance experiments. Ramachandran free energy plots^56^ (**Supplementary Figures 24-25**) were determined for the non-terminal residues for all tetra- and pentapeptides. These results indicate that all sequences can form left-handed or right-handed helices, except for those based on proline, consistent with CD measurements.

Results from MD simulations show that backbone hydrogen bonds, which play a key role in defining the secondary structure of the peptide^57^, form with remarkable consistency during the 6 Å end-to-end holding of all peptide sequences considered in this work (**Figures 3d,e** and **Supplementary Figures 26-27**). However, H-bonding interactions are completely abolished when the end-to-end distance is restrained to a distance of 12 Å. For the tetra- and pentapeptides considered here, a canonical secondary structure forms at small end-to-end distances, indicative of a beta turn. A beta turn is defined by an H-bond between the carbonyl oxygen of residue *i* and the amide hydrogen of residue *i+3* ^57^. In the tetrapeptides, a 1→4 H-bond is observed, whereas for the pentapeptides, a 2→5 H-bond is consistently observed. Two conformers are selected from the 6 Å and 12 Å holding stages (**Figures 3f,g**) of each peptide from the peak of the probability distributions of 1→4 H-bond and 2→5 H-bond distances for tetra- and pentapeptides, respectively. It is known that consecutive beta turns in a longer peptide sequence give rise to 3_10_ helices^57^. From this view, our work suggests that helical elements play a key role in the charge transport behavior of biomolecules with defined secondary structures.

We next performed a linear dimensionality reduction on the MD trajectories to quantify how individual interatomic distances contribute to the peptide conformational landscape. The main objective of this analysis is to identify conserved structural differences across all peptides of interest between various end-to-end holding stages (**Figure 4** and **Supplementary Figures 28-31**). Peptides are represented using a Euclidean distance matrix of the common molecular subgraph shared between all sequences. Using this approach, each peptide’s structural ensemble is projected onto a shared basis. In addition, the *i→i+3* H-bonding distances selected as a basis for conformer extraction are well captured in the first two principal components, indicating that these distances contribute significantly to the variance in molecular structure compared to other interatomic distances. Regions of conformational space corresponding to small *i→i+3* distances are shown to depopulate with increasing inter-anchor displacement across all peptide sequences. Based on these results, MD coupled with unsupervised machine learning (ML)-based data analysis clearly elucidates the key structural features for characterizing tetra- and pentapeptides in molecular junctions, revealing the most probable peptide conformations. The most probable conformations are then used in computationally efficient NEGF-DFT calculations to understand the role of molecular conformation on the electron transport properties of peptides.

**Figure 4:**
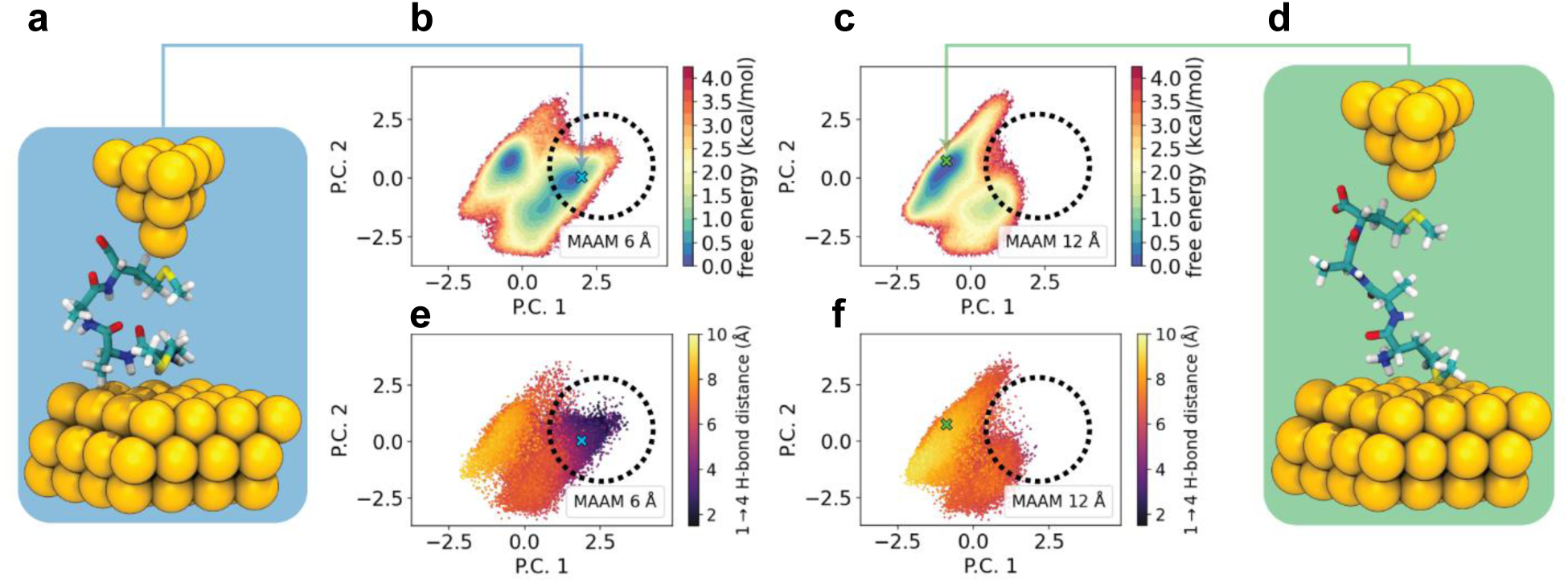
Principal component analysis (PCA) results showing the effect of end-to-end stretching on peptide conformation and molecular descriptors. **(a)** Structure of MAAM turn conformation from MD simulations shown in a single-molecule junction. **(b), (e)** Principal component projections for MAAM turn conformational landscapes denoted with respect to energy and H-bonding distance at a 6 Å holding stage. **(d)** Structure of MAAM extended conformation from MD simulations shown in a single-molecule junction. **(c), (f)** Principal component projections for MAAM extended conformational landscapes colored with respect to energy and H-bonding distance a 12 Å holding stage. Regions of conformational space corresponding to low *i → i+3* backbone H-bond distances are shown to deplete with increasing inter-anchor displacements as denoted by the black dotted circle. Blue and green crosses correspond to MAAM turn and extended conformations, respectively.

### NEGF-DFT calculations

To understand the role of molecular conformation on charge transport in peptide backbones, NEGF-DFT calculations are performed using the most probable simulated MD conformations. NEGF-DFT simulations are carried out for extended and turn conformations for each peptide sequence (**Figures 5a,b**) using the TranSiesta and Tbtrans package (Methods). The transmission probabilities as a function of energy indicate stark differences between the turn and extended peptide conformations (**Figures 5c,d** and **Supplementary Figures 32-34**). The conductance at zero bias differs significantly between the extended and the turn state of the tetra- and pentapeptides (**Figures 5e,f**). Our results show reasonable qualitative agreement between experiments and NEGF-DFT simulations (**Supplementary Table 3-6**). Results from the combined approach of using MD simulations with NEGF-DFT simulations support the hypothesis that the low conductance population arises from an extended peptide conformation, whereas the high conductance population is related to a more defined secondary structure (beta turn) in the peptide. **Figures 5e,f** also corroborate the role of amino acid side chains that was observed in experiments on tetra-and pentapeptides. The glycine-based tetrapeptide sequence has a lower value of the transmission probability near the Fermi level for the extended conformation compared to all other sequences. For the pentapeptides, MGGGM and MAAAM show similar conductance values in the extended state, albeit lower than MYYYM, MWWWM and MPPPM. Overall, NEGF-DFT results qualitatively agree with single-molecule charge transport experiments and provide insights into the role of side chains on oligopeptide charge transport.

**Figure 5:**
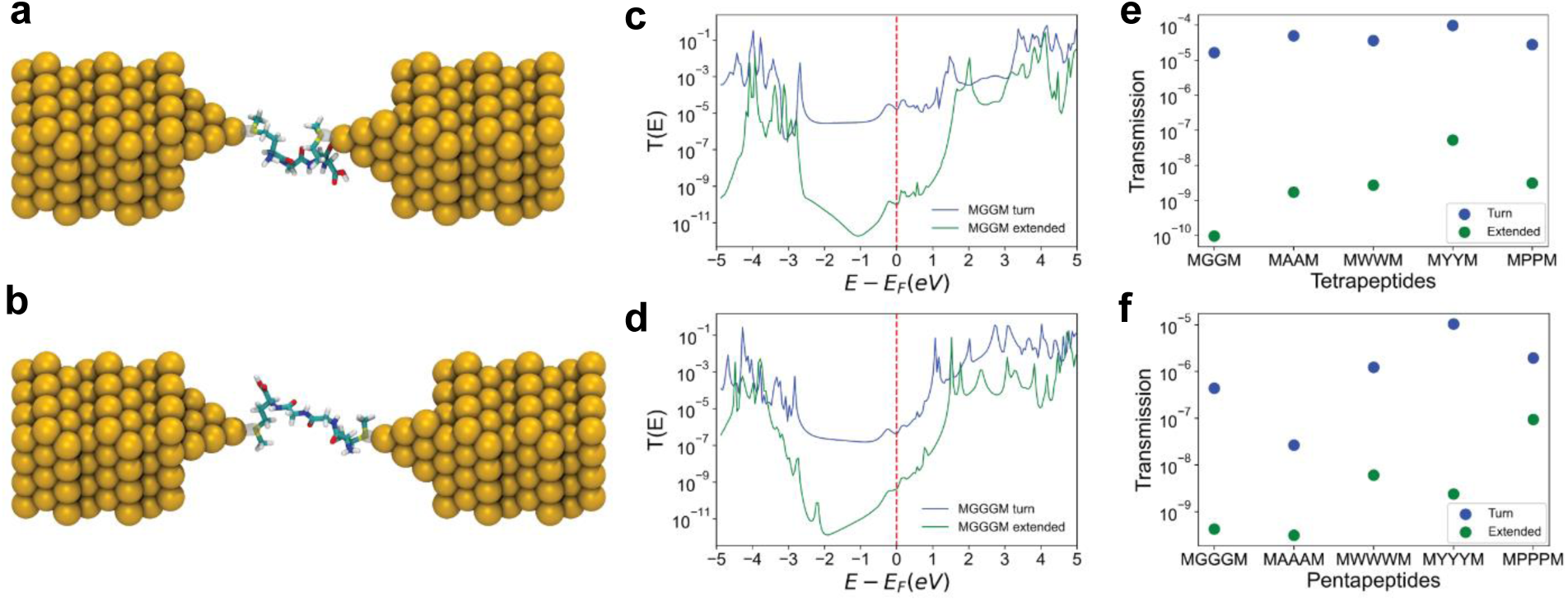
Non-equilibrium Green’s function-density functional theory (NEGF-DFT) calculations for electron transport. **(a), (b)** Schematic of molecular junctions showing gold metal electrodes and MGGM turn and extended conformations for NEGF-DFT calculations. **(c), (d)** Transmission probability as a function of energy (relative to the Fermi energy level) for MGGM and MGGGM, showing drastic differences in transmission probability at *E - E_F_* = 0 for the turn (blue) and the extended (green) configurations. **(e), (f)** Zero bias conductance for tetra- and pentapeptides indicating large differences in transmission probabilities between the two conformational states.

Site specific PDOS calculations were carried out for all tetra-and pentapeptides (**Supplementary Figure 35**) in the extended peptide conformation to understand the role of amino acid side chains on electron transport. For all tetrapeptides, PDOS calculations were performed for two carbon atoms along the backbone in the energy range of -5 to 5 eV (**Supplementary Figures 36 a,b**). At the Fermi energy level for the tetrapeptides (∼ -2.20 eV), the PDOS has a relatively low value (**Table 7**) due to the non-conjugated peptide backbone. These results imply that the backbone orbitals in peptides generally yield smaller conductance values near the Fermi level compared to fully π-conjugated systems. To compare these results for the various tetrapeptides, the behavior near the Fermi energy level was investigated (**Supplementary Figures 36 c,d**). Around the Fermi energy level, higher PDOS values are observed for sequences containing tyrosine and tryptophan. For all pentapeptides, PDOS calculations were also performed for three carbon atoms along the backbone in the energy range of -5 to 5 eV (**Supplementary Figures 37 a,b,c**). The DFT-derived Fermi energy level for the various pentapeptides is around -2.20 eV, with molecular LUMO levels being above the Fermi level by at least 200 meV, though it is important to note that standard DFT generally underestimates the molecular HOMO-LUMO gap. At the Fermi energy level, the value of PDOS approaches zero (**Table 8**), similar to the case of tetrapeptides. A similar analysis was performed for the pentapeptides near the Fermi energy level (**Supplementary Figures 37 d,e,f**), showing larger PDOS values for sequences containing tyrosine and tryptophan for the 1^st^ and 2^nd^ carbon atoms along the backbone. For the 3^rd^ carbon atom, a relatively high PDOS value is observed for MGGGM, MYYYM, and MWWWM. However, the transmission probability for MGGGM is significantly lower compared to MYYYM and MWWWM. Taken together, these results show that the orbitals of the aromatic side chains tend to mix more readily with the backbone orbitals compared to other amino acids, which leads to enhancement in conductance values.

PDOS calculations were also carried out for all carbon and hydrogen atoms (**Supplementary Figure 38**) for MGGM, MYYM, MGGGM, and MYYYM in the extended peptide conformation. Our results indicate significantly higher PDOS values for the sequences containing tyrosine compared to glycine. Overall, these results indicate that oligopeptide sequences with aromatic side chains have more contribution from the backbone orbitals to the overall electronic density and hence molecular conductance.

Molecular orbitals were plotted using Siesta^58^ and visualized using Vesta^59^ (**Supplementary Figures 39-42**). Here, HOMO, HOMO-1, LUMO and LUMO +1 are plotted for the glycine and tyrosine-based tetra- and pentapeptides using an isosurface value of 0.025. These results illustrate relatively weak coupling between the molecules and electrodes, which is consistent with the transmission function results observed for the oligopeptides, in agreement with the proposed tunneling mechanism. These results further suggest the absence of π-π stacking interactions between the tyrosine sidechains. Overall, these results are consistent with non-resonant tunneling rather than resonant tunneling or flickering resonant transport mechanisms for electron transport. Prior work by Xiao et al.^22^ characterized electron transport in short peptide sequences such as cysteamine-glycine-glycine-cysteine and cysteine-glycine-cysteine, with results showing an exponential decay in conductance as a function of molecular length, consistent with single-step tunneling as the dominant transport mechanism. The tetra- and pentapeptides based on glycine studied here are of similar length, with the primary difference of methionine as the N- and C-termini amino acids in place of cysteine or cysteamine. Based on these results, and the relatively short distance of transport observed in our molecular junctions (< 1.4 nm^60^), our results are fully consistent with off-resonant coherent tunneling for the oligopeptides studied in this work.

During the STM-BJ pulling experiments, we observe two conductance states related to two distinct molecular conformations. To further understand the role of H-bonding on transport pathways, we used a bond counting methodology based on the tunneling pathway model^61,62^. In general, there is a conductance decay associated with through-bond, through-space, and through-H-bond electron transport^63^. Tunneling is generally more efficient for through-bond compared to through-space transport due to the lower potential barrier^64^. As a rule of thumb, it can be assumed that the conductance decay through an H-bond is twice as large compared to the decay through a covalent bond^64^. **Supplementary Figure 43** indicates that if transport were to occur entirely through-bond, then the pathway would be approximately three bonds longer with an order of magnitude smaller decay compared to the case of electron transport through H-bonds^65^.

To further understand the importance of H-bonds on transport, we performed control experiments for STM-BJ using 1,16-hexadecanedithiol (**Supplementary Figure 44**) in 1,2,4-tricholorbenzene. 1,16-hexadecanedithiol has a similar contour length as the peptides studied in this work but with a flexible alkane chain backbone and no possibility of intramolecular H-bonding. Our results show that two conductance populations are observed for the peptides (at ∼10^-2.8^ *G*/*G_0_* and 10^-4.2^ *G*/*G_0_*), but no significant conductance peaks are observed for the flexible alkane backbones, though a faint population is observed between ∼10^-1^ -10^-2^ *G*/*G_0_*, which arises due to the use of different anchors and strong binding between the -SH terminal anchor groups and the gold electrode^66^. Overall, these results show a two order-of-magnitude increase in molecular conductance for a peptide compared to an alkane chain with similar contour length. We further compared these results to prior work in the literature. Inkpen et al.^67^ studied charge transport in alkane chains such as C_12_(SH)_2_ and C_12_(SMe)_2_, and only a single conductance population was observed below ∼10^-5^ *G*/*G_0_* for the C_12_ sequences. It should be noted that the average conductance values reported for the C_12_ sequences are approximately one order-of-magnitude lower compared to the low conductance state of the 17- or 19-mer oligopeptide sequences studied in this work. Taken together, these results show that the electron transport behavior of alkane chains is significantly different than peptides due to intramolecular backbone H-bonding.

In this work, we use a combination of single-molecule conductance experiments, MD simulations, and NEGF-DFT calculations to investigate the charge transport properties of a series of different peptide sequences. Our results unequivocally reveal the structure-function relationships governing the observed electron transport in peptides, highlighting the importance of secondary structure on charge transport in biomolecules. Unsupervised learning is used to analyze single-molecule conductance data, showing that peptides exhibit a bimodal conductance distribution with a low and high conductance population arising from distinct conformational states of peptide backbones. A key feature of our work lies in using MD simulations to sample and characterize the conformational space of the peptides and to identify conformations to be used in electron transport (NEGF-DFT) calculations. Moving forward, our work could provide new avenues to understand the interplay between molecular charge transport and secondary structure in more complex peptide sequences with mixed amino acids and/or longer peptides. Proteins are candidate materials for fabricating functional molecular electronic devices due to biocompatibility, anti-fouling properties^68^, and tunable redox activity due to aromatic amino acids^69^. From this view, our work can provide further insights into understanding the role of higher order assembled structures on biological charge transport, which can be used to inform the design self-assembled bioelectronic materials.

## Methods

### Oligopeptide sequences

All oligopeptide sequences were purchased from GenScript (Piscataway, NJ). Mass spectrometry data for these sequences are provided in the Supplementary Information (**Supplementary Figures 1-10**).

### Single-molecule conductance measurements

Single-molecule conductance measurements were performed using a custom-built scanning tunneling microscope break junction (STM-BJ)^48,49,66^. Gold STM tips were prepared using 0.25 mm Au wire (99.998%, Alfa Aesar). STM-BJ experiments were carried out in Milli-Q water (Specific resistance of 18.2 MΩ·cm @ 25 °C). Due to the polarity of the solvent, STM tips were coated with an Apiezon wax to prevent Faradaic currents from masking characteristic molecular features^70^. Gold substrates for the measurements were prepared by evaporating 120 nm of gold on polished AFM metal discs (Ted Pella). Peptide concentrations (<1 mM) were selected to yield Poisson statistics in molecular conductance traces. Conductance histograms (> 5000 traces) are generated for all molecules without data selection. Silhouette clustering and Gaussian mixture modelling (GMM) were further used to analyze the bimodal conductance distribution (Supplementary Information).

### MD simulations

Molecular dynamics (MD) simulations were performed to generate conformational ensembles for the tetra- and pentapeptide molecular junctions at three anchor displacements (referred to as stages 6 Å, 9 Å, and 12 Å). For each peptide, 16 initial structures were prepared using the PeptideBuilder python package^71^. Phi and psi backbone dihedrals of each of the 16 structures were randomized independently. Each backbone dihedral angle of non-proline residues was initialized to a random value between -180 and 180 degrees, whereas the phi angle of proline was initialized to a random value between -80 and -50 degrees. Hydrogens were added to the peptides with the VMD plugin PSFGEN^72^ using the NTER and CTER terminal patches to create positively and negatively charged N- and C-termini, respectively. Peptide structures were then solvated in a cubic box of TIP3P water of side length 38 Å using the VMD SOLVATE plugin^72^. The solvated systems were then subjected to MD simulations with the CHARMM36m protein force field^38,39^ using OpenMM 7.7.0^73^. Dynamics were integrated using the LangevinMiddleIntegrator^74^ with friction coefficient of 1 ps^-1^, temperature of 300 K, and a timestep of 4 fs. Hydrogen mass repartitioning was not utilized. Bonds involving hydrogen atoms, and all bonds and angles involving water were constrained^74^. Nonbonded interactions were computed with a cutoff of 12 Å with smooth switching starting at 10 Å. Electrostatic interactions were evaluated using particle mesh Ewald^75^ (PME) summation with error tolerance of 0.0005. Each replicate was simulated for 200 ns for each of three holding stages, for a total aggregate simulation time of 96.0 μs (10 peptides × 3 stages × 16 replicates × 200 ns). The conformational ensemble of each peptide is shown to converge after 200 ns of simulation per replicate per holding stage (**Supplementary Figures 45-46**). Holding stages were enforced using a series of custom external potentials, applied using OpenMM’s custom force classes, described below. The last 190 ns of each simulation was used for subsequent analysis.

A series of custom potentials were implemented to implicitly represent interactions between the peptide and gold particles. Three potentials were defined: (1) a potential to restrain the distance between the anchors of the molecular junction along the pulling axis to 6 Å, 9 Å, or 12 Å (representing the restraints imposed by connections to the gold electrodes); (2) a per-atom charge-dependent potential along the pulling axis accounting for electric field forces arising from a voltage-biased junction; and, (3) a potential that orients methionine’s thioether moiety such that the average position of each sulfur’s lone pairs are oriented towards the (implicitly represented) gold electrodes along the pulling axis. These potentials are described in detail in the next section and depicted in **Supplementary Figure 17**.

After MD simulations, characteristic conformations of each peptide were determined from their aggregate MD trajectories. For each peptide, two conformations were selected from their 6 Å and 12 Å holding-stage simulations at the peaks of their respective hydrogen-bond distance distribution histograms. The H-bond distance distributions used as the basis for conformation selection for the tetra- and pentapeptides were the 1⟶4 and 2⟶5 distances, respectively. All free energy plots (**Figures 4b,c** and **Supplementary Figures 21,23**) were prepared using PyEMMA 2.5.11^76^.

### Custom potential for implicit gold peptide interactions

A key challenge for simulating single-molecule pulling processes is large difference between the pulling rates used in experiments and those accessible by MD simulations. Typical experimental pulling rates are on the order of Angstroms per millisecond (1 Å per 5 ms in present study), whereas single-trajectory MD simulations (at most) typically reach ms timescales, e.g., with the use of bespoke hardware^77^ or massively distributed computing schemes^78^. In addition, the need for multiple independent simulation replicas to claim ensemble convergence and statistical certainty of key observables further restricts simulations to sub-experimental timescales. However, because the experimental pulling rate is also slow relative to characteristic relaxation timescales of small peptides, we assume that all molecular conformations accessible at a given end-to-end distance are sampled during each step of the experimental pulling process. In other words, experimental pulling occurs as an equilibrium process. Rather than performing costly simulations of the entire pulling process, it is more computationally feasible to simulate the molecular junction at various holding (end-to-end distance) stages representing the different separation distances arising during the pulling experiments.

Using this approach, we performed a series of independent simulations where we restrained the end-to-end (sulfur-sulfur) distance along the pulling axis to one of three distances spanning the range of end-to-end distances (6 Å, 9 Å, or 12 Å). We define the pulling axis as the *z*-axis in our simulations. Schematic illustrations for each potential are shown in **Supplementary Figure 17**. The functional form of the potential utilized to enforce this restraint is given in Equation 1:

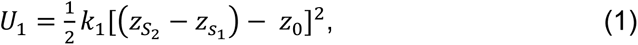

where the coefficient *k_1_* is the force constant of the harmonic potential, *z_S1_* and *z_S2_* are the *z*-coordinate of the sulfur atoms of the N-terminal and C-terminal methionine residues respectively, and *z_0_* is the equilibrium distance for the given stage. We use a value of 1 kcal/mol/Å^2^ for *k_1_*, and we utilize three independent holding stages with *z_0_* equal to either 6 Å, 9 Å, or 12 Å. This force constant was selected such that the resulting distributions of *z_S2_*– *z_S1_* distances have slight overlap (**Supplementary Figures 17a,d**).

By restraining the *z*-displacement between the sulfur atoms, rather than the distance, the movement of each sulfur atom is effectively restrained to one of two parallel planes which implicitly represent two parallel planes of gold electrode.

A potential is introduced to represent an applied electric field due to the voltage difference across the two electrodes. The functional form is given in Equation 2:

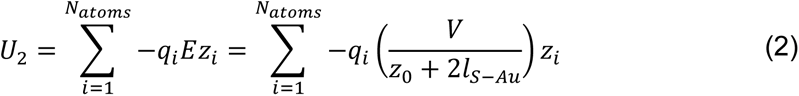

where *N_atoms_*is the total number of atoms in each system including solvent, *q_i_*is the charge of atom *i*, *z_i_* is the z-coordinate of atom *i*, *z_0_* is the equilibrium end-to-end distance (displacement along *z*) for a holding stage, and *l_S-Au_* is the length of the sulfur-gold bond.

We further introduce a potential to orient each sulfur atom’s lone pairs in either the positive or negative *z*-direction, such that a feasible dative bond may occur between the sulfur and a fictitious gold particle. This is a key step in ensuring that any conformation generated by MD simulations can be placed into a gold-gold junction for subsequent NEGF-DFT calculations. Because electron lone pairs are not explicitly represented in atomistic MD simulations, we define surrogate vectors that involve each sulfur’s adjacently bonded carbon atoms to act as a proxy for the direction of the electron lone pairs (**Supplementary Figure 17b**). We impose a restraint directly on the dot product of each surrogate vector with the pulling axis. The functional form of this potential is shown in Equation 3:

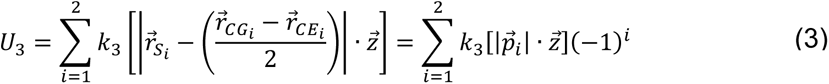

where 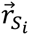 represents the three-dimensional Cartesian coordinates of the sulfur atom of interest, with S_1_ and S_2_ subscripts indicating the identity of the sulfur atoms in the N-terminal and C-terminal methionine residues, respectively, 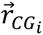 and 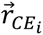 are Cartesian coordinates of the adjacent carbon atoms covalently bonded to each sulfur of interest, and 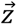 is the unit vector in the direction of the *z*-axis. Vertical lines denote vector normalization. The final term in the equation determines the sign of the potential (and thus the direction of the surrogate vector) allowing for one sulfur’s lone pair to be oriented in the positive *z*-direction while the other is oriented oppositely in the negative *z*-direction. The value of *k_3_* is taken as 10 kcal/mol, resulting in a strong potential that tightly secures the orientation of sulfur lone pairs towards the implicitly represented gold electrodes (**Supplementary Figures 17c,e**).

### Principal component analysis of MD trajectories

The resulting MD trajectory data was subjected to dimensionality reduction by means of principal components analysis (PCA). PCA was performed separately for the tetra- and pentapeptides simulations. For the tetrapeptides, the Cartesian coordinates of the peptide backbone heavy atoms were extracted. The Euclidean distance matrix upper triangle was computed for these 17 shared backbone atoms, resulting in a 136-dimensional vector representation for each trajectory frame. These vector representations, concatenated across all sequences and holding stages and each interatomic distance, were standardized with Z-score normalization. Finally, the first two principal components were calculated with PCA-whitening using the scikit-learn python package^79^. PCA of the pentapeptide trajectories was performed following that of the tetrapeptides, with the exception that the shared molecular subgraph of the pentapeptides was instead composed of 21 backbone heavy atoms, resulting in a 210-dimensional vector representation for each MD trajectory frame. All other steps were performed identically.

### NEGF-DFT calculations

NEGF-DFT calculations are performed with a DFT based non-equilibrium Green’s function (NEGF) approach using the TranSiesta and Tbtrans package^58,80,81^. The electrodes contain 8 layers of 16 gold atoms along with a pyramid of 10 Au atoms. Sulfur atoms in the oligopeptide were made to interact with the gold atoms using a trimer binding motif, as described in literature^30^. Geometry relaxation of the sequences were performed using generalized gradient approximation-Perdew-Burke-Ernzerhof (GGA-PBE) functional^82^ using the TranSiesta package^58^. SZP basis sets were used for all the gold atoms. DZP basis sets were used for carbon, hydrogen, oxygen, sulfur, and nitrogen. Electrode calculations were carried out with a 4 × 4 × 50 k-mesh. The geometry relaxation was carried out using a 4 × 4 × 1 k-mesh, which was performed till all the forces were < 0.05 eV/Å. After the junction was relaxed, the transport calculations were carried out using the TranSiesta package^80,81^ with the same functionals, basis sets, pseudopotential, and k-mesh as the geometry relaxation. Tbtrans^81^ was used to carry out the NEGF calculations and to obtain electron transmission as a function of energy (relative to the fermi energy level). NEGF calculations were carried out from -5 eV to 5 eV with 0.05 eV energy increments. The transmission plots are shifted with respect to the Fermi energy values of each peptide. The difference between charged and uncharged species for the MAAM turn configuration (this is a trial sequence, and not the same sequence obtained from the MD simulations using PCA) has been reported in **Supplementary Fig. 27**. There are similar qualitative agreements between charged (zwitterionic) and uncharged species.

PDOS calculations were carried out for the peptides in the molecular junctions from -5 eV to 5 eV using Siesta^58^. The PDOS calculations were carried out using two Au pyramids and two Au layers, repeated periodically. The PDOS calculations are carried out and plotted over a suitable energy range such that the Fermi energy level of each peptide falls within the interval. For the PDOS calculations, the plane-wave orbitals are projected into atomic orbitals, and the resulting projection coefficients and atomic orbital overlaps that correspond to a given value of the energy in the plot are multiplied together and summed over for each atom of interest. Orbital visualizations were carried out for the molecule with one gold atom on each side using Siesta^58^. The orbitals were visualized using Vesta^59^ to plot HOMO, HOMO-1, LUMO, and LUMO+1 energy levels.

## Supporting information

Samajdar_Supplementary

## Data Availability

Solvent-stripped molecular dynamics trajectories are available at: https://doi.org/10.5281/zenodo.7843691

All other data are available from the corresponding author upon request.

## Code Availability

STM-BJ data were acquired using a custom instrument controlled by custom software (Igor Pro, Wavemetrics). Codes for MD simulations and analysis are available at: github.com/moeenmeigooni/peptide-conductance

## Acknowledgments

This work was supported by the U.S. Department of Energy, Office of Science, Basic Energy Sciences under Award No. DE-SC0022035 for M. Meigooni, H.Y., J.L., E.T., X.L, and C.M.S. and the National Science Foundation under Award 2227399 for R.S. and C.M.S.

## Author contributions

R.S., M. Meigooni, and C.M.S. conceived this study. R.S. performed STM-BJ experiments and NEGF-DFT calculations. M. Meigooni performed MD simulations and PCA analysis. H.Y. assisted with control experiments and J.L. assisted with GMM modeling. X.L assisted with the CD measurements. M. Mosquera and N.E.J. assisted with the NEGF-DFT calculations. E.T. and C.M.S. supervised the research. The manuscript was written by R.S., M. Meigooni, E.T. and C.M.S. with contributions from all authors.

## Additional Information

Supplementary information contains supplementary figures, supplementary tables, and supplementary text.

## References

1. Stillman, M. Biological Inorganic Chemistry. Structure and Reactivity. Edited by Ivano Bertini, Harry B. Gray, Edward I. Stiefel and Joan S. Valentine. Angew. Chem. Intl. Ed. 46, 8741–8742 (2007).

2. Winkler, J. R. & Gray, H. B. Long-range electron tunneling. J. Am. Chem. Soc, 136, 2930–2939 (2014).

3. Amdursky, N. et al. Electronic Transport via Proteins. Adv. Mat. 26, 7142–7161 (2014).

4. Gray, H. B. & Winkler, J. R. Electron tunneling through proteins. Qtly. Rev. of Biophysics 36, 341–372 (2003).

5. Fereiro, J. A. et al. Tunneling explains efficient electron transport via protein junctions. Proc. Natl Acad. Sci. USA 115, 4577–4583 (2018).

6. Nocera, D. G., Winkler, J. R., Yocom, K. M., Bordignon, E. & Gray, H. B. Kinetics of Intramolecular Electron Transfer from Ru11 to FeIH in Ruthenium-Modified Cytochrome c. J. Am. Chem. Soc 106, 5145–5150 (1984).

7. Shipps, C. et al. Intrinsic electronic conductivity of individual atomically resolved amyloid crystals reveals micrometer-long hole hopping via tyrosines. Proc. Natl Acad. Sci. USA 118, e2014139118 (2021)

8. Záliš, Stanislav, et al. Photoinduced hole hopping through tryptophans in proteins. Proc Natl Acad Sci USA 118, e2024627118 (2021)

9. Wang, F. et al. Structure of Microbial Nanowires Reveals Stacked Hemes that Transport Electrons over Micrometers. Cell 177, 361–369 (2019).

10. Ru, X., Zhang, P. & Beratan, D. N. Assessing Possible Mechanisms of Micrometer-Scale Electron Transfer in Heme-Free Geobacter sulfurreducens Pili. J. Phys. Chem. B 123, 5035–5047 (2019).

11. Dahl, P. J. et al. A 300-fold conductivity increase in microbial cytochrome nanowires due to temperature-induced restructuring of hydrogen bonding networks. Sci. Adv. 8, eabm7193 (2022).

12. Williamson, H. R., Dow, B. A. & Davidson, V. L. Mechanisms for control of biological electron transfer reactions. Bioorg. chem. 57, 213–221 (2014).

13. Hines, T. et al. Transition from tunneling to hopping in single molecular junctions by measuring length and temperature dependence. J. Am. Chem. Soc. 132, 11658– 11664 (2010).

14. Zhang, Y. et al. Biological charge transfer via flickering resonance. PNAS 111, 10049–10054 (2014).

15. Li, Songsong, et al. Transition between nonresonant and resonant charge transport in molecular junctions. Nano letters 21, 8340–8347 (2021).

16. Cordes, M. & Giese, B. Electron transfer in peptides and proteins. Chem. Soc. Rev. 38, 892–901 (2009).

17. Juhaniewicz, J., Pawlowski, J. & Sek, S. Electron Transport Mediated by Peptides Immobilized on Surfaces. Israel J. Chem. 55, 645–660 (2015).

18. 16. Scullion, L., et al. Large conductance changes in peptide single molecule junctions controlled by pH. J. Phys. Chem. C 115, 8361–8368 (2011).

19. Baghbanzadeh, M. et al. Charge Tunneling along Short Oligoglycine Chains. Angew. Chem. 127, 14956–14960 (2015).

20. Juhaniewicz, J. & Sek, S. Peptide molecular junctions: Distance dependent electron transmission through oligoprolines. Bioelectrochemistry 87, 21–27 (2012).

21. Guo, Cunlan, et al. Tuning electronic transport via hepta-alanine peptides junction by tryptophan doping. PNAS 113, 10785–10790 (2016).

22. Xiao, Bingqian Xu, and Tao. Conductance titration of single-peptide molecules. JACS 126 5370–5371 (2004).

23. Malak, R. A., Gao, Z., Wishart, J. F. & Isied, S. S. Long-range electron transfer across peptide bridges: The transition from electron superexchange to hopping. J. Am. Chem. Soc. 126, 13888–13889 (2004).

24. Marcus, R. A., Sutin, N. & Amos, A. Electron transfers in chemistry and biology. Biochimica et Biophysica Acta 811, 265–322 (1985).

25. Beratan, David N., J. N. Betts, and J. N. Onuchic. Protein electron transfer rates set by the bridging secondary and tertiary structure. Science 252, 1285–1288 (1991).

26. Amdursky, N. Electron Transfer across Helical Peptides. ChemPlusChem 80, 1075– 1095 (2015).

27. Sepunaru, L. et al. Electronic transport via homopeptides: The role of side chains and secondary structure. J. Am. Chem. Soc. 137, 9617–9626 (2015).

28. Mandal, H. S. & Kraatz, H. B. Electron transfer mechanism in helical peptides. J. Phys. Chem. Lett. 3, 709–713 (2012).

29. Horsley, J. R., Yu, J., Moore, K. E., Shapter, J. G. & Abell, A. D. Unraveling the interplay of backbone rigidity and electron rich side-chains on electron transfer in peptides: The realization of tunable molecular wires. J. Am. Chem. Soc. 136, 12479–12488 (2014).

30. Brisendine, J. M. et al. Probing Charge Transport through Peptide Bonds. J. Phys. Chem. Lett. 9, 763–767 (2018).

31. Stefani, D. et al. Conformation-dependent charge transport through short peptides. Nanoscale 13, 3002–3009 (2021).

32. Lin, Jianping, and David N. Beratan. Tunneling while pulling: the dependence of tunneling current on end-to-end distance in a flexible molecule. J. Phys. Chem A 108, 5655–5661 (2004).

33. Schneebeli, Severin T., et al. Single-molecule conductance through multiple π− π-stacked benzene rings determined with direct electrode-to-benzene ring connections. JACS 133, 2136–2139 (2011).

34. Zhang, Bintian, et al. Role of contacts in long-range protein conductance. PNAS 116, 5886–5891 (2019).

35. Kretchmer, Joshua S., et al. Fluctuating hydrogen-bond networks govern anomalous electron transfer kinetics in a blue copper protein. PNAS 115, 6129–6134 (2018).

36. Ciudad, S. et al. Aβ(1-42) tetramer and octamer structures reveal edge conductivity pores as a mechanism for membrane damage. Nat. Commun. 11, 3014–3028 (2020).

37. Biswas, M., Lickert, B. & Stock, G. Metadynamics Enhanced Markov Modeling of Protein Dynamics. J. Phys. Chem. B 122, 5508–5514 (2018).

38. Brooks, B. R. et al. CHARMM: The biomolecular simulation program. J. Comput. Chem. 30, 1545–1614 (2009).

39. Huang, J. et al. CHARMM36m: An improved force field for folded and intrinsically disordered proteins. Nat. Methods 14, 71–73 (2016).

40. Wang, H. & Leng, Y. Gold/Benzenedithiolate/Gold Molecular Junction: A Driven Dynamics Simulation on Structural Evolution and Breaking Force under Pulling. J. Phys. Chem. C 119, 15216–15223 (2015).

41. Mejía, L., Renaud, N. & Franco, I. Signatures of Conformational Dynamics and Electrode-Molecule Interactions in the Conductance Profile during Pulling of Single-Molecule Junctions. J. Phys. Chem. Lett. 9, 745–750 (2018).

42. Strange, M., Lopez-Acevedo, O. & Häkkinen, H. Oligomeric gold-thiolate units define the properties of the molecular junction between gold and benzene dithiols. J. Phys. Chem. Lett. 1, 1528–1532 (2010).

43. Li, Z. & Franco, I. Molecular Electronics: Toward the Atomistic Modeling of Conductance Histograms. J. Phys. Chem. C 123, 9693–9701 (2019).

44. Batra, A. et al. Tuning rectification in single-molecular diodes. Nano Lett. 13, 6233– 6237 (2013).

45. Kumar, P., Paterson, N. G., Clayden, J., & Woolfson, D. N., De novo design of discrete, stable 310-helix peptide assemblies. Nature, 607(7918), 387-392 (2022).

46. Brown, R. A., Marcelli, T., De Poli, M., Solà, J., & Clayden, J. (2012). Induction of unexpected left-handed helicity by an N-terminal L-amino acid in an otherwise achiral peptide chain. Angewandte Chemie, 124(6), 1424–1428 (2012).

47. Wei, Y., Thyparambil, A. A., & Latour, R. A. Protein helical structure determination using CD spectroscopy for solutions with strong background absorbance from 190 to 230 nm. (BBA)-Proteins and Proteomics, 1844(12), 2331–2337 (2014).

48. Li, S. et al. Charge Transport and Quantum Interference Effects in Oxazole-Terminated Conjugated Oligomers. J. Am. Chem. Soc. 141, 16079–16084 (2019).

49. Li, B. et al. Intrachain Charge Transport through Conjugated Donor-Acceptor Oligomers. ACS Appl. Electron. Mater. 1, 7–12 (2019).

50. Hybertsen, M. S. et al. Amine-linked single-molecule circuits: Systematic trends across molecular families. J. of Phys. Condens. Matter. 20, 374115 (2008).

51. Bamberger, N. D., Ivie, J. A., Parida, K. N., McGrath, D. v. & Monti, O. L. A. Unsupervised Segmentation-Based Machine Learning as an Advanced Analysis Tool for Single Molecule Break Junction Data. J. Phys. Chem. C 124, 18302–18315 (2020).

52. Lin, L. et al. Spectral clustering to analyze the hidden events in single-molecule break junctions. J. Phys. Chem. C 125, 3623–3630 (2021).

53. Liu, B., Murayama, S., Komoto, Y., Tsutsui, M. & Taniguchi, M. Dissecting Time-Evolved Conductance Behavior of Single Molecule Junctions by Nonparametric Machine Learning. J. Phys. Chem. Lett. 11, 6567–6572 (2020).

54. Cabosart, D. et al. A reference-free clustering method for the analysis of molecular break-junction measurements. Appl. Phys. Lett. 114, 143102 (2019).

55. Rousseeuw, P. J. Silhouettes: a graphical aid to the interpretation and validation of cluster analysis. J. Compu. and App. Math. 20, 53–65 (1987).

56. Hollingsworth, Scott A. and Karplus, P. Andrew. “A fresh look at the Ramachandran plot and the occurrence of standard structures in proteins” Biomolecular Concepts 1, 271–283 (2010)

57. Kabsch, W. & Sander, C. Dictionary of protein secondary structure: Pattern recognition of hydrogen-bonded and geometrical features. Biopolymers 22, 2577– 2637 (1983).

58. Soler, J. M. et al. The SIESTA method for ab initio order-N materials simulation. J. Phys.: Condens. Matter. 14, 2745–2781 (2002).

59. Momma, K. & Izumi, F. VESTA 3 for three-dimensional visualization of crystal, volumetric and morphology data. J Appl Crystallogr 44, 1272–1276 (2011).

60. Page, Christopher C., et al. Natural engineering principles of electron tunnelling in biological oxidation–reduction. Nature 402, 47–52 (1999).

61. Beratan, D. N., Onuchic, J. N. & Hopfield, J. J. Electron tunneling through covalent and noncovalent pathways in proteins. J Chem. Phys. 86, 4488–4498 (1987).

62. Wuttke, Deborah S., et al. Electron-tunneling pathways in cytochrome c. Science 256, 1007–1009 (1992).

63. Onuchic, J. N. & Beratan, D. N. A predictive theoretical model for electron tunneling pathways in proteins. J Chem. Phys. 92, 722–733 (1990).

64. Beratan, D. N., Onuchic, J. N., Winkler, J. R. & Gray, H. B. Electron-Tunneling Pathways in Proteins. Science 258, 1740–1741 (1992).

65. Betts, J. N., Beratan, D. N., & Onuchic, J. N. Mapping electron tunneling pathways: an algorithm that finds the“ minimum length”/maximum coupling pathway between electron donors and acceptors in proteins. JACS, 114(11), 4043–4046 (1992).

66. Inkpen, M. S. et al. Non-chemisorbed gold–sulfur binding prevails in self-assembled monolayers. Nat. Chem. 11, 351–358 (2019).

67. Venkataraman, Latha, et al. Single-molecule circuits with well-defined molecular conductance. Nano letters 6, 458–462 (2006).

68. Hostert, J. D. et al. Self-Assembly and Rearrangement of a Polyproline II Helix Peptide on Gold. Langmuir 37, 6115–6122 (2021).

69. Creasey, R. C. G. et al. Biomimetic Peptide Nanowires Designed for Conductivity. ACS Omega 4, 1748–1756 (2019).

70. Nagahara, L. A., Thundat, T. & Lindsay, S. M. Preparation and characterization of STM tips for electrochemical studies. Rev. Sci. Instr. 60, 3128–3130 (1989).

71. Tien, M. Z., Sydykova, D. K., Meyer, A. G., & Wilke, C. O. PeptideBuilder: A simple Python library to generate model peptides. PeerJ 1, e80 (2013).

72. Humphrey, W., Dalke, A., & Schulten, K. VMD: visual molecular dynamics. J. mol. graphics 14, 33–38 (1996).

73. Eastman, P. et al. OpenMM 7: Rapid development of high performance algorithms for molecular dynamics. PLoS Comput. Biol. 13, e1005659 (2017).

74. Zhang, Z., Liu, X., Yan, K., Tuckerman, M. E. & Liu, J. Unified Efficient Thermostat Scheme for the Canonical Ensemble with Holonomic or Isokinetic Constraints via Molecular Dynamics. J. Phys. Chem. A 123, 6056–6079 (2019).

75. Darden, T., York, D. & Pedersen, L. Particle mesh Ewald: An N·log(N) method for Ewald sums in large systems. J. Chem. Phys. 98, 10089–10092 (1993).

76. Scherer, M. K. et al. PyEMMA 2: A Software Package for Estimation, Validation, and Analysis of Markov Models. J. Chem. Theory Comput. 11, 5525–5542 (2015).

77. David shaw, by E., et al. Anton, a Special-Purpose Machine for Molecular Dynamics Simulation. Commun. of the ACM 51, 91-97 (2008).

78. Shirts, M. & Pande, V. S. Screen savers of the world unite. Science 290 1903–1904 (2000).

79. Pedregosa, Fabian, et al. “Scikit-learn: Machine learning in Python. The J. MLR 12, 2825–2830 (2011)

80. Brandbyge, M., Mozos, J. L., Ordejón, P., Taylor, J. & Stokbro, K. Density-functional method for nonequilibrium electron transport. Phys. Rev. B 65, 1654011–16540117 (2002).

81. Papior, N., Lorente, N., Frederiksen, T., García, A. & Brandbyge, M. Improvements on non-equilibrium and transport Green function techniques: The next-generation TRANSIESTA. Comput. Phys. Commun. 212, 8–24 (2017).

82. Perdew, J. P., Burke, K. & Ernzerhof, M. Generalized Gradient Approximation Made Simple. Phys. Rev. lett. 77, 3865–3869 (1996).

