## Supplementary material for "Secondary structure determines electron transport in peptides": Samajdar_Supplementary

**Table of Contents:**

S1. Mass spec data …………………………………………………………………. 3

S2. Circular dichroism (CD) data ………………………………………………….. 13

S3. Single-molecule charge transport experiments ……………………….……….. 18

S4. Gaussian mixture modeling (GMM) …………………………………………….. 22

S5. Molecular dynamics (MD) simulations ………………………………………….. 26

S11. Tunneling pathway model: Bond counting for pathway determination ……. 45

S12. Control experiment: 1,16- hexadecanedithiol ……………………………….. 46

S13. Convergence plots for MD simulations ……………………………………….. 47

**S1. Mass spectrometry data**

Mass spectrometry data for all oligopeptide sequences are reported in this section. The data were obtained by the commercial vendor (GenScript).

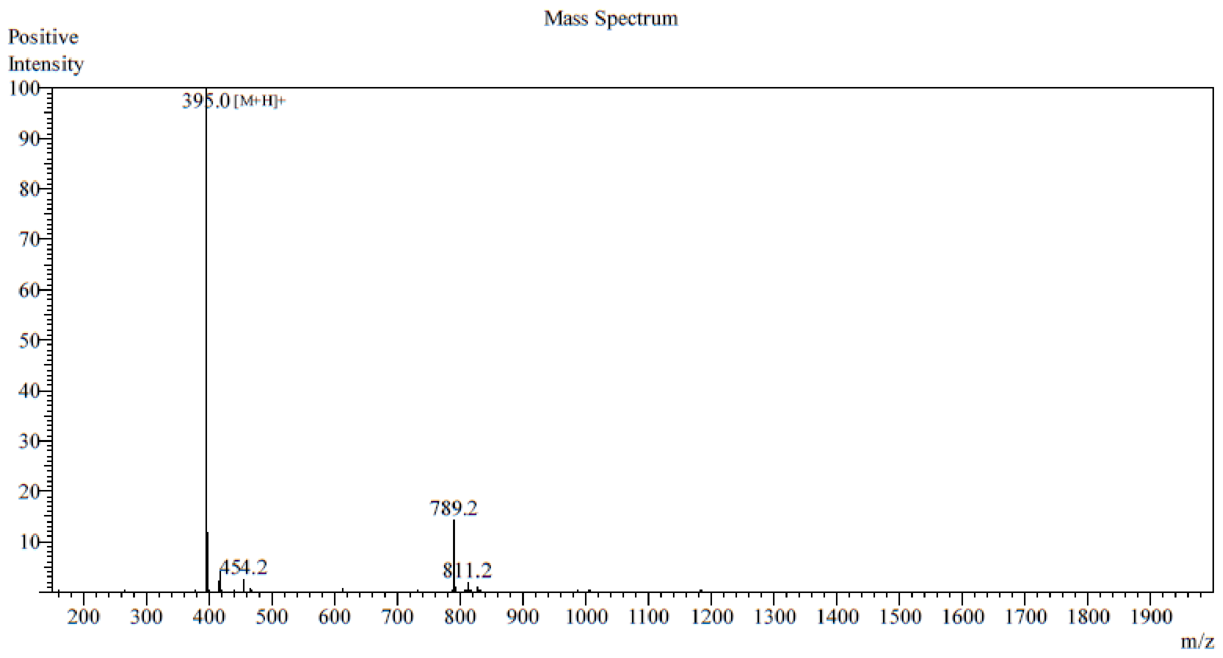

**Supplementary Figure 1**: Electrospray ionization (ESI) mass spectrometry data for peptide sample MGGM. Theoretical molecular weight is 394.51 m/z. Observed molecular weight is 394.0 m/z.

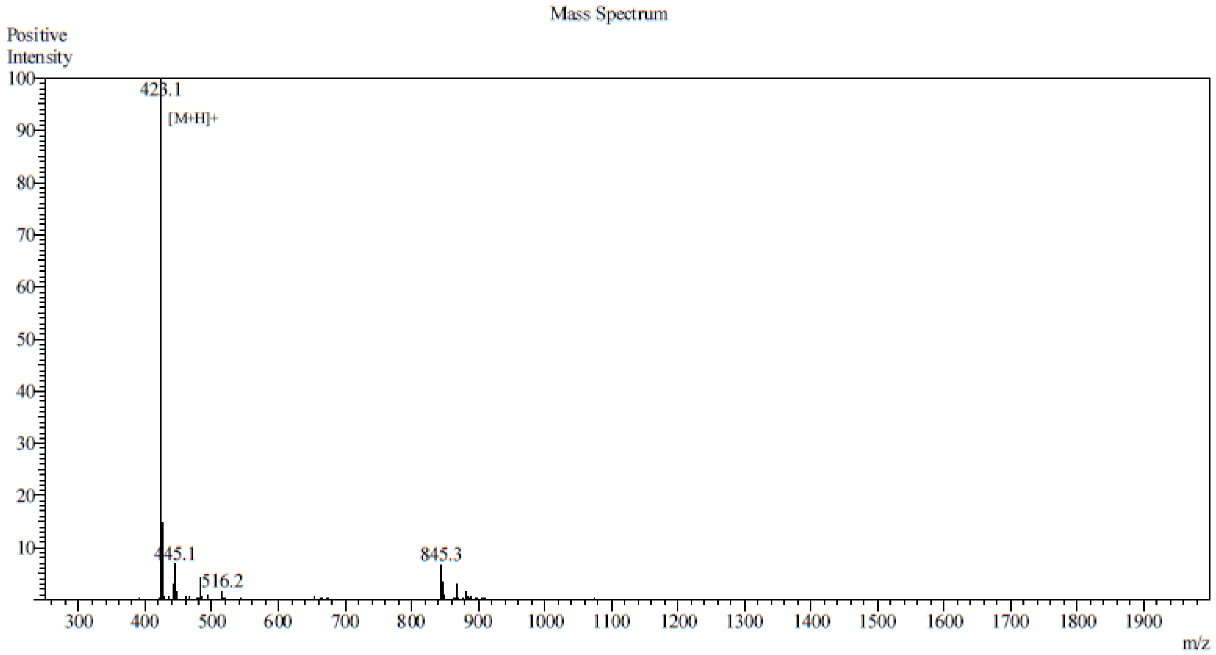

**Supplementary Figure 2**: Electrospray ionization (ESI) mass spectrometry data for peptide sample MAAM. Theoretical molecular weight is 422.57 m/z. Observed molecular weight is 422.1 m/z.

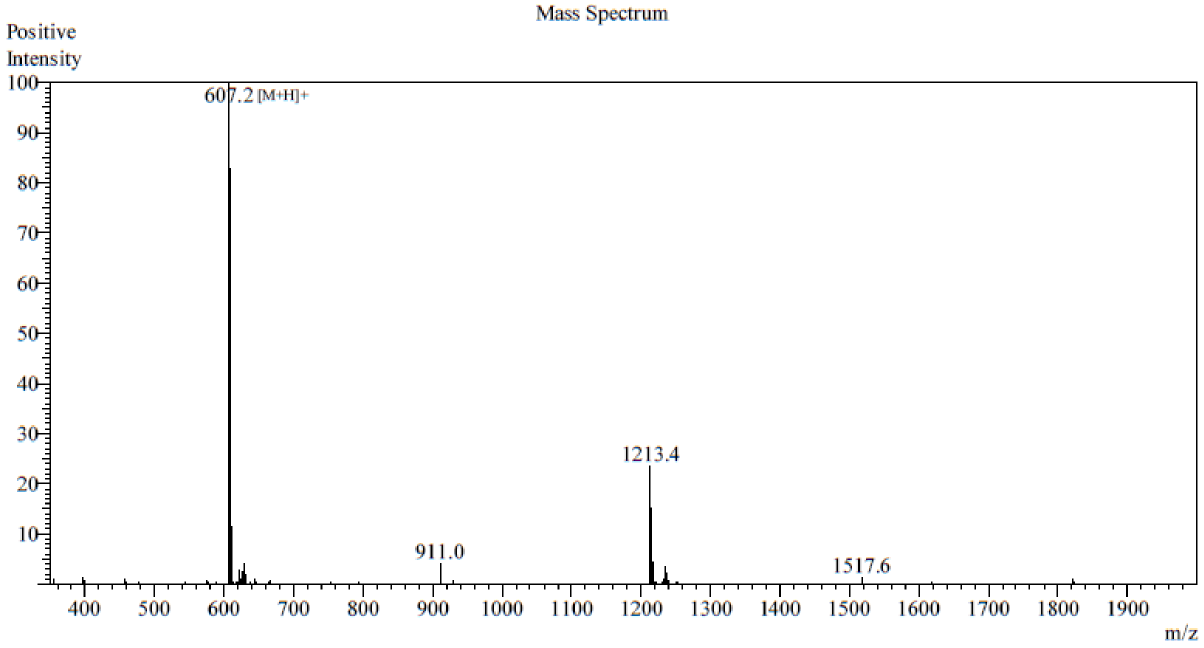

**Supplementary Figure 3**: Electrospray ionization (ESI) mass spectrometry data for peptide sample MYYM. Theoretical molecular weight is 606.76 m/z. Observed molecular weight is 606.2 m/z.

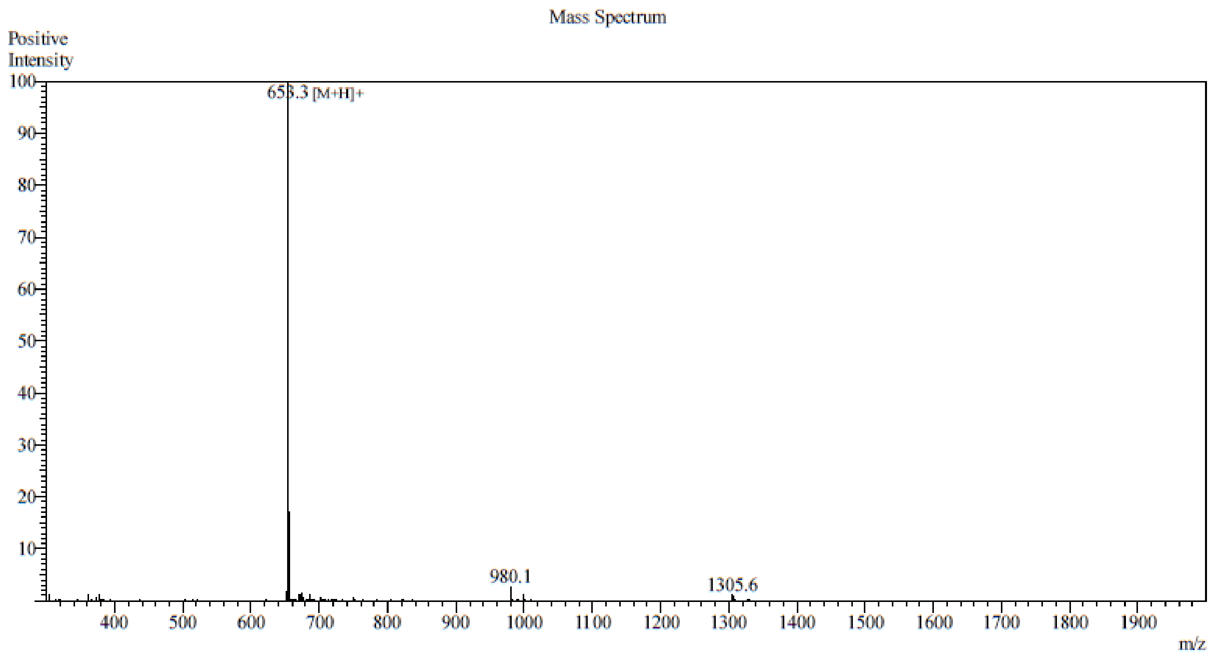

**Supplementary Figure 4**: Electrospray ionization (ESI) mass spectrometry data for peptide sample MWWM. Theoretical molecular weight is 652.83 m/z. Observed molecular weight is 652.3 m/z.

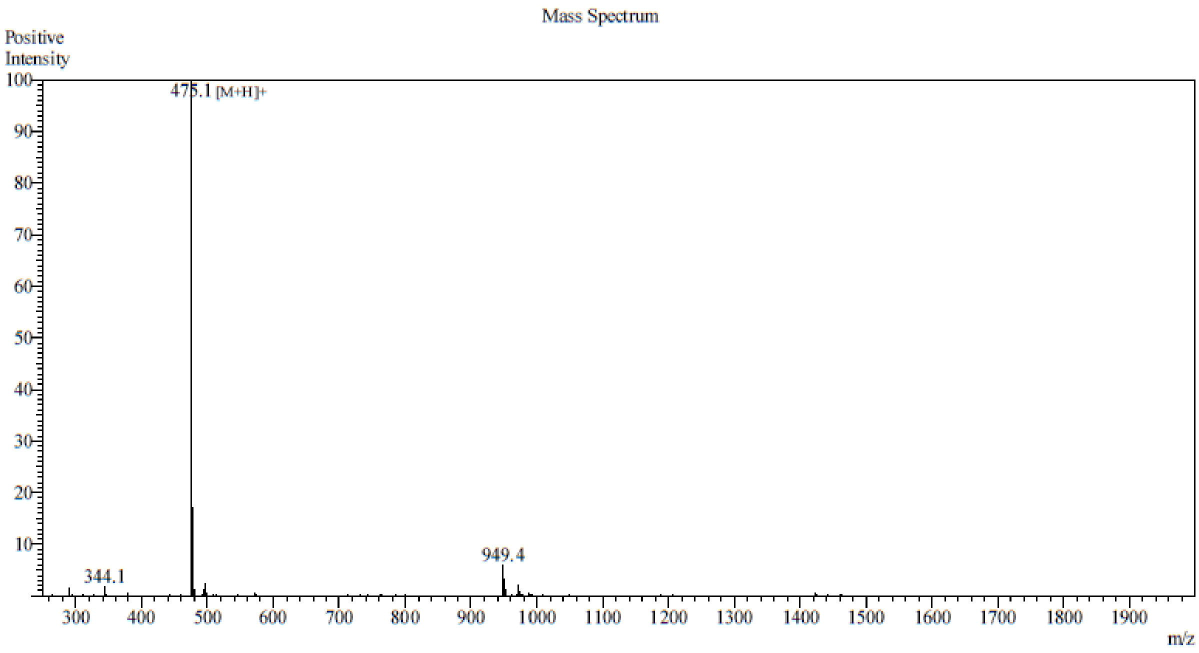

**Supplementary Figure 5**: Electrospray ionization (ESI) mass spectrometry data for peptide sample MPPM. Theoretical molecular weight is 474.64 m/z. Observed molecular weight is 474.1 m/z.

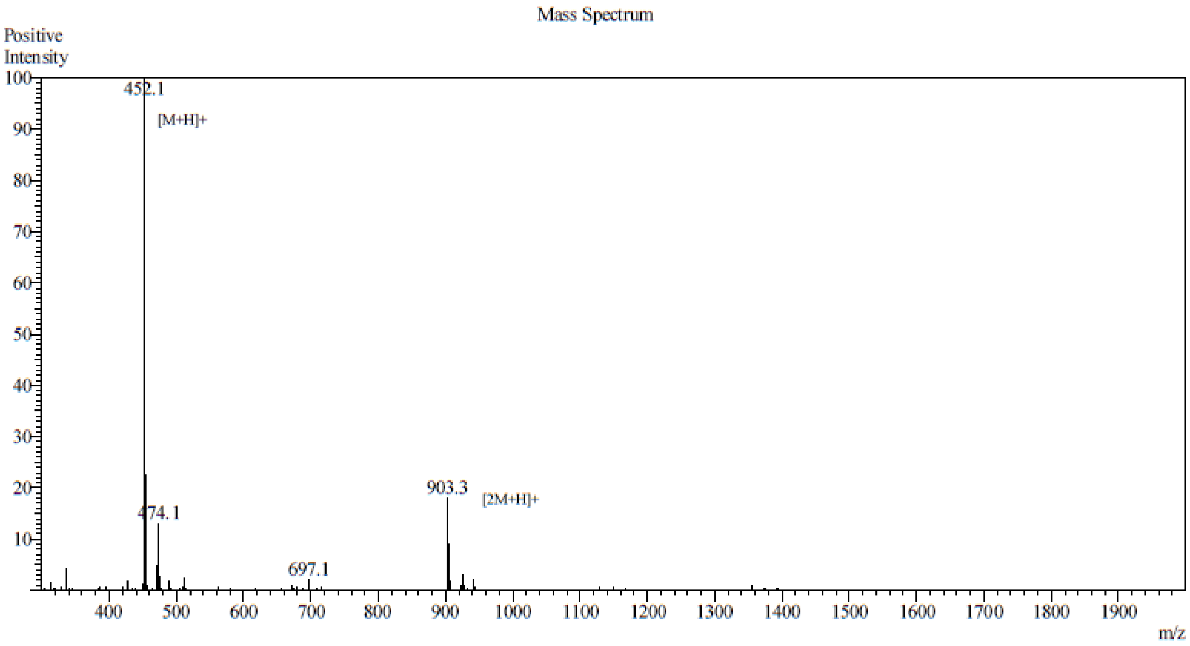

**Supplementary Figure 6**: Electrospray ionization (ESI) mass spectrometry data for peptide sample MGGGM. Theoretical molecular weight is 455.16 m/z. Observed molecular weight is 455.1 m/z.

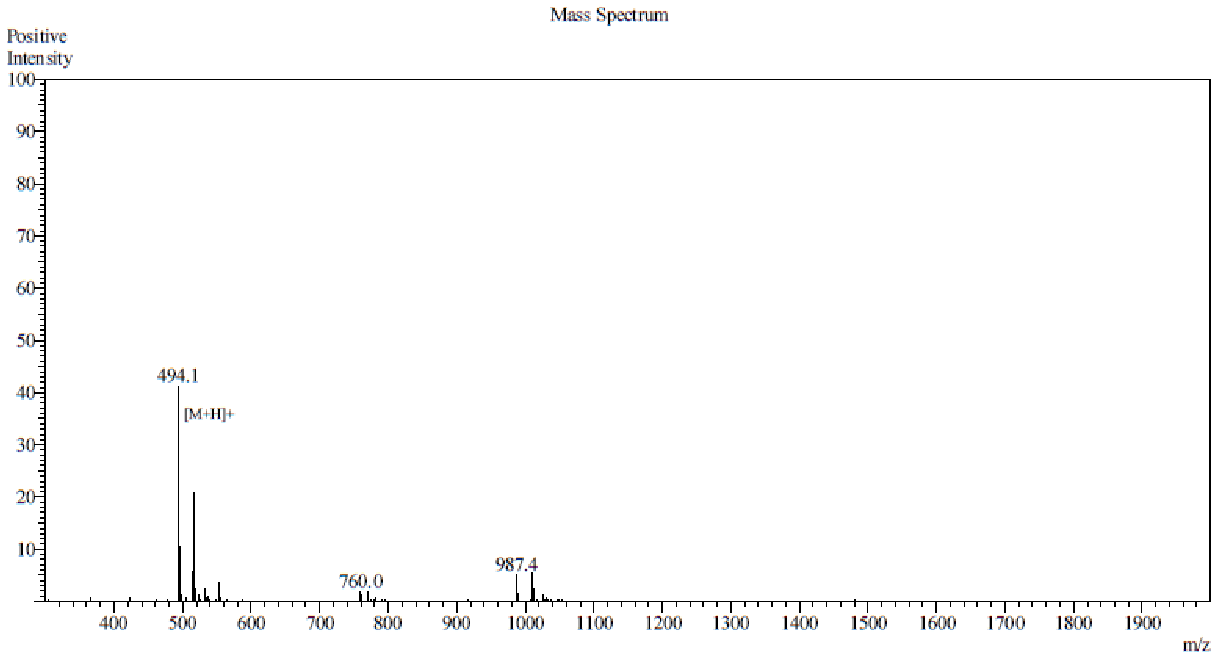

**Supplementary Figure 7**: Electrospray ionization (ESI) mass spectrometry data for peptide sample MAAAM. Theoretical molecular weight is 493.64 m/z. Observed molecular weight is 493.1 m/z.

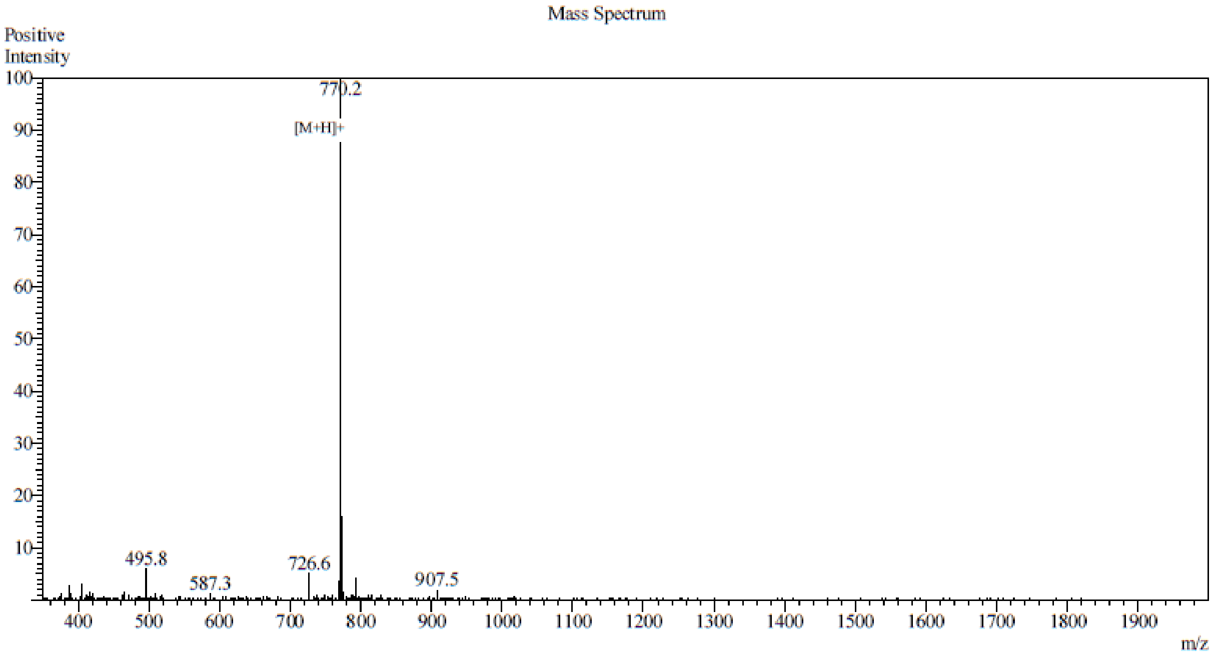

**Supplementary Figure 8**: Electrospray ionization (ESI) mass spectrometry data for peptide sample MYYYM. Theoretical molecular weight is 769.93 m/z. Observed molecular weight is 769.2 m/z.

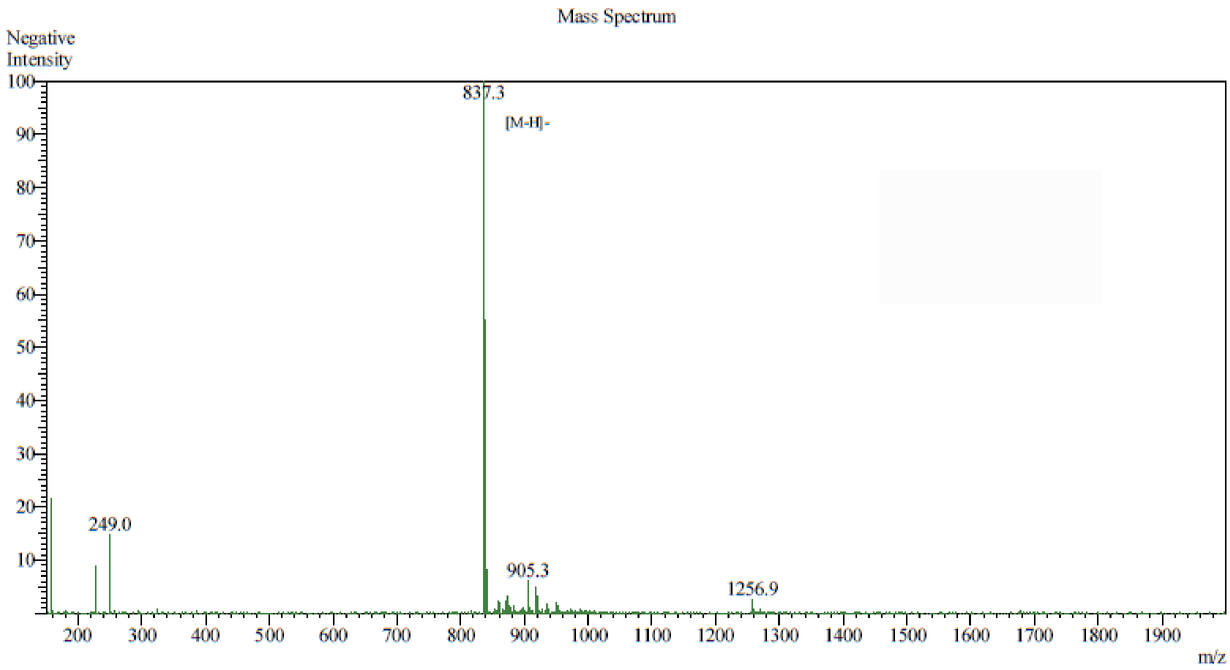

**Supplementary Figure 9:** Electrospray ionization (ESI) mass spectrometry data for peptide sample MWWWM. Theoretical molecular weight is 839.04 m/z. Observed molecular weight is 838.3 m/z.

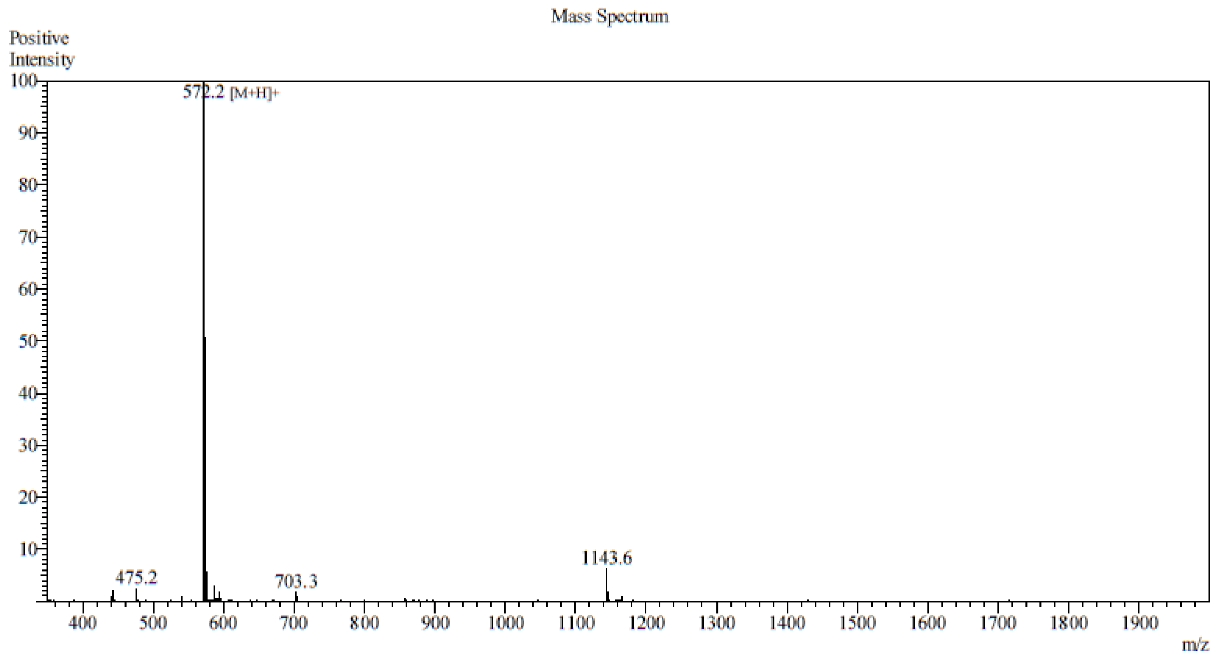

**Supplementary Figure 10**: Electrospray ionization (ESI) mass spectrometry data for peptide sample MPPPM. Theoretical molecular weight is 571.76 m/z. Observed molecular weight is 571.2 m/z.

**S2. Circular dichroism (CD) data**

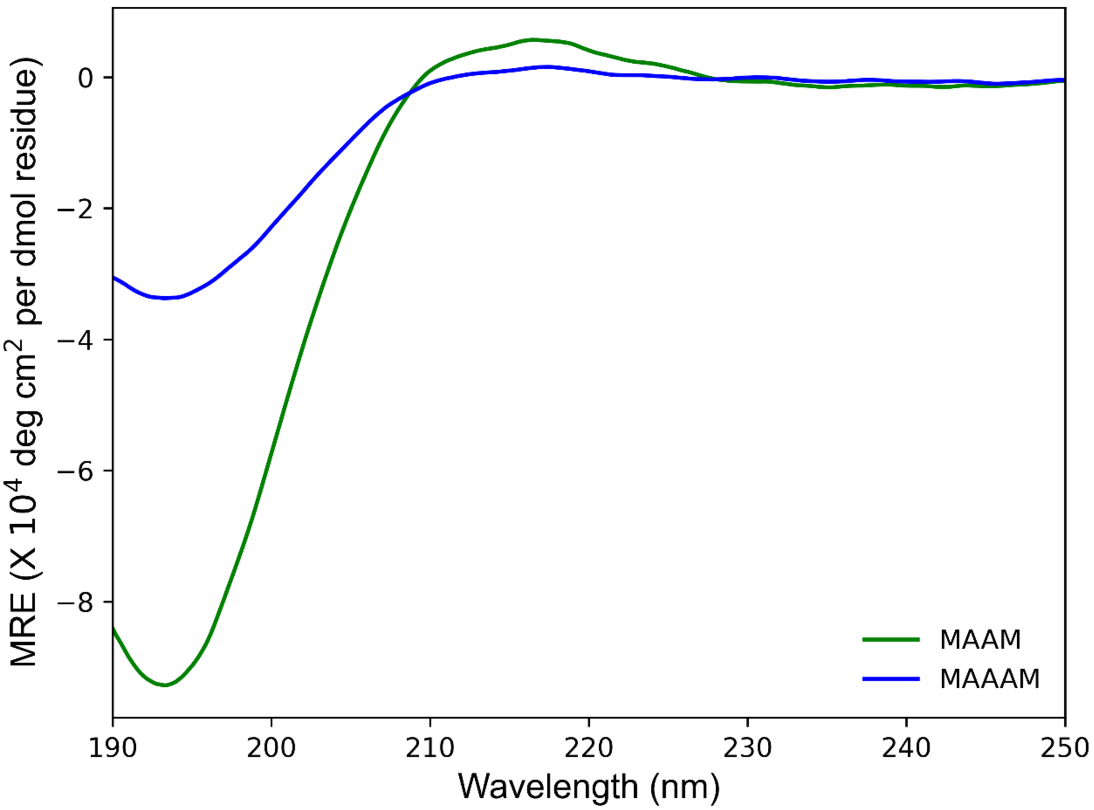

**Supplementary Figure 11**: Circular dichroism (CD) spectra for MAAM and MAAAM peptide sequences.

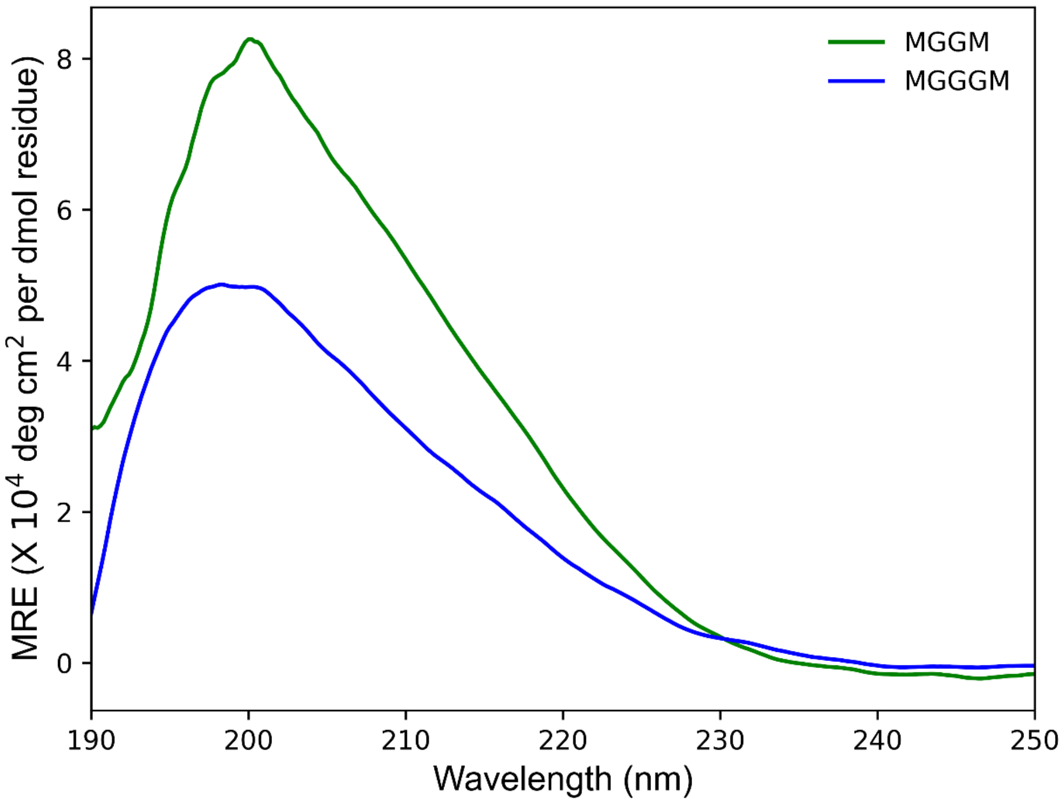

**Supplementary Figure 12**: Circular dichroism (CD) spectra for MGGM and MGGGM peptide sequences.

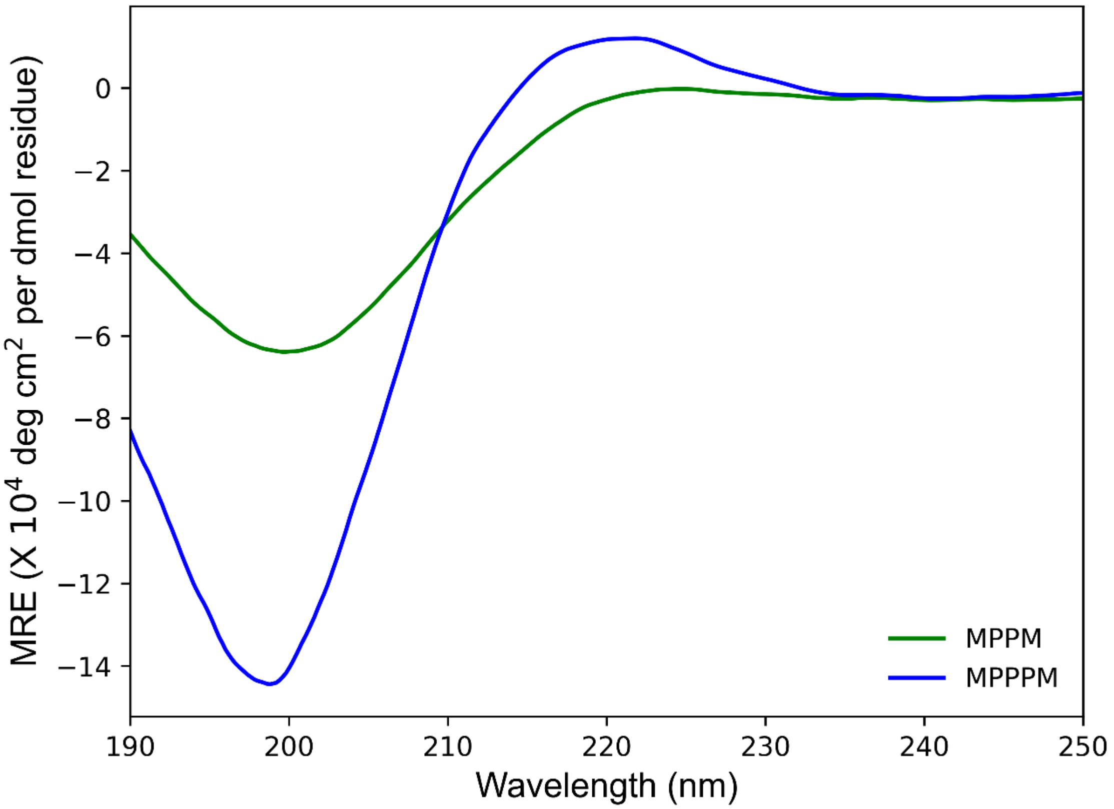

**Supplementary Figure 13**: Circular dichroism (CD) spectra for MPPM and MPPPM peptide sequences.

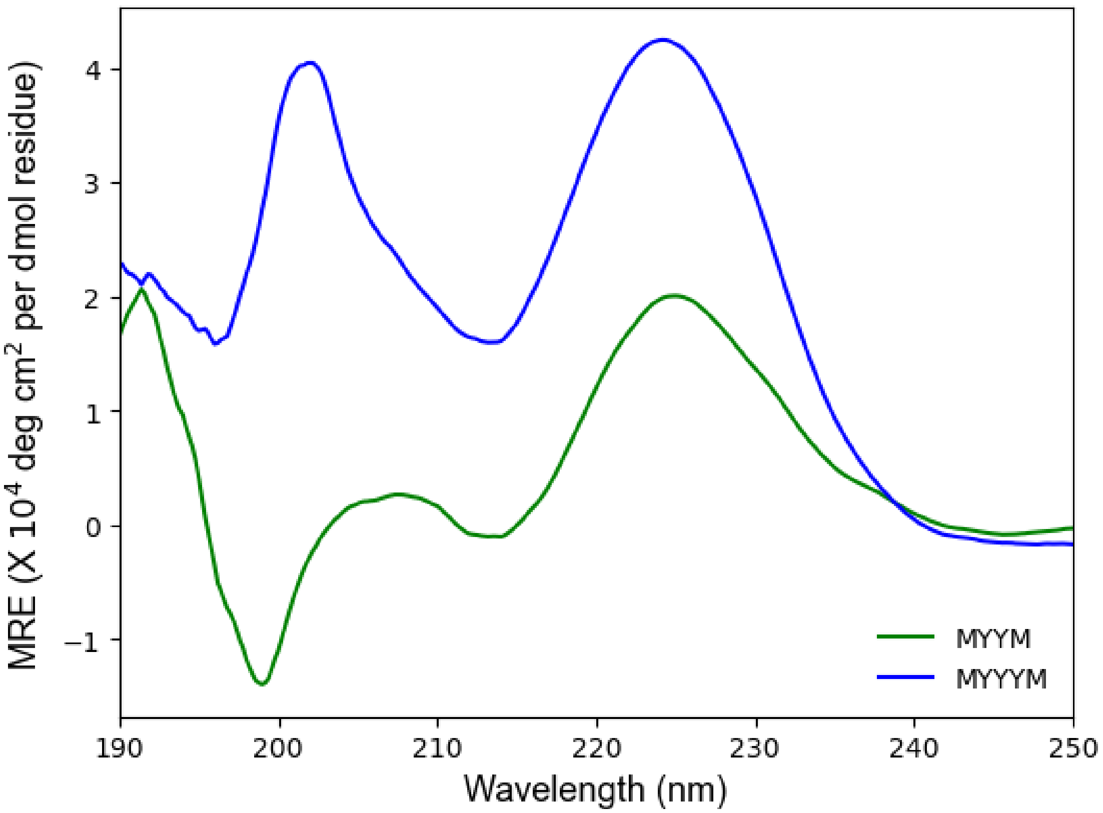

**Supplementary Figure 14**: Circular dichroism (CD) spectra for MYYM and MYYYM peptide sequences.

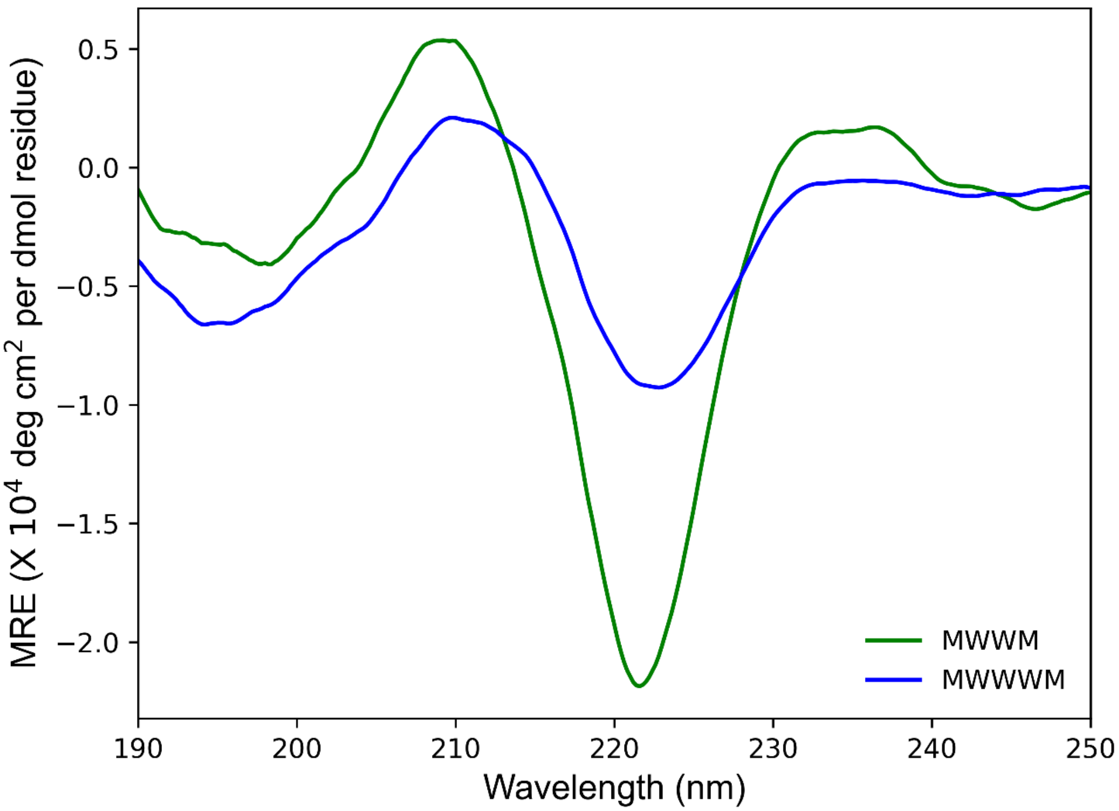

**Supplementary Figure 15**: Circular dichroism (CD) spectra for MWWM and MWWWM peptide sequences.

### S3. Single-molecule conductance experimental data (STM-BJ data)

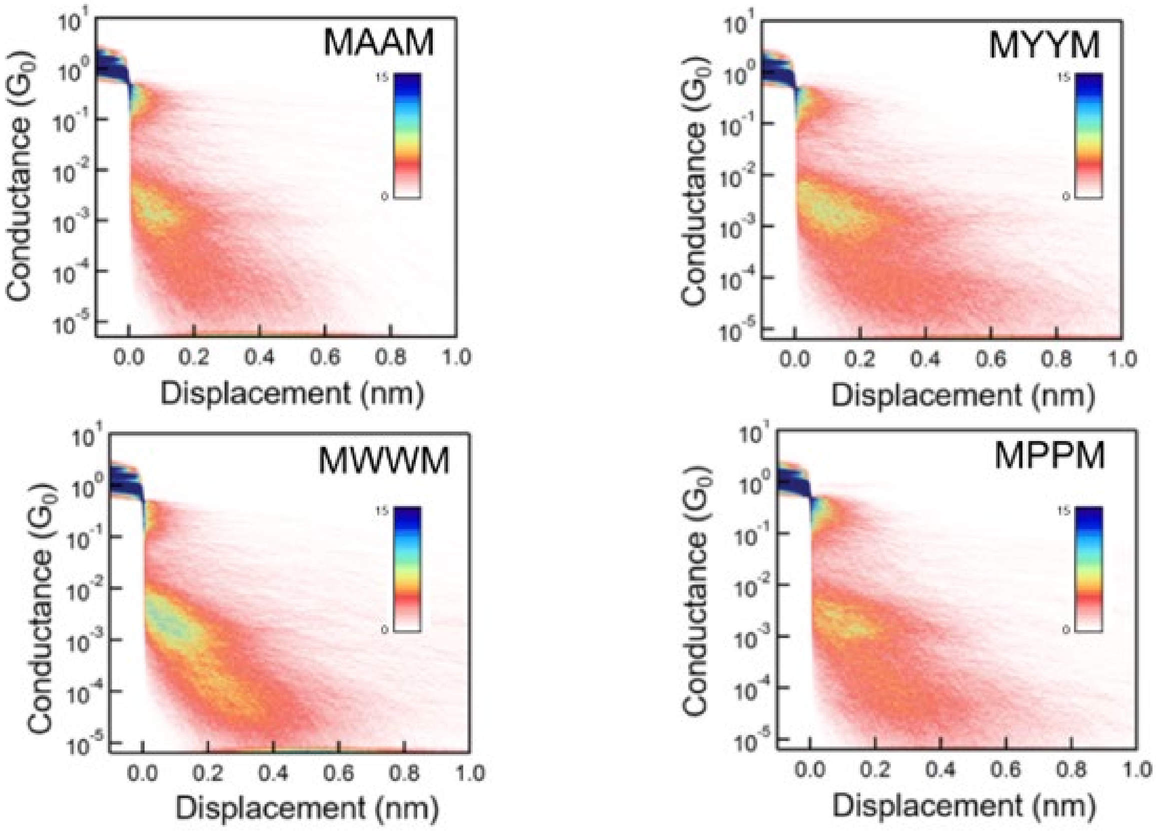

**Supplementary Fig. 16:** 2D conductance histograms for tetrapeptides at 250 mV. All tetramer sequences indicate a bimodal conductance distribution arising due to the conformational flexibility of the peptide backbone.

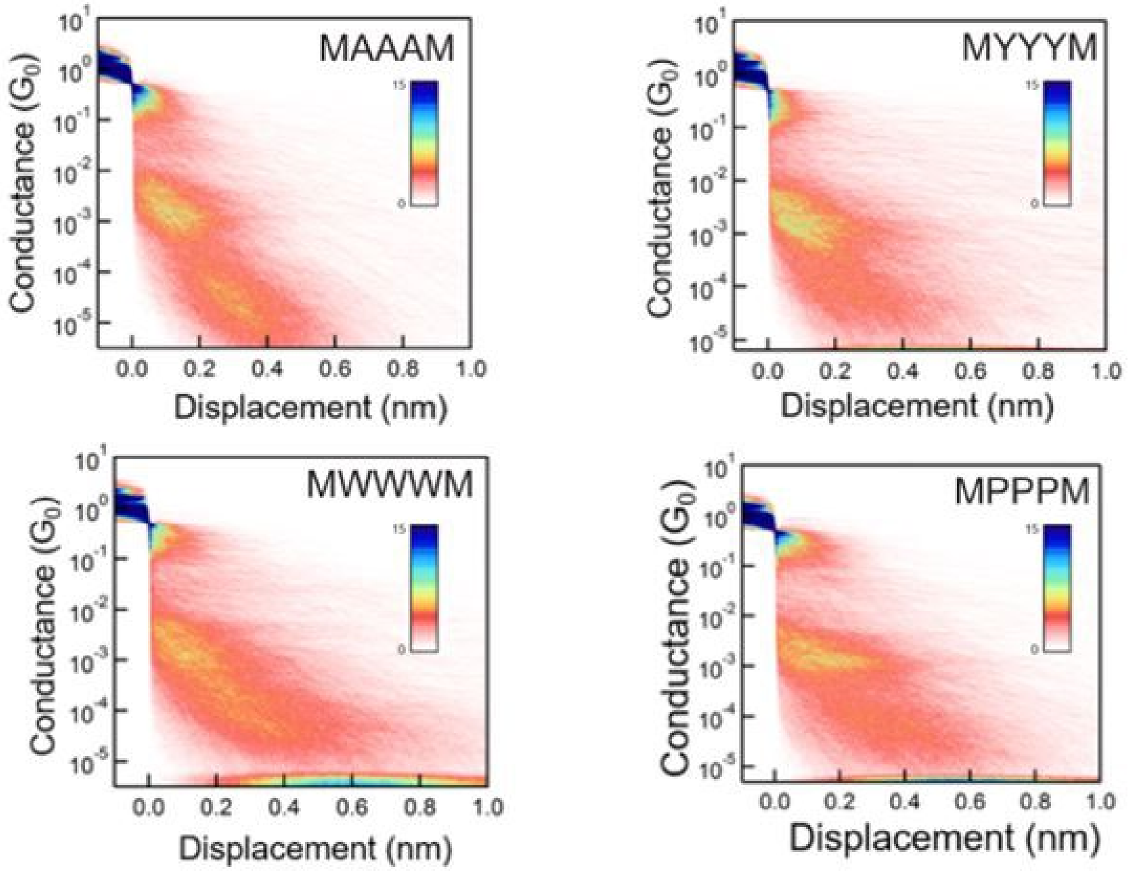

**Supplementary Fig. 17**: 2D conductance histograms for pentapeptides at 250 mV. All pentamer sequences indicate a bimodal conductance distribution arising due to the conformational flexibility of the peptide backbone.

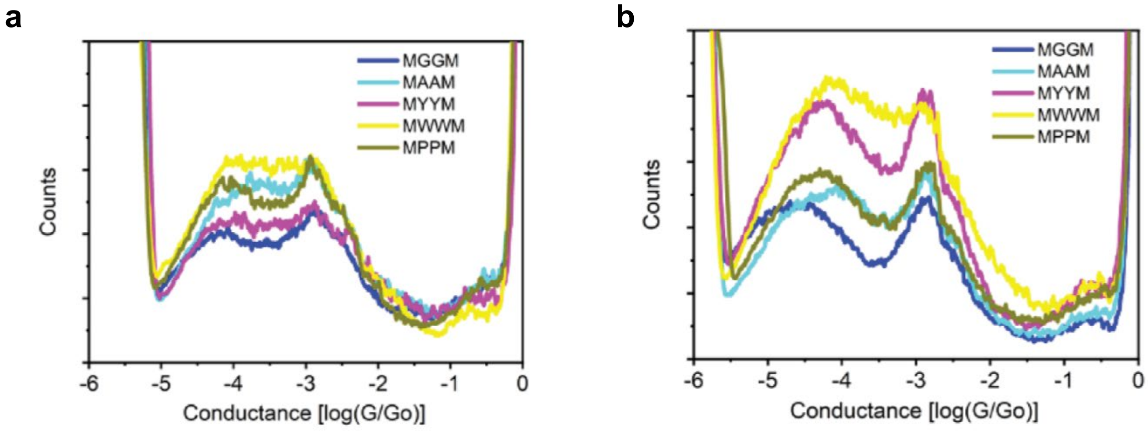

**Supplementary Fig. 18**: 1D conductance histogram for the tetrapeptides at applied bias of **(a)** 100 mV and **(b)** 400 mV. Multiple conductance populations are observed irrespective of applied bias.

**Table 1:** High conductance and low conductance peaks for the tetrapeptides at 250 mV. The conductance value, in log scale, for each peak (high conductance and low conductance) is determined from the center position of a Lorentzian fit to the peak.

| **Peptide sequence** | **High conductance peak** [log($G/G_0$)] | **Low conductance peak** [log($G/G_0$)] |
|---|---|---|
| Met-Ala-Ala-Met | -2.86 | -4.22 |
| Met-Gly-Gly-Met | -2.78 | -4.52 |
| Met-Tyr-Tyr-Met | -2.80 | -4.32 |
| Met-Trp-Trp-Met | -2.80 | -4.20 |
| Met-Pro-Pro-Met | -2.79 | -4.29 |

**Table 2**: High conductance and low conductance peaks for the pentapeptides at applied bias of 250 mV. The conductance value, in log scale, for each peak (high conductance and low conductance) is determined from the center position of a Lorentzian fit to the peak.

| **Peptide sequence** | **High conductance peak** [log($G/G_0$)] | **Low conductance peak** [log($G/G_0$)] |
|---|---|---|
| Met-Ala-Ala-Ala-Met | -2.91 | -4.63 |
| Met-Gly-Gly-Gly-Met | -2.88 | -4.75 |
| Met-Tyr-Tyr-Tyr-Met | -2.92 | -4.04 |
| Met-Trp-Trp-Trp-Met | -2.94 | -4.33 |
| Met-Pro-Pro-Pro-Met | -2.87 | -4.22 |

### S4. Gaussian Mixture modeling + silhouette score clustering

Unsupervised learning algorithms (such as K-means ++[1] and spectral clustering analysis[2]) have been used to analyze single-molecule charge transport data. To understand the bimodal conductance distribution observed for peptides, we employ a classification algorithm for data clustering based on Gaussian mixture modelling (GMM). GMM offers advantages over alternative methods such as K-means due to its ability to detect sub-populations of unequal covariance. Clustering is carried out using 1D and 2D conductance histograms (**Figure 2** and **Supplementary Figures 12-13**).

From each individual trace, a 30-by-30 two-dimensional histogram and a 100-bin one dimensional histogram are extracted. These are combined to form a 1000-dimensional feature space (30 × 30 + 100) over which GMM operates. The classification operates on the conductance range of 0 to -5.5 log($G/G_o$), and displacement range of -0.1 to 1 nm (0 nm corresponds to the point where the junction gets broken, the -0.1 nm to 0 nm regime indicates metal-metal contact. All traces are aligned at 0.5 $G_o$ as the starting point of displacement). Silhouette scores[3] are calculated for calculating the number of clusters based on GMM. Silhouette scores indicate how similar a feature is to its own cluster as compared to another cluster. Silhouette score value ranges from -1 to 1. For all the tetra- and pentapeptides, the silhouette score values are computed for the number of clusters varying from 2 to 5. The largest value of silhouette score is taken as an indication for optimal number of clusters. The goal of using silhouette scores is to find the optimal number of clusters for our bimodal conductance distribution and to observe if the two-conductance populations are part of a single trace, indicating conformation induced charged transport (dynamic heterogeneity), or they occur in separate traces (static heterogeneity), indicating different native conformations or distinct charge transport pathways.

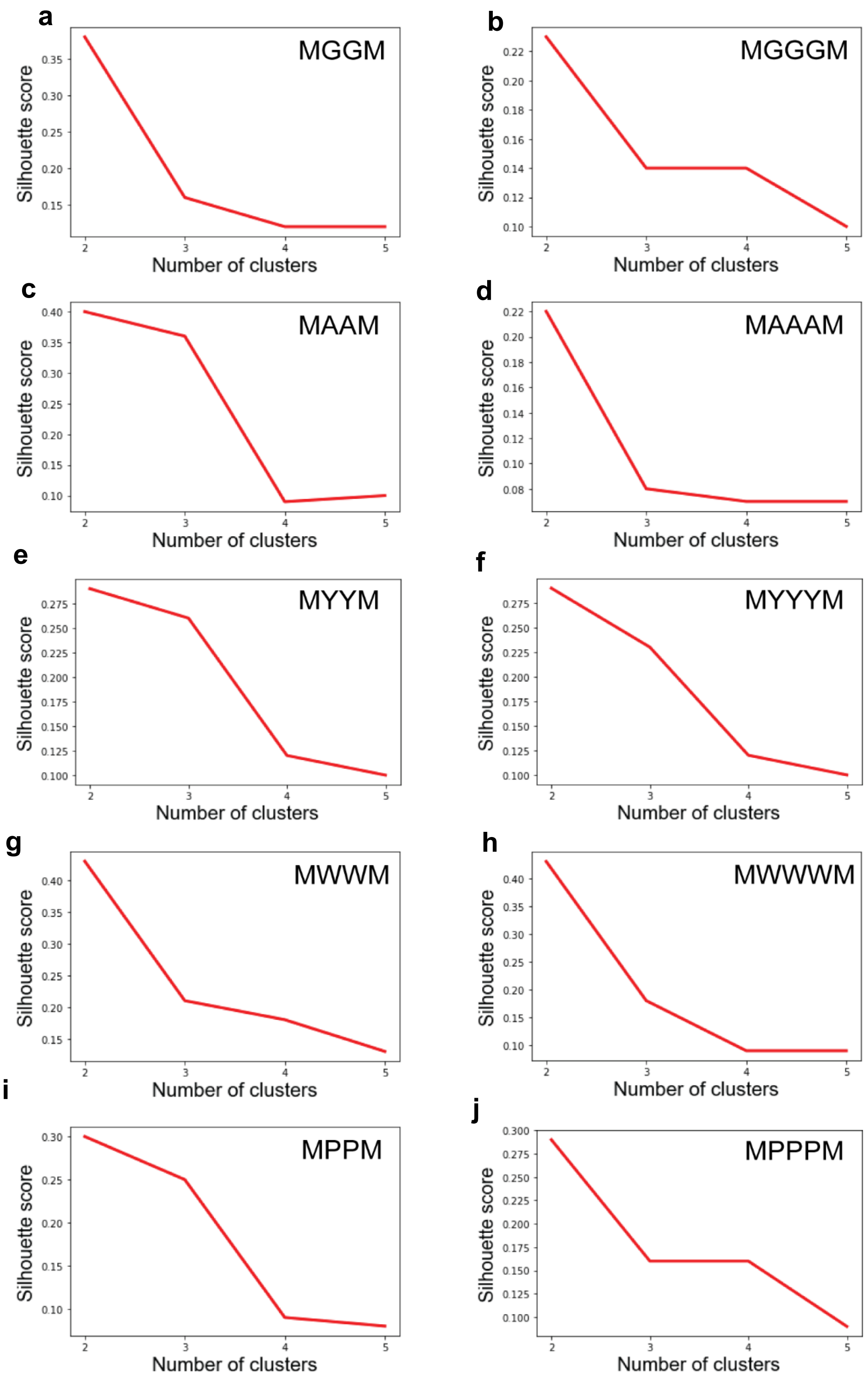

**Supplementary Fig. 19**: Silhouette scores tetra- and pentapeptides when the number of clusters is varied from 2 to 5. **a,c,e,g,i**. Tetramer sequences **b,d,f,h,j**. Pentamer sequences. The optimal number of clusters for all the tetra- and pentapeptides is 2.

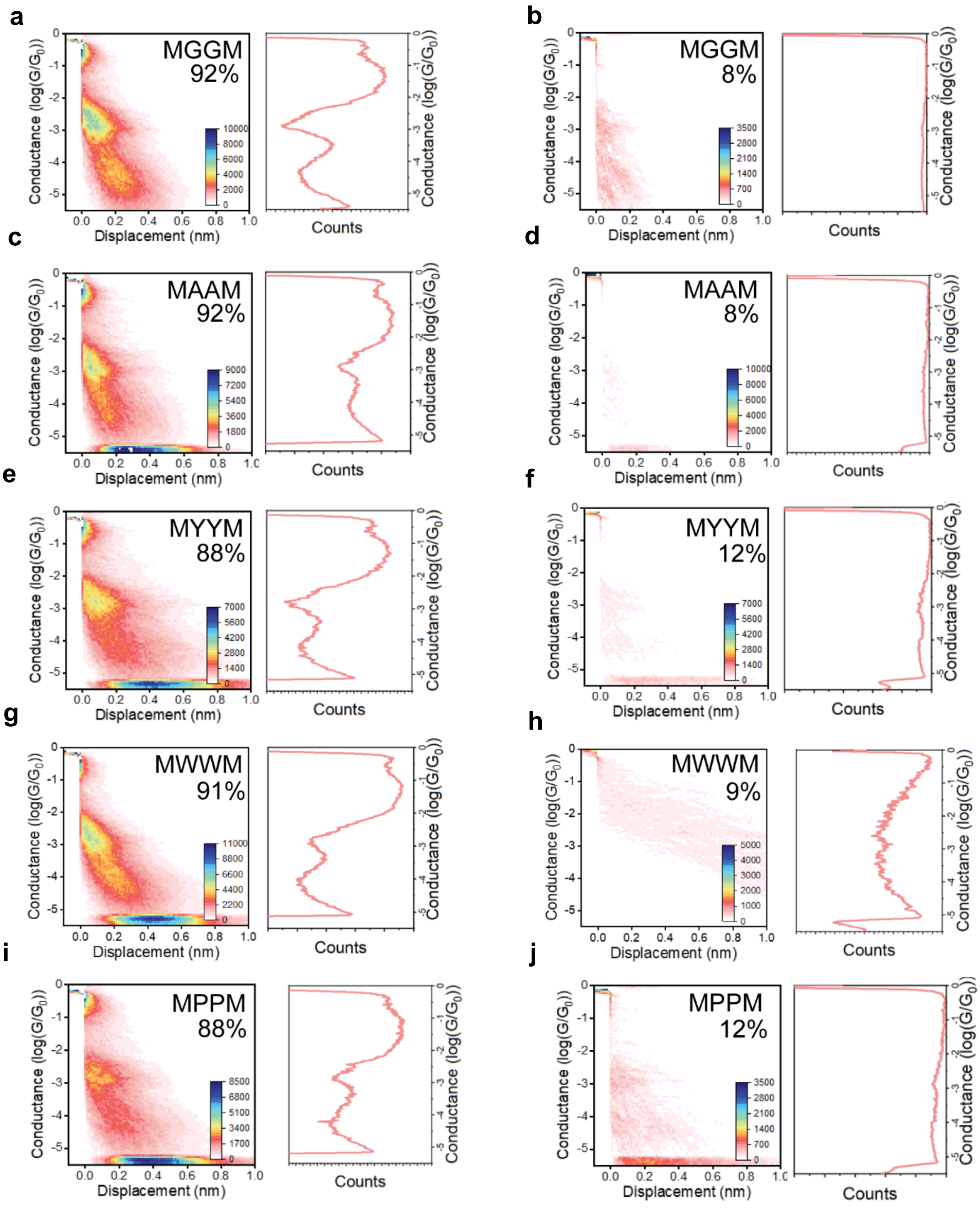

**Supplementary Fig. 20**: GMM for tetramer sequences **a,c,e,g,i.** Dominant cluster (85-95% of data) indicating traces in which both characteristic conductance populations appear together **b,d,f,h,j.** Traces in which no molecule is detected (or some background noise is observed).

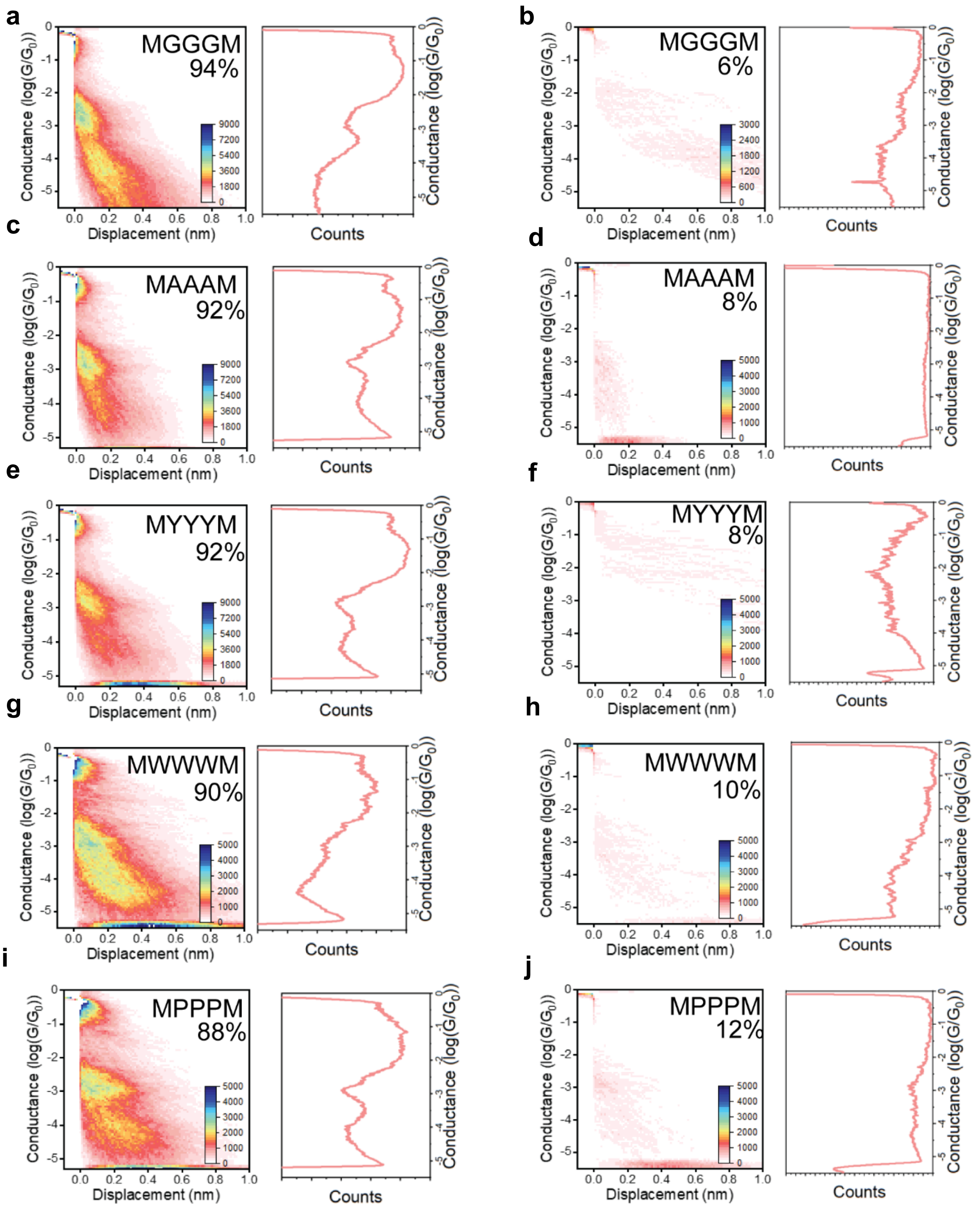

**Supplementary Fig. 21**: GMM for pentamer sequences **a,c,e,g,i**. Dominant cluster (85-95%) indicating traces in which both characteristic conductance populations appear together **b,d,f,h,j.** Traces in which no molecule is detected (or some background noise is observed)

### S5. MD simulations

The overall protocol and resulting collective variable distributions for MD simulations used in this study is described in Supplementary Fig. 17.

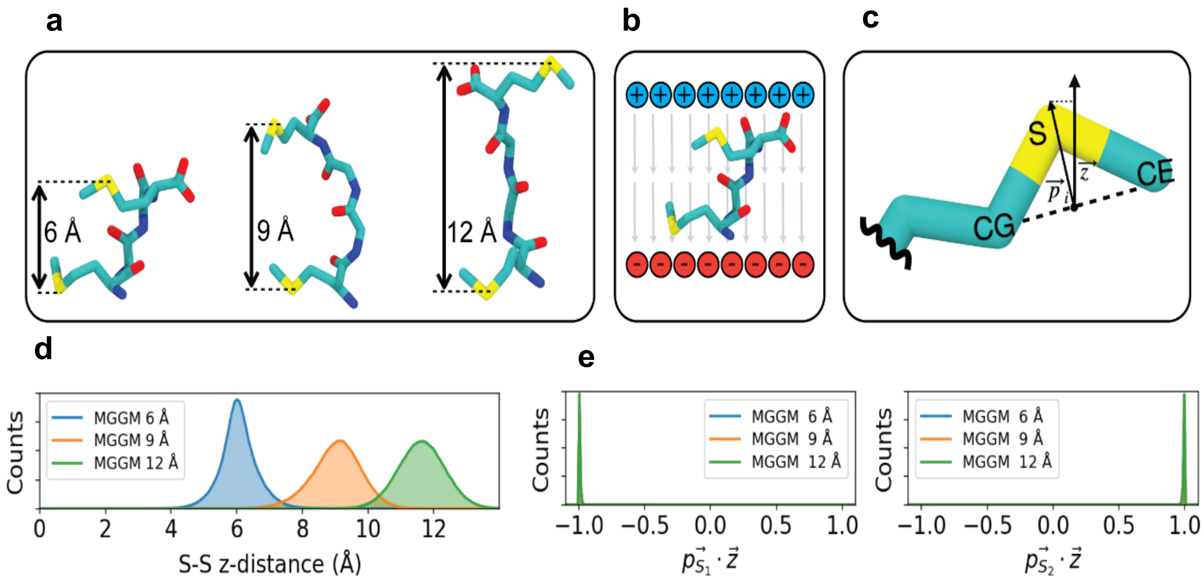

**Supplementary Figure 22:** Schematic illustration of implicit potentials utilized for MD simulations. **(a), (d)** Inter-anchor displacement potentials and resulting distributions at 6 Å, 9 Å, and 12 Å holding stages, corresponding to Equation 1 (main text). **(b)** Applied electric field defined in Equation 2 (main text). **(c), (e)** Sulfur-orienting potential and resulting distributions defined in Equation 3 (main text).

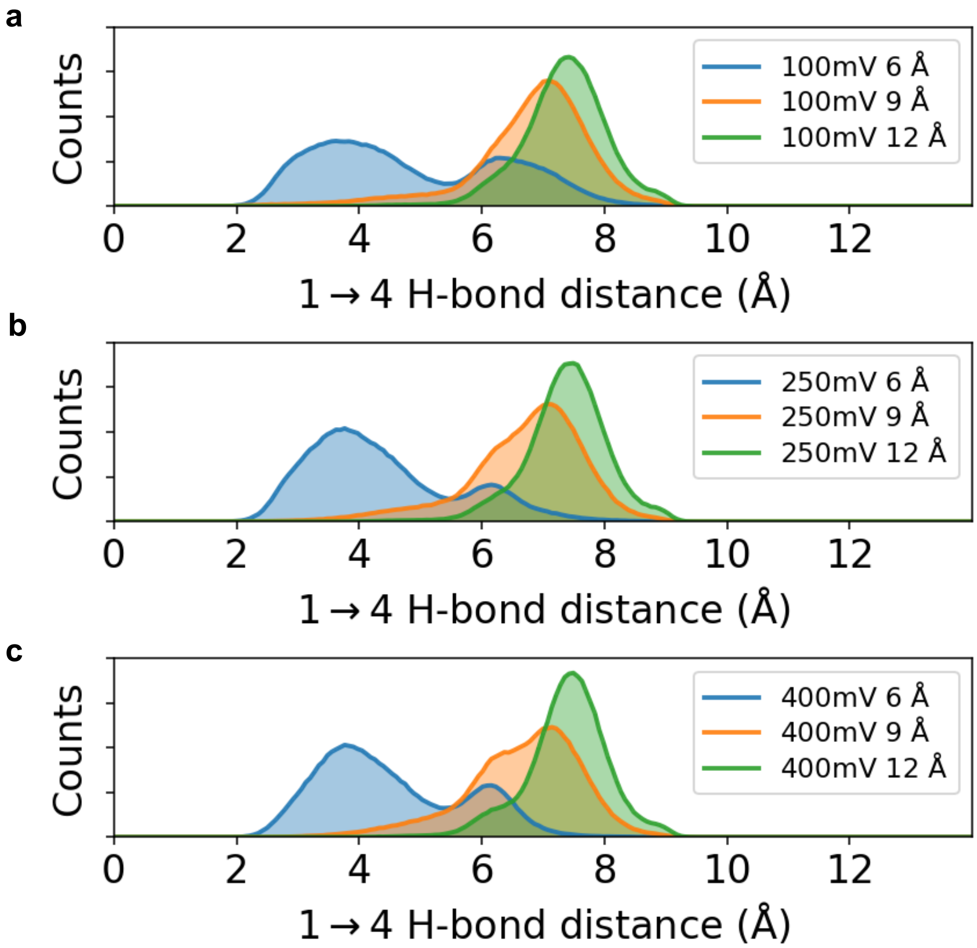

**Supplementary Figure 23**: Conformations observed in MD simulations are not significantly affected by a change in bias **a,b,c.** Backbone hydrogen bonding distance distribution for MGGM peptide sequence at (a) 100 mV, (b) 250 mV, and (c) 400 mV indicating elimination of intramolecular hydrogen bonds at higher displacement holding stages. It should be noted that the ground state charge distributions that dictate the partial charges in the equilibrium MD simulation will look very different from those on the atoms when there is current flowing through the molecule (charges and force fields do not represent the instantaneous charges of the junction).

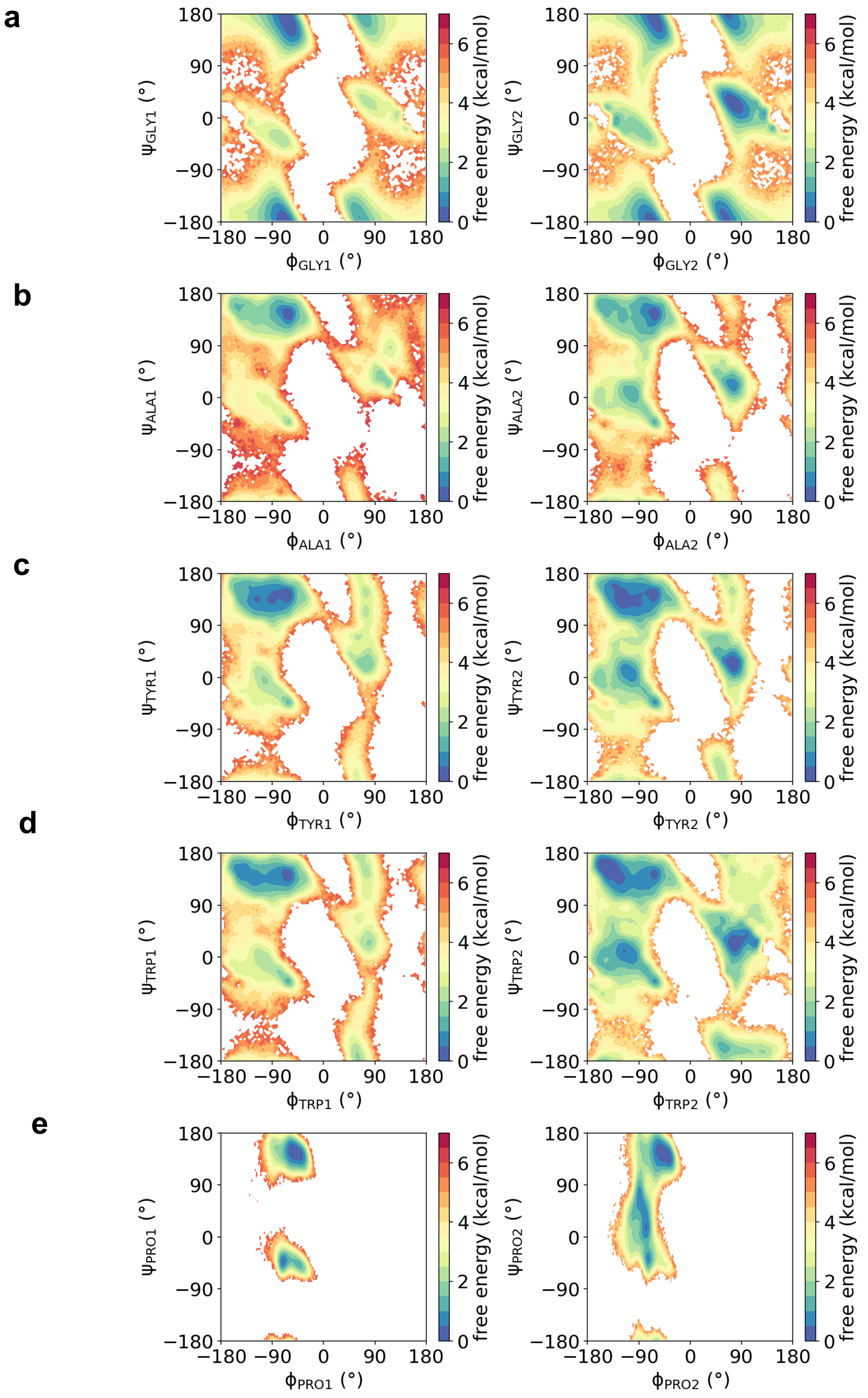

**Supplementary Figure 24**: Ramachandran free energy plots for non-terminal residues of tetrameric peptides, concatenated across all holding stages. (a) MGGM, (b) MAAM, (c) MYYM, (d) MWWM, (e) MPPM. Left column shows residue 1, right column shows residue 2 (0-based residue index).

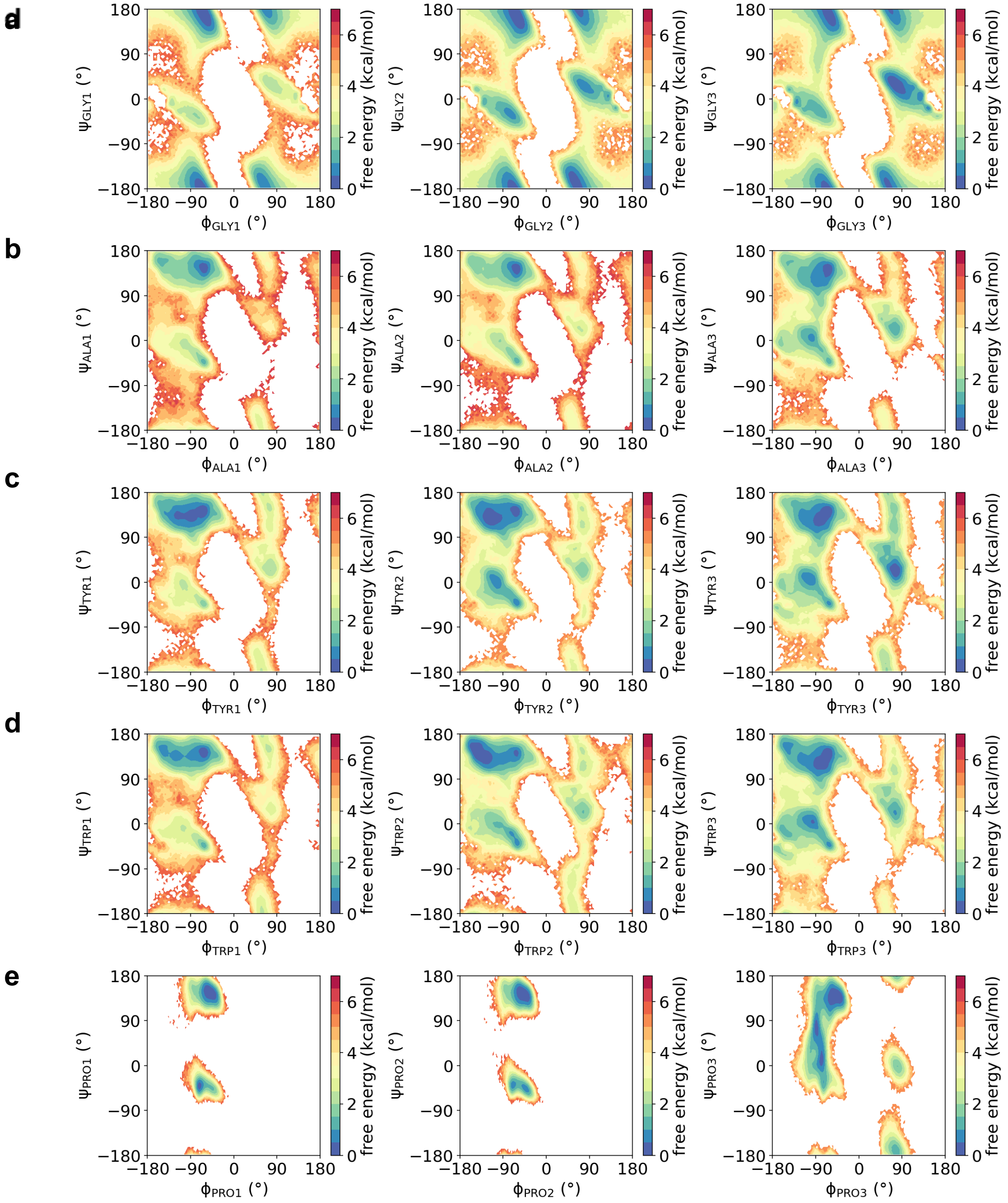

**Supplementary Figure 25**: Ramachandran free energy plots for non-terminal residues of pentameric peptides, concatenated across all holding stages. (a) MGGGM, (b) MAAAM, (c) MYYYM, (d) MWWWM, (e) MPPPM. Left column shows residue 1, middle column shows residue 2, and right column shows residue 3 (0-based residue index).

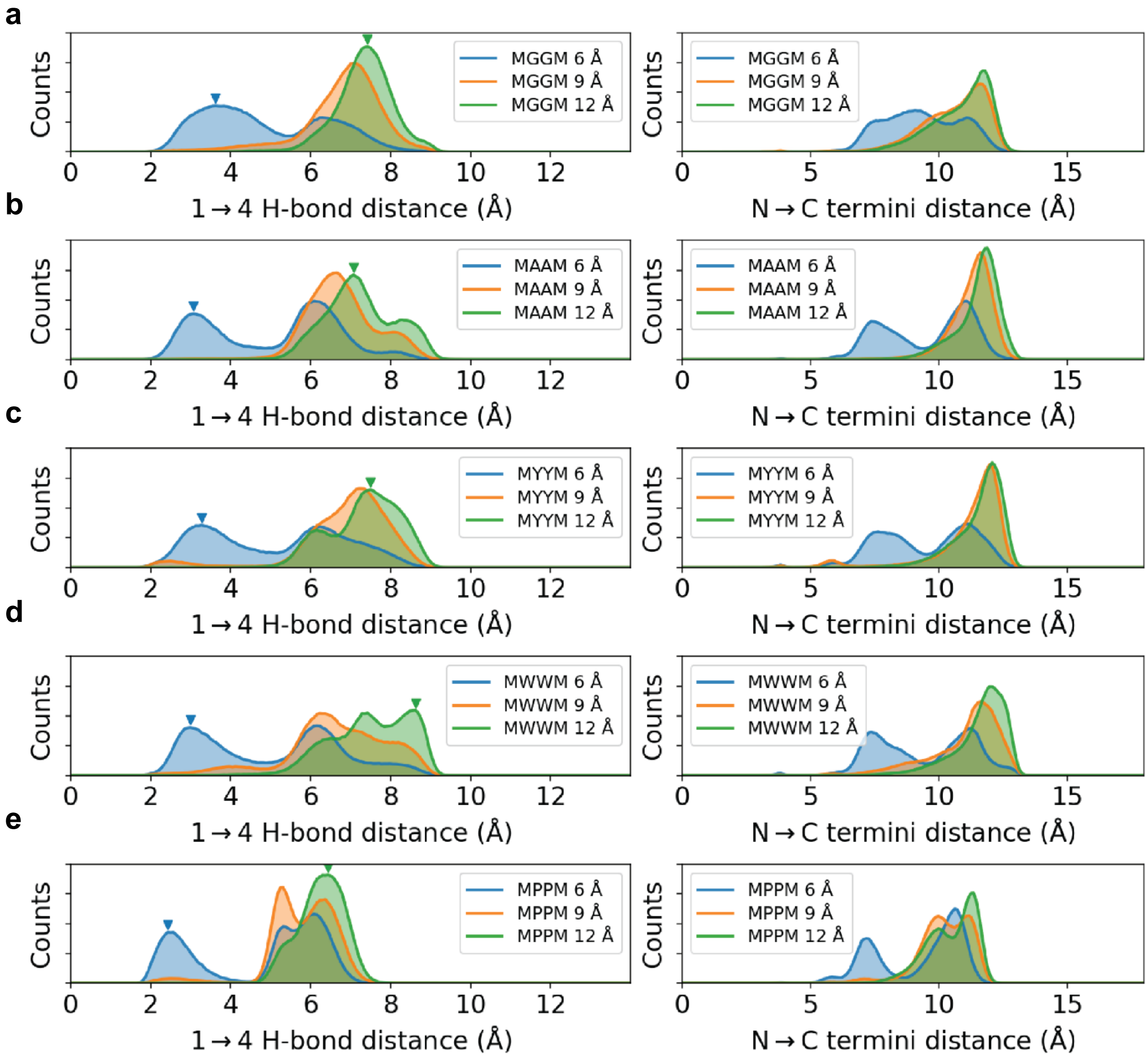

**Supplementary Figure 26:** MD simulations results for the tetrapeptides sequences. **a,b,c,d,e.** Backbone hydrogen bonding distance distribution for (a) MGGM, (b) MAAM, (c) MYYM, (d) MWWM, and (e) MPPM indicating elimination of intramolecular H-bonds at higher displacement holding stages. Inverted triangles in these figures indicate distribution peaks at which peptide conformers were selected for NEGF/DFT calculations.

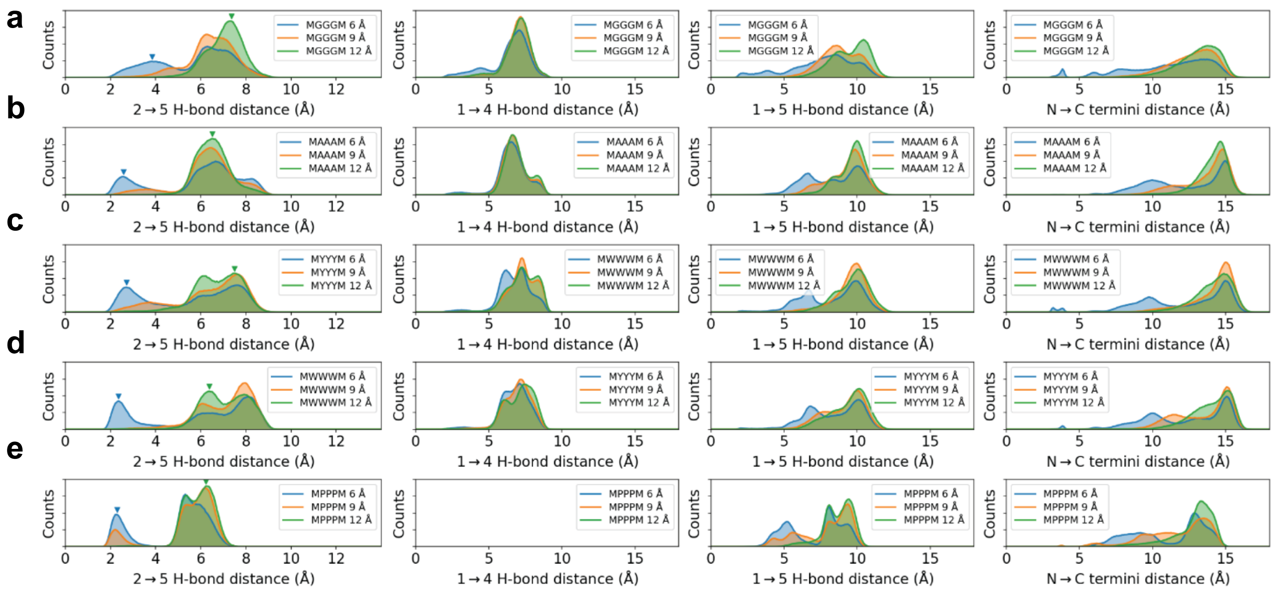

**Supplementary Figure 27:** MD simulation results for the pentapeptide sequences. **a,b,c,d,e.** Backbone hydrogen bonding distance distribution for (a) MGGGM, (b) MAAAM, (c) MYYYM, (d) MWWWM, and (e) MPPPM, indicating elimination of intramolecular H-bonds at higher displacement holding stages. Inverted triangles in these figures indicate distribution peaks at which peptide conformers were selected for NEGF/DFT calculations. Figures from left to right indicate different backbone intramolecular H-bonds. It should be noted that 1→4 H-bonding is absent in MPPPM due to the lack of a H-bond donor on position 4.

### S6. Principal component analysis (PCA)

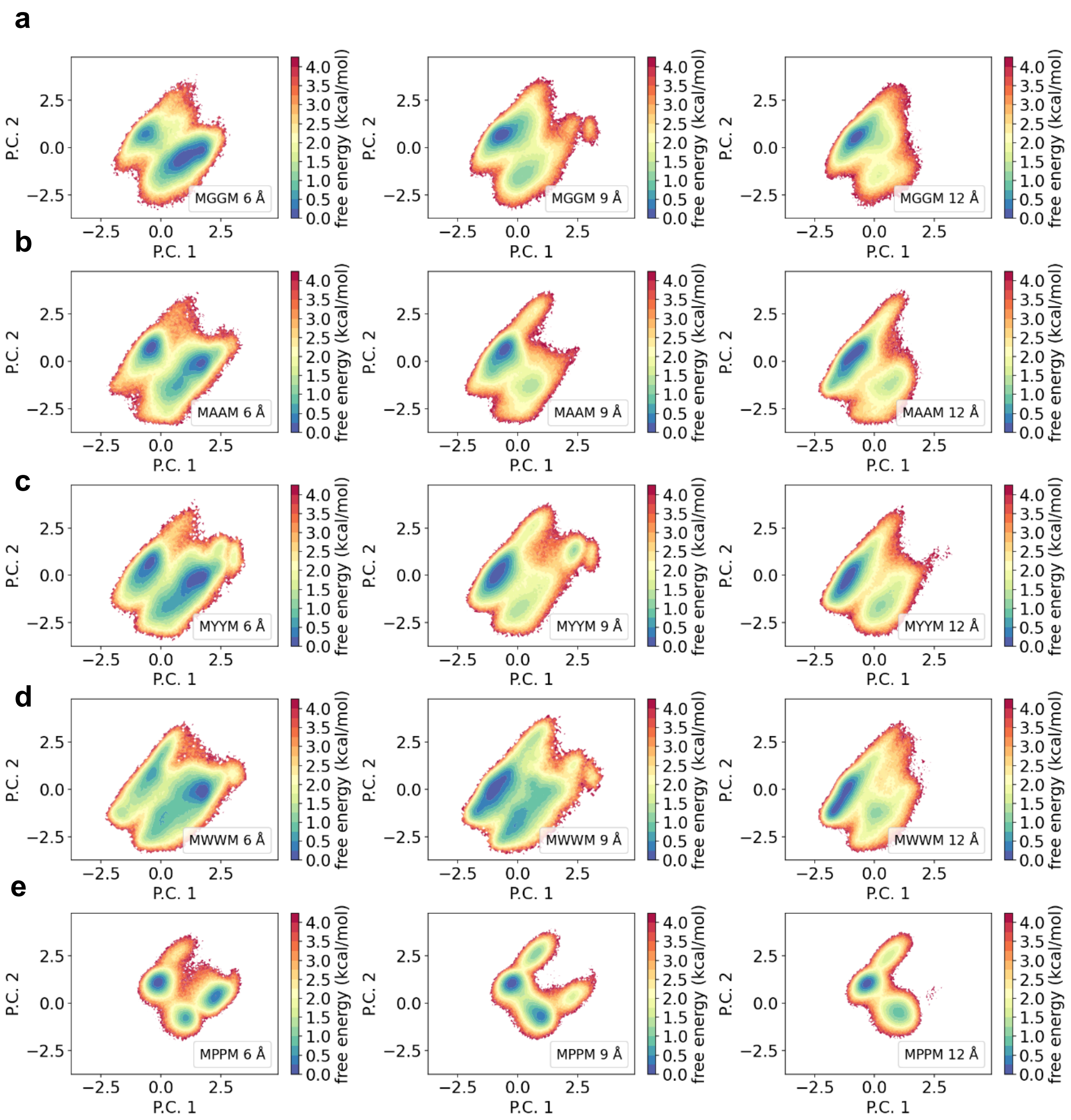

**Supplementary Figure 28:** Principal component analysis (PCA) results for tetrapeptides colored with respect to energy. Principal component projections for (a) MAAM, (b) MYYM, (c) MWWM and (d) MPPM conformational landscapes at various holding stages (left: 6 Å, middle: 9 Å, right: 12 Å).

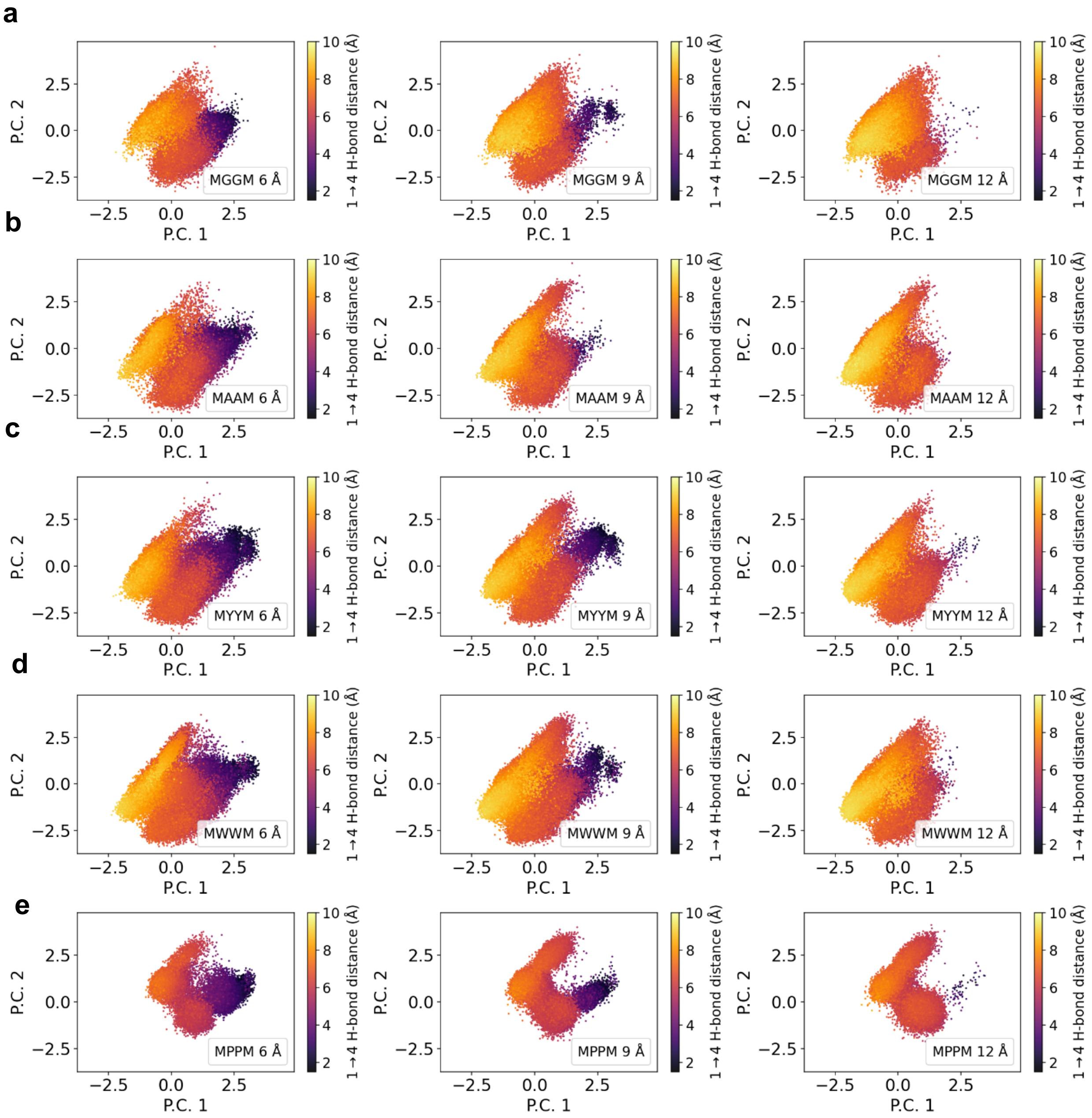

**Supplementary Figure 29:** Principal component analysis (PCA) results for tetrapeptides colored with respect to hydrogen bond distance. Principal component projections for (a) MAAM, (b) MYYM, (c)MWWM and (d) MPPM conformational landscapes at various holding stages (left: 6 Å, middle: 9 Å, right: 12 Å).

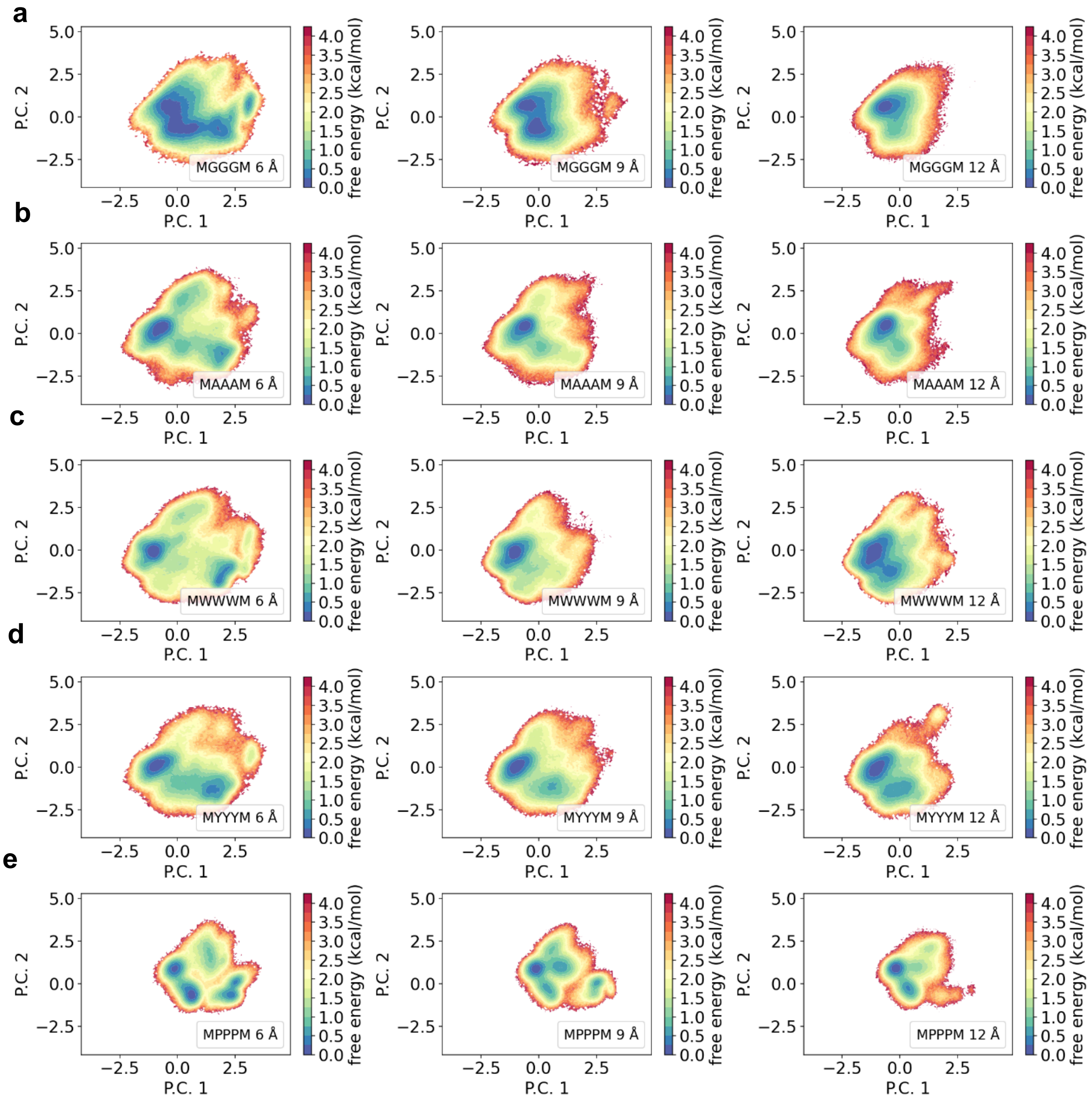

**Supplementary Figure 30:** Principal component analysis (PCA) results for pentapeptides colored with respect to energy. Principal component projections for (a) MAAAM, (b) MYYYM, (c)MWWWM and (d) MPPPM conformational landscapes at various holding stages (left: 6 Å, middle: 9 Å, right: 12 Å).

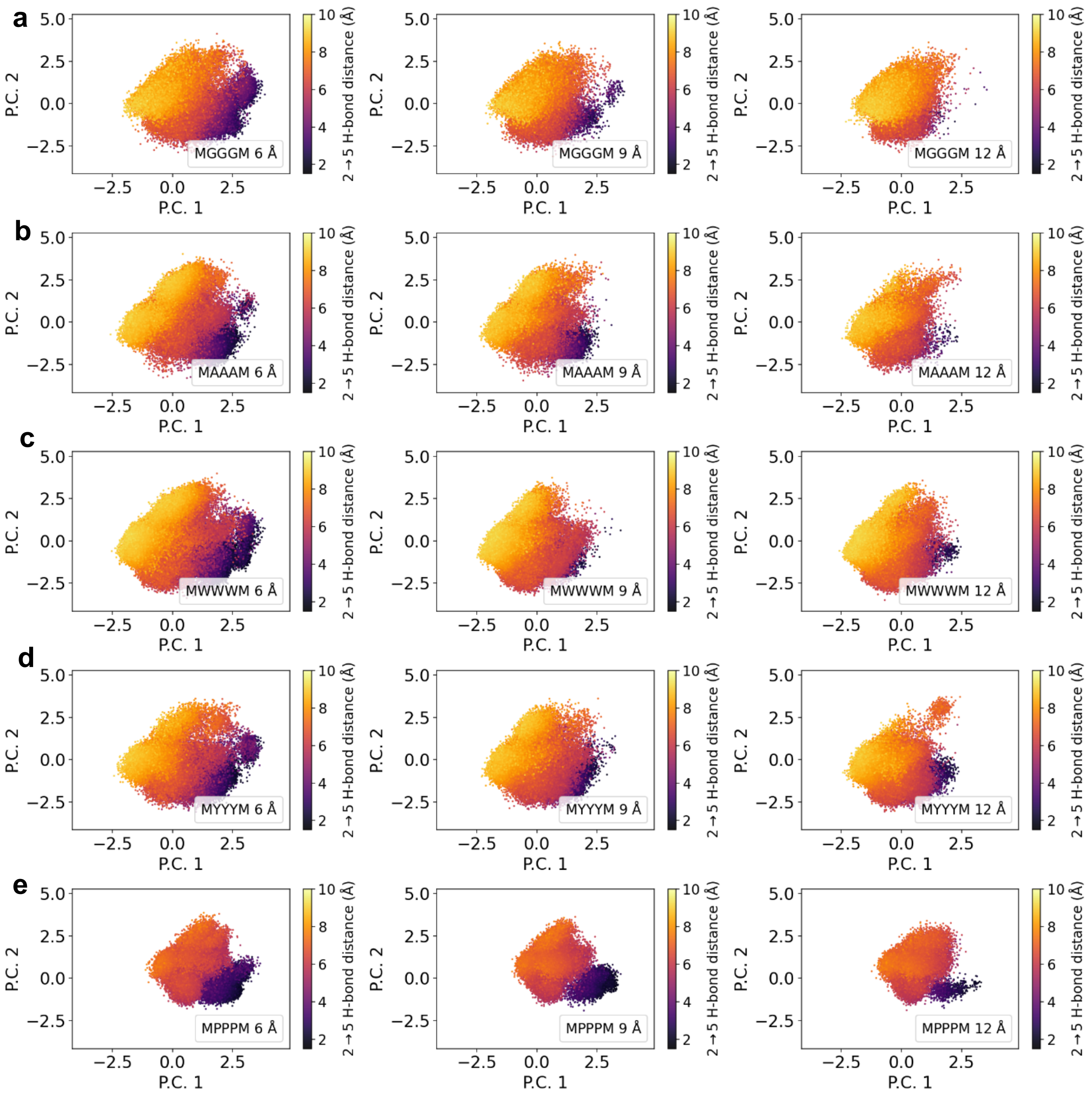

**Supplementary Figure 31:** Principal component analysis (PCA) results for pentapeptides colored with respect to hydrogen bond distance. Principal component projections for (a) MAAAM, (b) MYYYM, (c) MWWWM and (d) MPPPM conformational landscapes at various holding stages (left: 6 Å, middle: 9 Å, right: 12 Å).

### S7. NEGF-DFT simulations

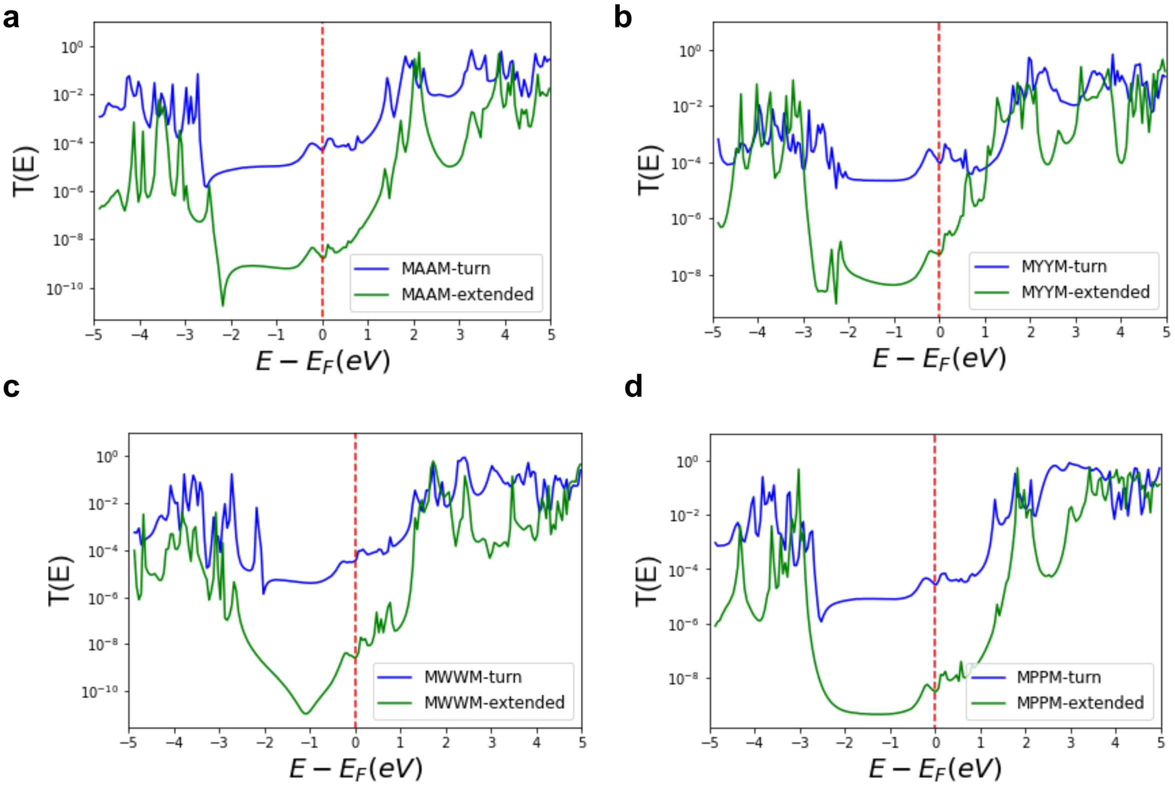

**Supplementary Figure 32:** NEGF-DFT calculations for tetrapeptides. Transmission as a function of energy (relative to the Fermi energy level) for (a) MAAM (b) MYYM (c) MWWM, and (d) MPPM. Significant differences at $E - E_F = 0$ are observed for the turn (blue) and the extended (green) configurations.

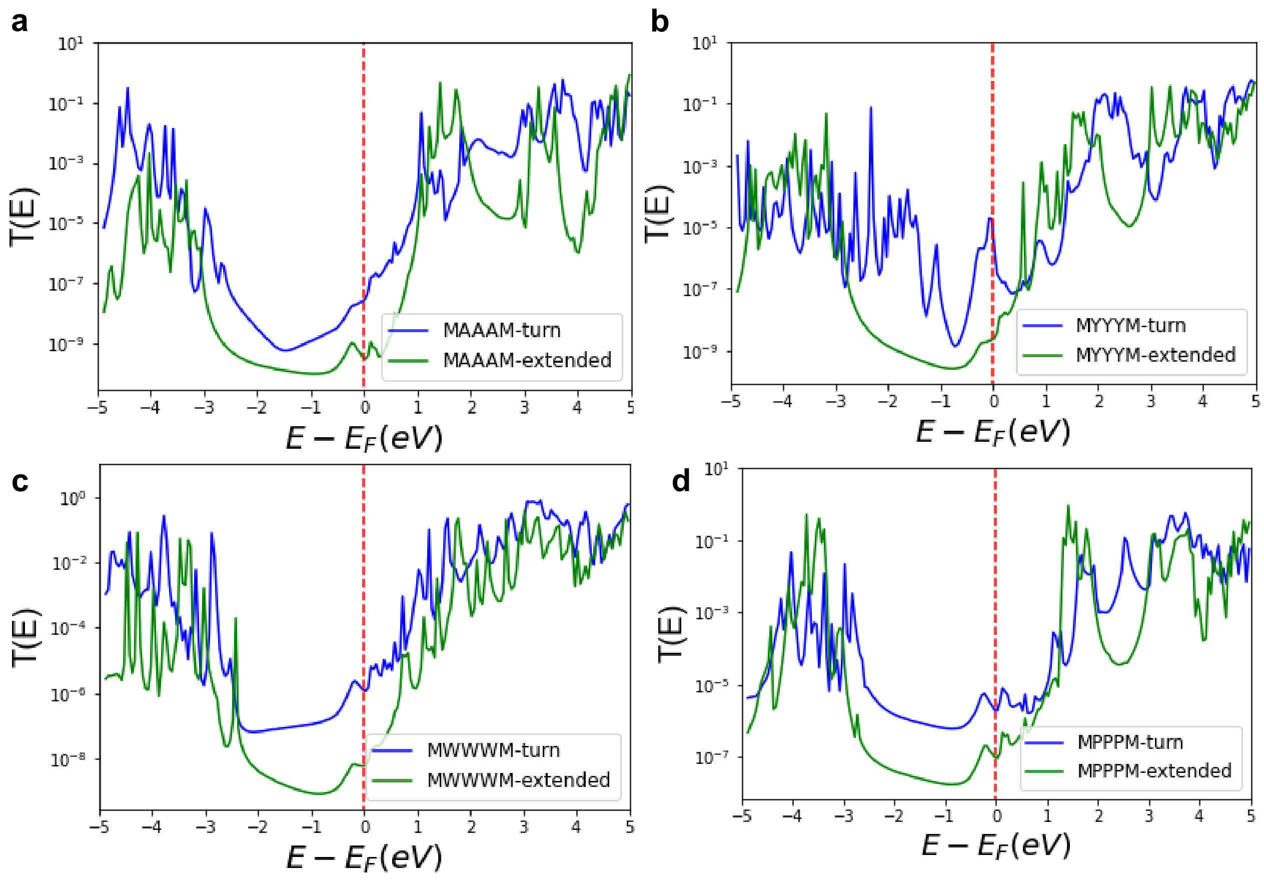

**Supplementary Figure 33:** NEGF-DFT calculations for pentapeptides. **a,b,c,d**. Transmission as a function of energy (relative to the Fermi energy level) for (a) MAAAM (b) MYYYM (c) MWWWM, and (d) MPPPM. Significant differences at $E - E_F = 0$ are observed for the turn (blue) and the extended (green) configurations.

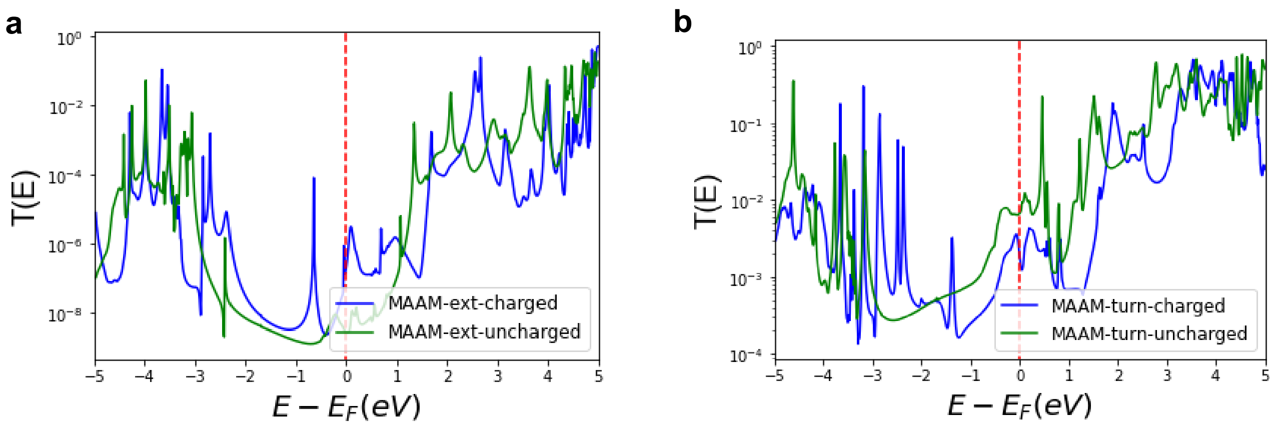

**Supplementary Figure 34:** Transmission probabilities showing differences between charged and uncharged species. Transmission versus energy (relative to Fermi Energy level) for (**a**) MAAM-ext configuration for the charged and uncharged states and (**b**) MAAM-turn configuration for the charged and uncharged states.

### S8. Comparison of experimental and computational transmission values

**Table 3**: Comparison of experimental and computational high conductance state value ($G/G_0$) for tetrapeptides

| Sequence | Computational value | Experimental value |
|---|---|---|
| MGGM | $1.63 \times 10^{-5}$ | $1.65 \times 10^{-3}$ |
| MAAM | $4.93 \times 10^{-5}$ | $1.38 \times 10^{-3}$ |
| MPPM | $2.78 \times 10^{-5}$ | $1.62 \times 10^{-3}$ |
| MWWM | $3.59 \times 10^{-5}$ | $1.58 \times 10^{-3}$ |
| MYYM | $9.75 \times 10^{-5}$ | $1.58 \times 10^{-3}$ |

**Table 4**: Comparison of experimental and computational low conductance state value ($G/G_0$) for tetrapeptides

| Sequence | Computational value | Experimental value |
|---|---|---|
| MGGM | $9.63 \times 10^{-11}$ | $3.01 \times 10^{-5}$ |
| MAAM | $1.71 \times 10^{-9}$ | $6.02 \times 10^{-5}$ |
| MPPM | $3.10 \times 10^{-9}$ | $5.10 \times 10^{-5}$ |
| MWWM | $2.70 \times 10^{-9}$ | $6.30 \times 10^{-5}$ |
| MYYM | $5.28 \times 10^{-8}$ | $4.78 \times 10^{-5}$ |

**Table 5**: Comparison of experimental and computational high conductance state value ($G/G_0$) for pentapeptides

| Sequence | Computational value | Experimental value |
|---|---|---|
| MGGGM | $4.63 \times 10^{-7}$ | $1.31 \times 10^{-3}$ |
| MAAAM | $2.64 \times 10^{-8}$ | $1.23 \times 10^{-3}$ |
| MPPPM | $1.95 \times 10^{-6}$ | $1.34 \times 10^{-3}$ |
| MWWWM | $1.22 \times 10^{-6}$ | $1.14 \times 10^{-3}$ |
| MYYYM | $1.04 \times 10^{-5}$ | $1.20 \times 10^{-3}$ |

**Table 6**: Comparison of experimental and computational low conductance state value ($G/G_0$) for pentapeptides

| Sequence | Computational value | Experimental value |
|---|---|---|
| MGGGM | $4.24 \times 10^{-10}$ | $1.77 \times 10^{-5}$ |
| MAAAM | $3.11 \times 10^{-10}$ | $2.34 \times 10^{-5}$ |
| MPPPM | $9.42 \times 10^{-8}$ | $6.02 \times 10^{-5}$ |
| MWWWM | $6.03 \times 10^{-9}$ | $4.64 \times 10^{-5}$ |
| MYYYM | $2.37 \times 10^{-9}$ | $9.10 \times 10^{-5}$ |

### S9. Projected density of states (PDOS)

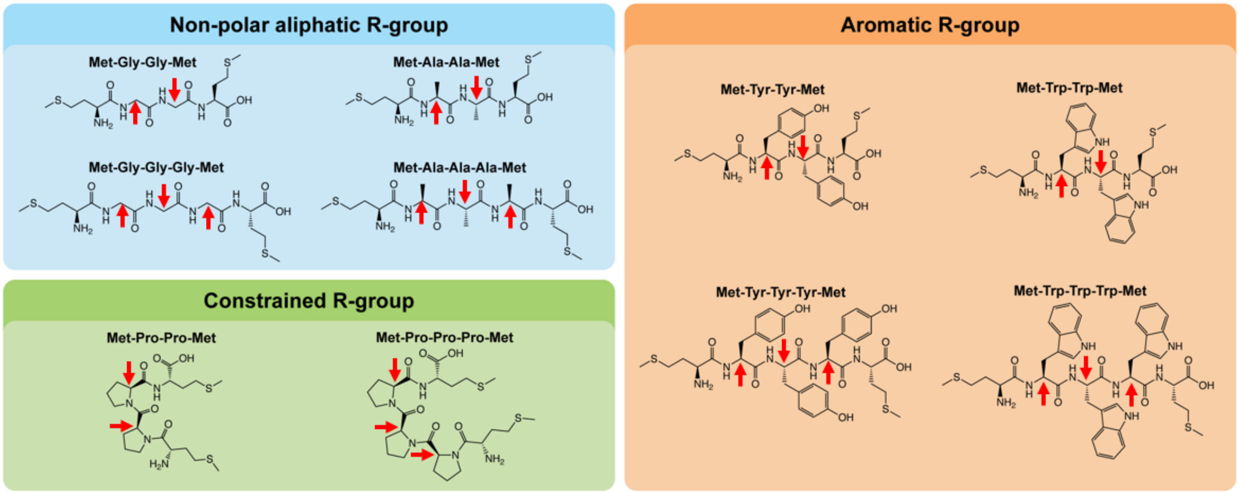

**Supplementary Figure 35**: Site specific projected density of states (PDOS) calculations for carbon atoms on the peptide backbone for which the side chain is attached. Red arrows indicate these carbon atoms.

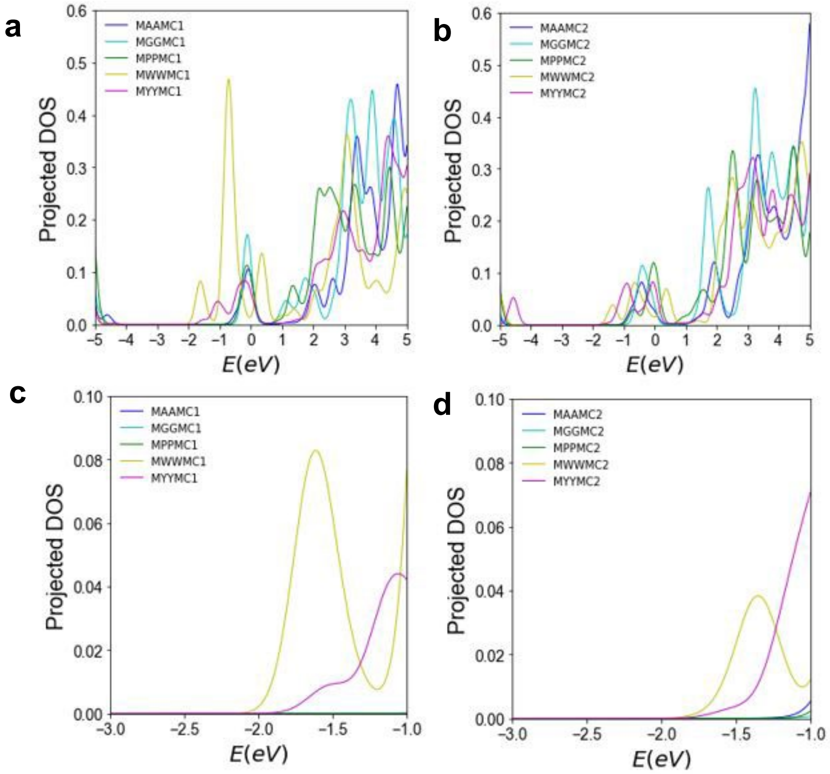

**Supplementary Figure 36**: Site specific PDOS calculations of all tetrapeptides. (a), (b) 1st and 2nd carbon atom with the side chain from -5 to 5 eV. (c), (d) 1st and 2nd carbon atom with the side chain around the Fermi energy level.

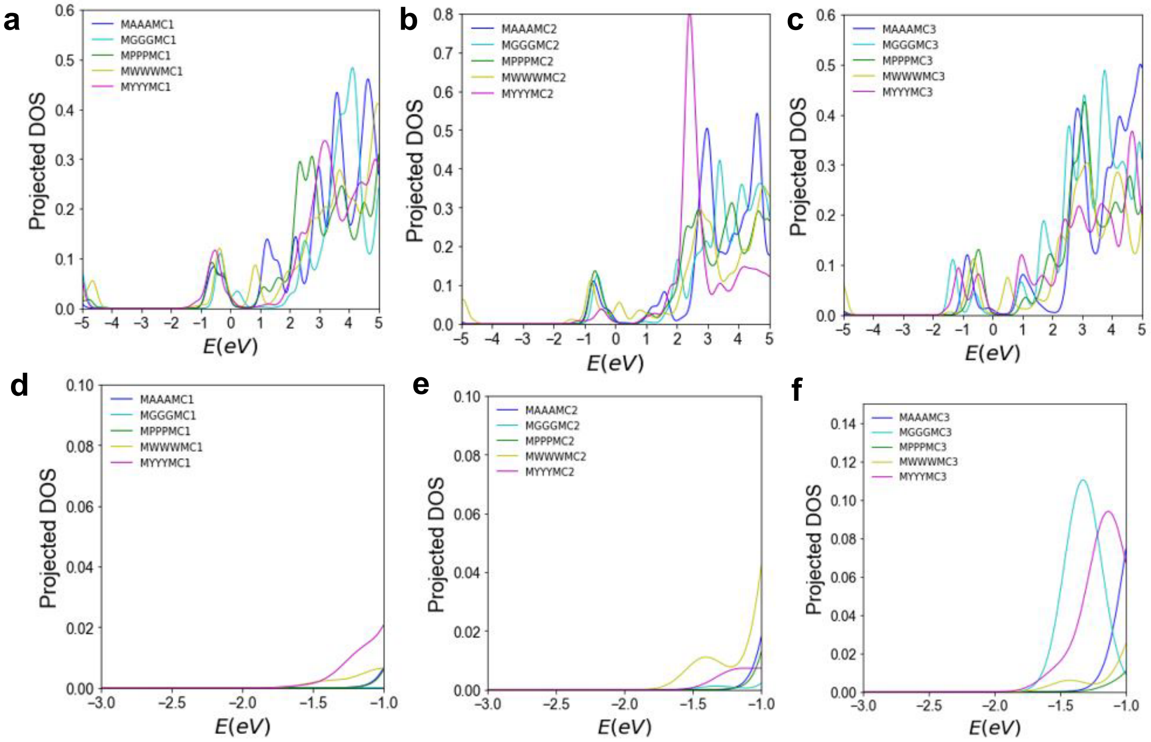

**Supplementary Figure 37:** Site specific PDOS calculations of all pentapeptides. (a), (b), (c) 1st, 2nd, and 3rd carbon atom with the side chain from -5 to 5 eV. (d), (e), (f) 1st, 2nd, and 3rd carbon atom with the side chain around the Fermi energy level.

**Table 7**: PDOS values near Fermi energy level for 1st and 2nd carbon atom with a side chain for all tetrapeptides.

| Sequence | 1st carbon atom with sidechain | 2nd carbon atom with sidechain |
|---|---|---|
| MAAM | 2.21 X $10^{-5}$ | 2.21 X $10^{-5}$ |
| MGGM | 1.14 X $10^{-6}$ | 3.06 X $10^{-6}$ |
| MPPM | 6.31 X $10^{-6}$ | 1.29 X $10^{-5}$ |
| MWWM | 6.96 X $10^{-5}$ | 4.28 X $10^{-5}$ |
| MYYM | 7.88 X $10^{-5}$ | 2 X $10^{-5}$ |

**Table 8**: PDOS values near Fermi energy level for 1st, 2nd, and 3rd carbon atom with a side chain for all pentapeptides.

| Sequences | 1st carbon atom with a side chain | 2nd carbon atom with a side chain | 3rd carbon atom with a side chain |
|---|---|---|---|
| MAAAM | 2.21 X $10^{-5}$ | 2.78 X $10^{-6}$ | 6 X $10^{-5}$ |
| MGGGM | 1 X $10^{-5}$ | 1.68 X $10^{-6}$ | 6.62 X $10^{-6}$ |
| MPPPM | 9.62 X $10^{-6}$ | 1.75 X $10^{-5}$ | 4.22 X $10^{-5}$ |
| MWWWM | 1.16 X $10^{-5}$ | 2.98 X $10^{-5}$ | 2.51 X $10^{-6}$ |
| MYYYM | 1.76 X $10^{-5}$ | 6.20 X $10^{-7}$ | 1.23 X $10^{-4}$ |

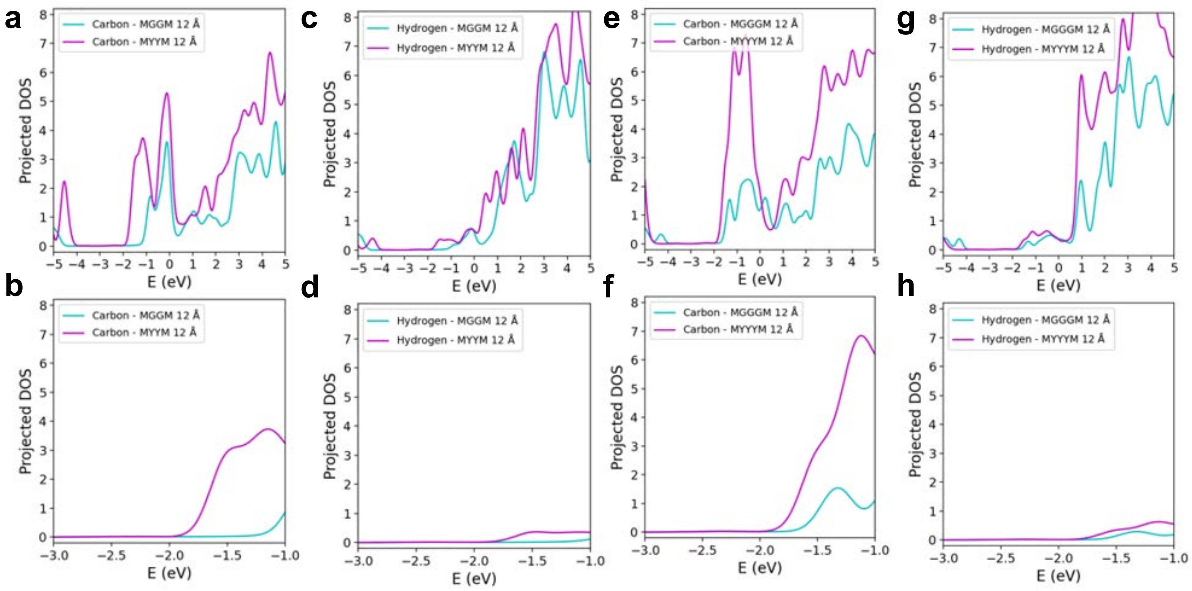

**Supplementary Figure 38**: All carbon atoms PDOS for MGGM and MYYM from (a) -5 to 5 eV and (b) near Fermi energy level. All hydrogen atoms PDOS for MGGM and MYYM from (c) -5 to 5 eV and (d) near Fermi energy level. All carbon atoms PDOS for MGGGM and MYYYM from (e) -5 to 5 eV and (f) near Fermi energy level. All hydrogen atoms PDOS for MGGM and MYYM from (g) -5 to 5 eV and (h) near Fermi energy level.

### S10. Orbital visualization

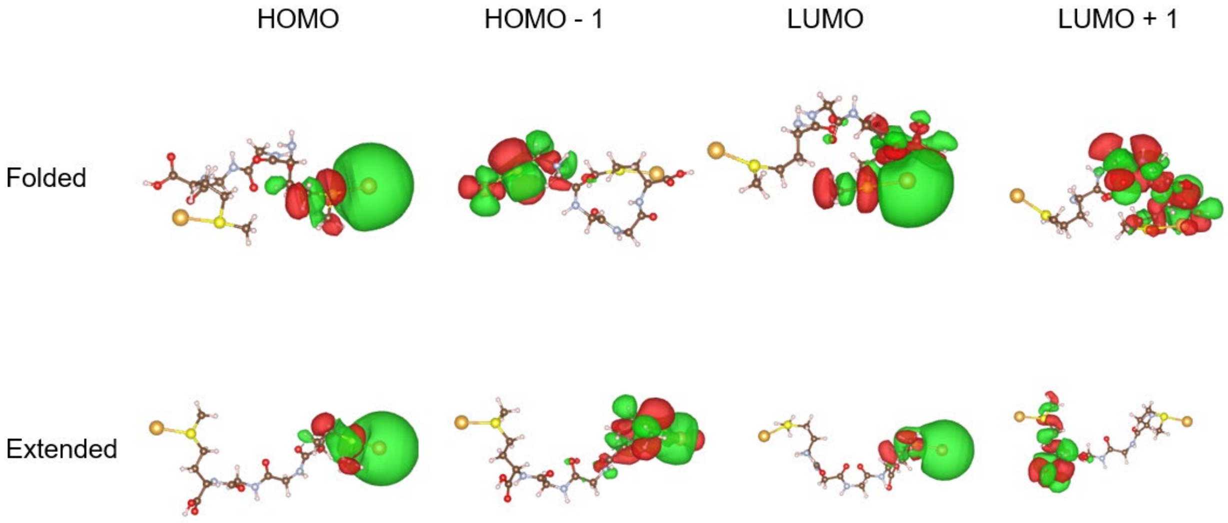

**Supplementary Figure 39**: Isosurface plots for HOMO, HOMO-1, LUMO and LUMO +1 orbital visualization for MGGM

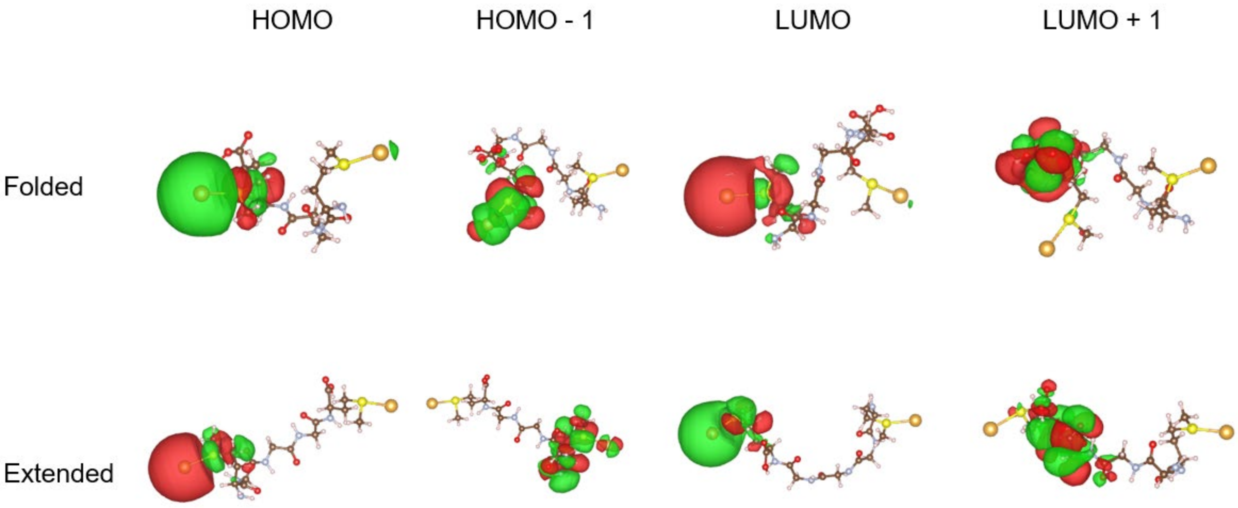

**Supplementary Figure 40**: Isosurface plots for HOMO, HOMO-1, LUMO and LUMO +1 orbital visualization for MGGGM

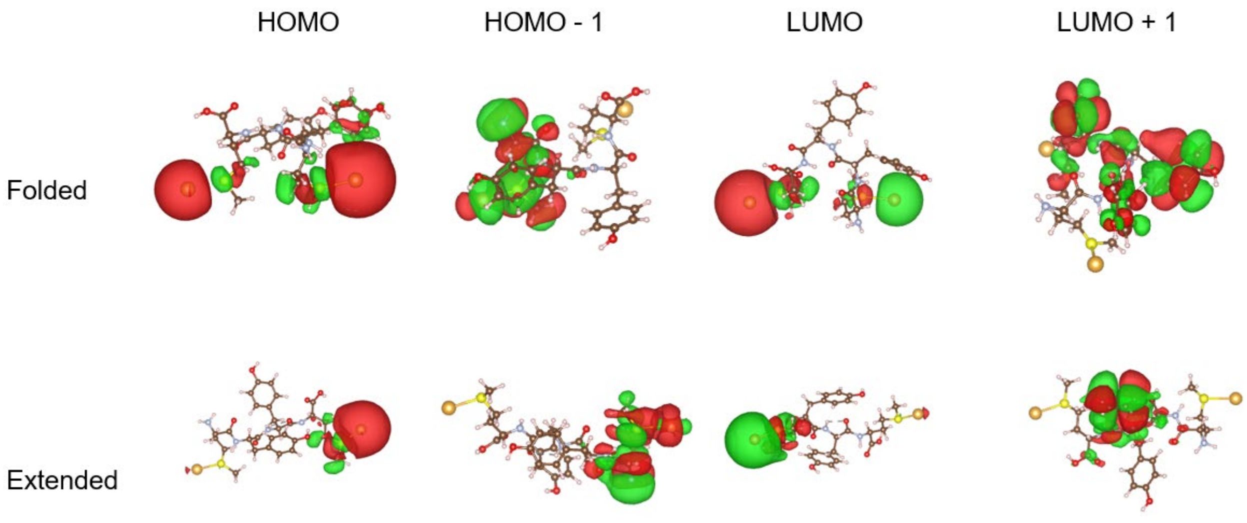

**Supplementary Figure 41**: Isosurface plots for HOMO, HOMO-1, LUMO and LUMO +1 orbital visualization for MYYM

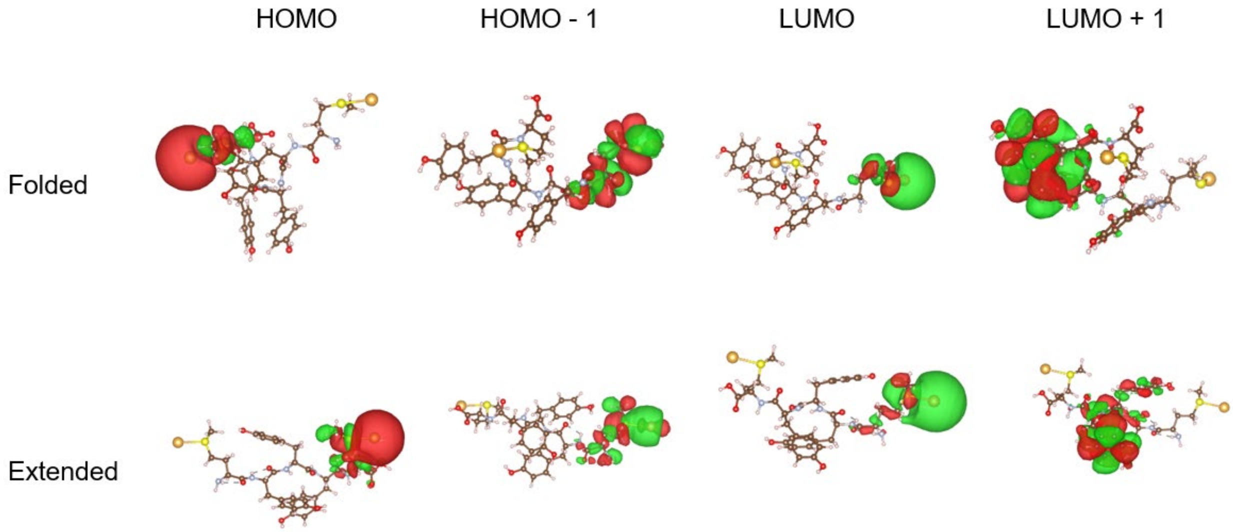

**Supplementary Figure 42**: Isosurface plots for HOMO, HOMO-1, LUMO and LUMO +1 orbital visualization for MYYYM

### S11. Tunneling pathway model: Bond counting for pathway determination

In this section, we explore the various electron transport pathways that are possible for MAAM sequence **(Supplementary Figure 36)**. If the charge transport is entirely through bond, then transport is required between atoms 1→16 in the figure below (16 atoms). For the folded structures that correspond to the high conductance state, an H-bond occurs between atoms 5* and 12*. In the case of electron transport through a hydrogen bond, transport can occur through 13 atoms [1→5 +5→5*+5*→12 (H-bond) + 12→16] as compared to 16 atoms.

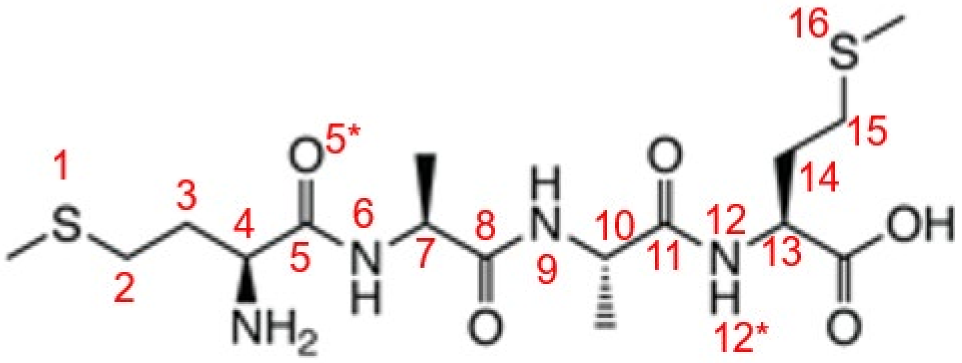

**Supplementary Figure 43:** Bond counting methods to understand tunneling pathway for MAAM sequence.

The decay associated with a covalent bond and through a H-bond[4] are given by the expressions below.

$$\epsilon_C = 0.6 \quad (1)$$

$$\epsilon_H = 0.36 \exp[-1.7(R - 2.8)] \quad (2)$$

The electron needs to cross 15 bonds for an entirely through bond mediated charge transport (1→16). This would indicate the decay to be $(0.6)^{15}=0.00047$. On the other hand, if the electron transport occurs via a combination of through bond and through H-bond ( 1→5 +5→5*+5*→12 (H-bond) + 12→16), the electron needs to travel 9 covalent bonds and one H-bond. This would indicate that the decay is $(0.6)^9\{0.36 \exp[-1.7(R - 2.8)\} = 0.0026$ where R is approximately 3 Angstrom (**Supplementary Figure 19**) for the MAAM sequence discussed above.

### S12. Control experiment: 1,16- hexadecanedithiol

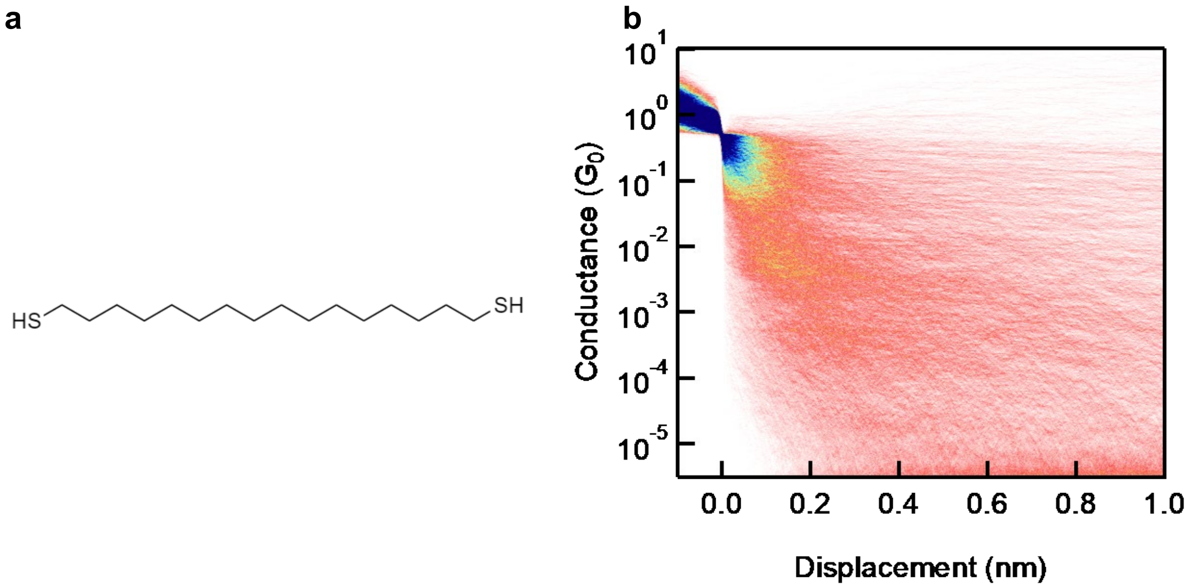

**Supplementary Figure 44**: Chemical structure and 2D conductance histogram for 1,16-hexadecanedithiol.

### S13. Convergence plots for MD simulations

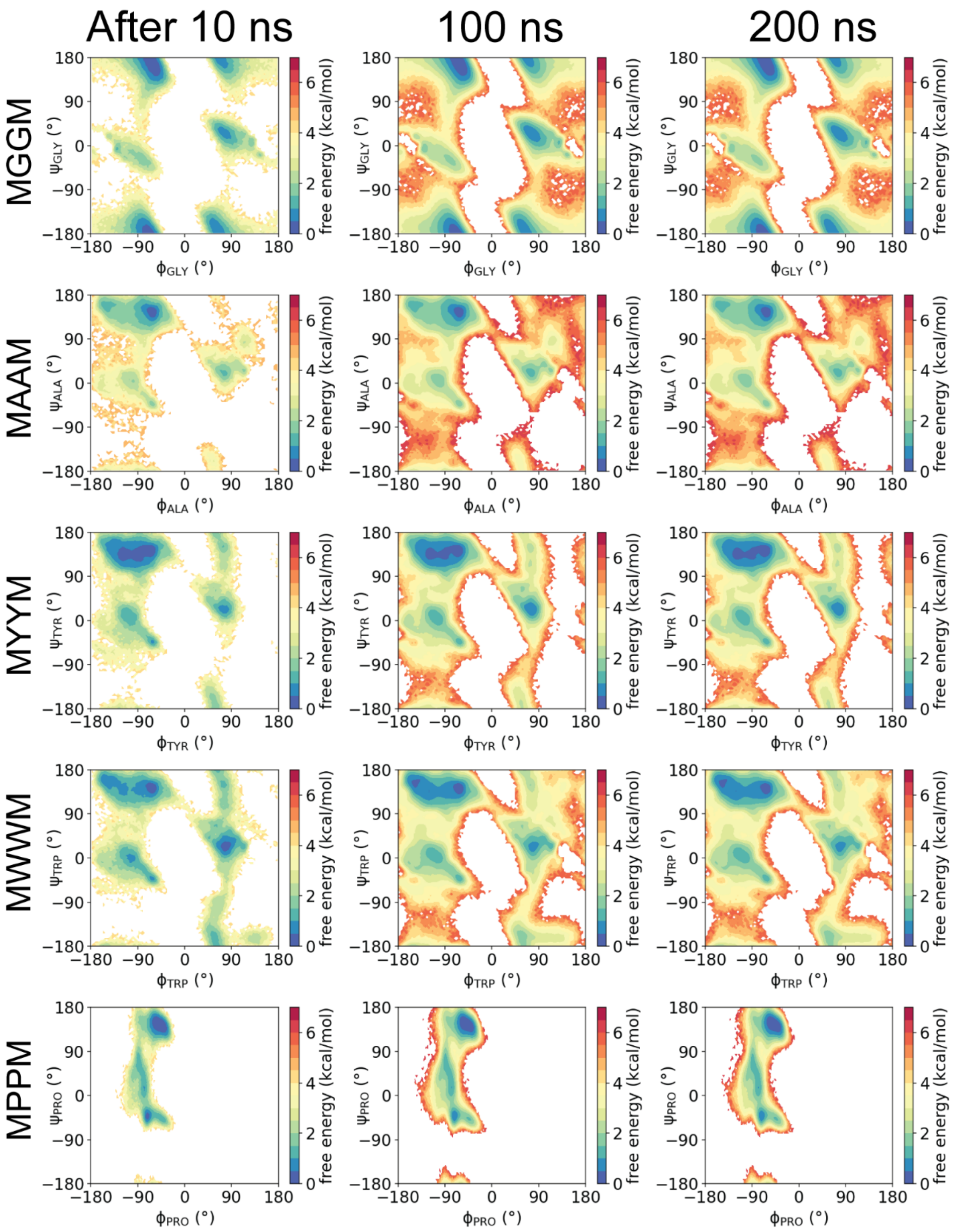

**Supplementary Figure 45**: Ramachandran plots for tetrapeptides. Identical plots are obtained after 100 ns and 200 ns, which indicates that the simulations are converged.

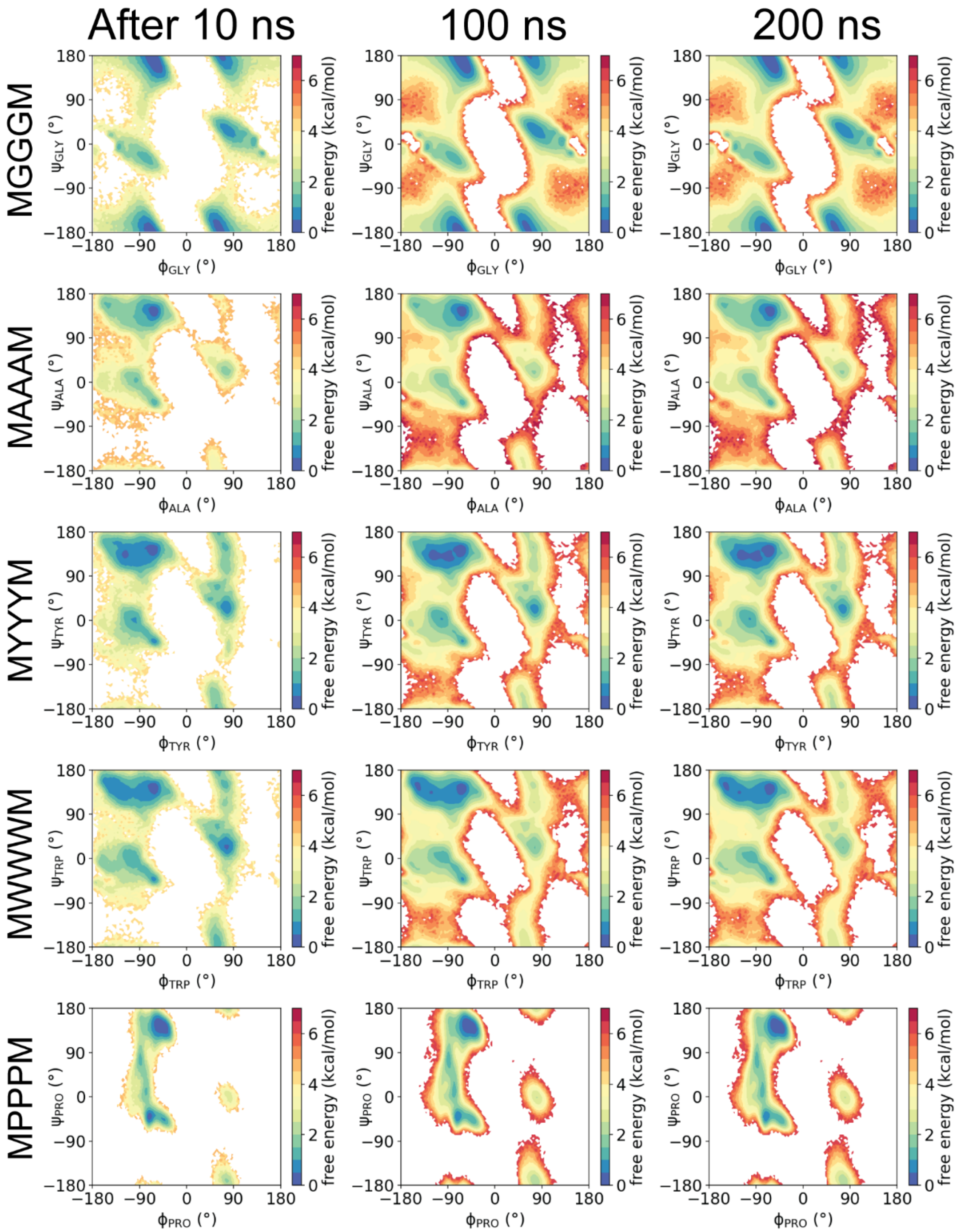

**Supplementary Figure 46**: Ramachandran plots for pentapeptides. Identical plots are obtained after 100 ns and 200 ns, which indicates that the simulations are converged.

### References

1. Stefani, D. *et al. Conformation-dependent charge transport through short peptides*. (2021).

2. Lin, L. *et al.* Spectral clustering to analyze the hidden events in single-molecule break junctions. *Journal of Physical Chemistry C* **125**, 3623–3630 (2021).

3. Rousseeuw, P. J. *Silhouettes: a graphical aid to the interpretation and validation of cluster analysis*. *Journal of Computational and Applied Mathematics* vol. 20 (1987).
